# A reciprocal interplay between 5-HT_2A_ and mGlu_5_ receptors underlies neuroplasticity

**DOI:** 10.1101/2025.07.23.666382

**Authors:** Tomas del Olmo, Mathilde Decourcelle, Martial Séveno, Joël Bockaert, Philippe Marin, Carine Bécamel

## Abstract

The serotonin (5-HT)_2A_ receptor is the primary target of numerous psychoactive drugs including serotonergic psychedelics, and mediates psychedelics-induced neuroplasticity, but the signaling mechanisms involved remain poorly characterized. Using quantitative phosphoproteomics, we show that the administration of the hallucinogenic 5-HT_2A_ receptor agonist 2,5-dimethoxy-4-iodoamphetamine (DOI) to mice promotes the phosphorylation of synaptic proteins belonging to a strongly interconnected protein network and comprising the metabotropic glutamate (mGlu)_5_ receptor and the scaffolding protein Shank3. Functional studies revealed that hallucinogenic and non-hallucinogenic 5-HT_2A_ receptor agonists promote synaptic targeting of mGlu_5_ receptor and its association with Shank3. Furthermore, they gate neuroplasticity in cortical neurons through a mechanism requiring mGlu_5_ receptor, protein kinase C and Shank3. Conversely, neuroplasticity elicited by mGlu_5_ receptor activation depends on 5-HT_2A_ receptor. Collectively, these findings demonstrate that neuroplasticity-promoting properties of psychedelics depend on a functional, reciprocal interplay between 5-HT_2A_ and mGlu_5_ receptors involving the synaptic scaffolding protein Shank3.

## Introduction

The serotonin (5-hydroxytryptamine, 5-HT)_2A_ (5-HT_2A_) receptor is a member of the Class A G protein-coupled Receptor (GPCR) family canonically coupled to G_q/11_ that has always raised a particular interest in view of its implication in several psychiatric conditions and the action of numerous psychoactive drugs, such as serotonergic psychedelics and antipsychotics. While the psycho-mimetic effects of psychedelics, such as lysergic acid diethylamide (LSD), mescaline and psilocybin, are mediated by the activation of 5-HT_2A_ receptors, they are not reproduced by other agonists exhibiting similar affinities for the receptor, such as ergotamine and lisuride. This paradox was partially solved by the demonstration that psychedelics induce a specific transcriptomic signature through the specific engagement of a Pertussis toxin-sensitive G_i/o_-Src signaling pathway ^1,2^ and their preferential activation of β2-arrestin-dependent signaling ^3,4^, compared with non-hallucinogenic agonists of 5-HT_2A_ receptors. Quantitative phosphoproteomics also revealed that hallucinogenic agonists of 5-HT_2A_ receptor, but not non-hallucinogenic agonists, promote receptor phosphorylation at Ser^280^, a process preventing its desensitization and internalization and thereby leading to a more sustained receptor activation^5^.

More recently, interest in the 5-HT_2A_ receptor has further increased with the discovery that serotonergic psychedelics induce rapid and long-lasting antidepressant effects even in patients resistant to conventional antidepressants, such as serotonin reuptake inhibitors ^6–8^. Psychedelics induce more prolonged antidepressant effects and less pronounced adverse effects than the NMDA receptor antagonist ketamine, another fast-acting antidepressant ^7^. A single dose of psychedelics also induces rapid long-lasting structural changes in neurons ^9,10^, that sustain their therapeutic effects without the need for chronic administration.

5-HT_2A_ receptors are mostly expressed in pyramidal neurons of the prefrontal cortex (PFC) ^11,12^ that is considered to be the primary site for the therapeutic action of novel fast-acting antidepressants. The PFC is involved in the top-down control of Dorsal Raphe Nuclei (DRN) serotonergic neurons that underlies the antidepressant effects induced by ketamine or deep brain stimulation in rodents ^13–15^. 5-HT_2A_ receptor stimulation by psychedelics gates synaptic plasticity at PFC glutamatergic synapses ^16–18^ and promotes dendritic growth of PFC pyramidal neurons ^19^, highlighting its key influence upon structural and functional neuroplasticity in this brain region. However, the molecular mechanisms by which psychedelics promote neuroplasticity in the PFC remain poorly characterized, an issue we addressed by comparing changes in the synaptic phosphoproteome induced by the synthetic hallucinogenic 5-HT_2A_ receptor agonist 2,5-Dimethoxy-4-iodoamphetamine (DOI) in the PFC of wild type (WT) and 5- HT_2A_ receptor knockout (*htr2A^-/-^*) mice. These studies allowed us to identify a network of synaptic proteins interacting with the metabotropic glutamate 5 (mGlu_5_) receptor, which itself exhibited enhanced phosphorylation upon DOI administration. These findings prompted further functional studies that revealed a reciprocal interplay between 5-HT_2A_ and mGlu_5_ receptors involving the Shank3 protein to trigger structural plasticity in primary cortical neurons.

## Materials & Methods

### Animals

Wild-type male or gestating C57BL/6J mice were purchased from Janvier Laboratories. Mice were housed under standardized conditions with a 12-h light/dark cycle, stable temperature (22 ± 1°C), controlled humidity (55 ± 10%), and free access to food and water. *Htr2A^−/−^* mice have a C57BL6/J background. *Shank3*^ΔC/ΔC^ were kindly provided by Dr Paul F Worley, Johns Hopkins University, and have a C57BL6/J background. Animal husbandry and experimental procedures were performed in compliance with the animal use and care guidelines of the University of Montpellier, the French Agriculture Ministry, and the European Council Directive (86/609/EEC).

### Antibodies

PSD95 (mouse, NeuroMab #75-028), mGlu_5_ receptor (rabbit, Cell Signaling #559205) and Shank3 (rabbit, Genetex #GTX133133) antibodies were diluted at 1:1,000 for Western blotting and at 1:/100 for immunohistochemistry. The MAP2 antibody (Abcam, Ab183830) was diluted at 1:100 in structural plasticity experiments. Rabbit Alexafluor488 (Invitrogen # A11034) and mouse Alexafluor546 (Invitrogen # A11030) secondary antibodies were diluted at 1:500. MGlu_5_ receptor (mouse, Millipore #Mabn540) and Shank3 (rabbit, Genetex #GTX133133) antibodies were diluted at 1:100 for the proximity ligation assay (PLA). The 5-HT_2A_ receptor (rabbit, Immunostar #24288) and β-actin HRP (Santa Cruz #sc-47778) antibodies were diluted at 1:1,000 and 1:4,000 for Western blotting, respectively.

### Drugs

For *in vivo* experiments, 2,5-Dimethoxy-4-iodoamphetamine (DOI; Sigma #D101) was injected intraperitoneally at the dose of 5 mg/kg. For experiments in primary cultures, the 5-HT_2A_ receptor was stimulated with DOI or lisuride maleate (Santa Cruz Biotechnologies) at the concentration of 10 μM. The mGlu_5_ receptor was inhibited with 10 μM of 2-Methyl-6- (phenylethynyl) pyridine (MPEP; Hellobio #HB0426) and stimulated with 50 μM of (S)-3,5- Dihydroxyphenylglycine (DHPG; Hellobio #HB0045). PKC was inhibited with 10 μM of GF109203X (Tocris #0741).

### Preparation of synapse-enriched fractions

Adult mice (postnatal day 60 to 90) were injected intraperitoneally either with 5 mg/kg of DOI or vehicle solution, 1 h before sacrifice. They were briefly anesthetized with isoflurane and sacrificed by cervical disruption. Brains were rapidly removed and placed into ice-cold dissection buffer containing 25 mM NaHCO_3_ (Sigma #S5761), 1.25 mM NaH_2_PO_4_ (Sigma #S8282), 2.5 mM KCl (Sigma #P3911), 0.5 mM CaCl_2_ (Sigma #C5080), 7 mM MgCl_2_ (Sigma #M2670), 25 mM glucose (Sigma #G7021), 110 mM choline chloride (Sigma #C1879), 11.6 mM ascorbic acid (Sigma #A4034), and 3.1 mM pyruvic acid (Sigma #P3256), and maintained under 5% CO_2_/95% O_2_. PFCs were dissected and frozen in liquid nitrogen. They were homogenized in an ice-cold buffer (0.32 M sucrose, 10 mM HEPES (Sigma #H3784), pH 7.4, and cocktails of protease (Roche, #11836170001) and phosphatase inhibitors (Roche, #04906837001) and cleared by centrifugation at 1,000 × g for 10 min to remove nuclei and large debris. Ten percent of the homogenates were collected to get whole protein extracts. The remaining material was centrifuged 1,000 x g once again. Supernatant was harvested and centrifuged at 12,000 x g for 20 min. Pellets were resuspended in a buffer containing 4 mM HEPES pH 7.4, 1 mM EDTA (Sigma #11836170001) and the protease and phosphatase inhibitor cocktails, and centrifuged again at 12,000 x g for 20 min. This step was repeated once. The last pellet containing the synapse-enriched fraction was then homogenized in a buffer containing 20 mM Tris-HCl pH 6, 1% Triton X-100 (Sigma #X100) and protease and phosphatase inhibitors. Protein concentration in whole protein extracts and synapse-enriched fractions was determined by the bicinchoninic acid method.

### Phosphoproteomic analysis

Label-free quantitative phosphoproteomics was performed on three biological replicates. Proteins from synapse-enriched fractions (80 µg) were digested on S-TrapTM mini spin columns (Protifi) according to the manufacturer’s protocol. Enzymatic digestion was performed by addition of 1 µg of trypsin (Gold, Promega, Madison USA) for 1 h at 47°C. Peptides were eluted with 50 mM of TEABC (Triethylammonium hydrogen carbonate buffer, Sigma #T7408) and 0.2% of formic acid. Phosphopeptides were subjected to sequential enrichment using metal oxide affinity chromatography (SMOAC) on Titanium dioxide (High- Select^TM^ TiO_2_ phosphopeptide enrichment kit, Thermo Fisher Scientific) and ferric nitrilotriacetate (High-Select^TM^ Fe-NTA phosphopeptide enrichment kit, Thermo Fisher Scientific) columns. Peptides from the non-enriched and enriched fractions were analysed by nanoflow liquid chromatography coupled to tandem mass spectrometry (nano-LC-MS/MS) using a Q-Exactive HF mass spectrometer coupled with an Ultimate 3000 RSLC system (Thermo Fischer Scientific). Desalting and pre-concentration of samples were performed online on a PepMap® precolumn (0.3 mm x 5 mm; Thermo Fisher Scientific). Peptides were introduced onto the column (Pepmap® 100 C18; 0.075 mm x 500 mm; Thermo Fisher Scientific) with buffer A (0.1% acid formic) and eluted with a 123-min gradient of 2-40% of buffer B (0.1% acid formic 80% CAN) at a flow rate of 300 nL/min. Spectra were acquired with Xcalibur software (v4.1 Thermo Fisher Scientific). MS/MS analyses were performed in data-dependent acquisition mode. Full scans (375 – 1500 m/z) were acquired in the Orbitrap mass analyzer with a 60,000 resolution at 200 m/z. For the full scans, 3 x 10^6^ ions were accumulated within a maximum injection time of 60 ms and detected in the Orbitrap analyzer. The twelve most intense ions with charge states > 2 were sequentially isolated to target value of 1 x 10^5^ with a maximum injection time of 100 ms and fragmented by HCD (Higher-energy collision dissociation) in the collision cell (normalized energy of 28%) and detected in the Orbitrap analyzer at 30,000 resolutions. MS/MS data analyses were performed using the Maxquant environment (v2.0.3.0) [1] ^20^and Andromeda for database search. All MS/MS spectra were matched against the UniProt Reference proteome (ID UP000000589, Mus musculus, UniProt release 2020_03) and 250 frequently observed contaminants as well as reversed sequences of all entries. Enzyme specificity was set to trypsin/P and the search included cysteine carbamidomethylation as fixed modification, and oxidation of methionine, acetylation and phosphorylation of Ser, Thr, Tyr residues (STY) as variable modifications. Up to two missed cleavages were allowed for protease digestion. FDR was set at 0.01 for peptides and proteins and the minimal peptide length at 7 residues. Relative protein and phosphorylated peptide quantification in synapse-enriched fractions was performed using the label-free quantification (LFQ) algorithm (https://maxquant.net/maxquant/).

### Bioinformatic analysis of phosphoproteomic data

Phosphopeptide intensities were normalized to the mean intensity of all proteins before enrichment for each condition. Results of LFQ for total proteins and phosphorylated peptides were analysed thanks to Prostar v1.34.5 software (https://www.prostar-proteomics.org/). Missing values were imputed through the det quantile algorithm. Each condition was compared one to one using the Student’s t-test, and the threshold fold change was fixed at 2. P-value was corrected through the Benjamin-Hochberg method, and the threshold p-value fixed at 0.03. Volcano plots were generated on Prostar v1.34.5. Phosphopeptide signals were analysed using Skyline v20.1 (https://skyline.ms/) and peak areas were exported, normalized on the corresponding protein intensity and analysed with Prostar (with the same settings as before) and Prism v10 software. Venn diagram was generated on Venny 2.1.1 (https://bioinfogp.cnb.csic.es/tools/venny/). GO biological process analysis and protein network generation were performed using the String plugin in Cytoscape v3.9.1 after Skyline validation. Kinase prediction was performed on KSEA software (https://casecpb.shinyapps.io/ksea/) with the phosphosite + Networkin database, with the score cutoff set at 2, p-value at 0.05 and substrate count at 5. The biological terms associated with regulated phosphosites were determined using the KEA2 software (https://www.maayanlab.net/KEA2/index.html) using the biological terms associated with phosphosites from literature mining.

### Western blotting

Proteins were resolved on 4–15% gradient gels (Bio-Rad) and transferred electrophoretic ally onto nitrocellulose membranes (Bio-Rad). Membranes were incubated in blocking buffer (Tris-HCl, 50 mM, pH 7.5; NaCl, 200 mM; Tween-20, 0.1%, and skimmed dried milk, 5%) for 1 h at room temperature and overnight with primary antibodies (see the “Antibodies” section) in blocking buffer. Then, membranes were washed and incubated with horseradish peroxidase-conjugated anti-rabbit (Cell Signaling #7074s) or anti-mouse (Cell Signaling #7076s) antibodies (1:10,000 in blocking buffer) for 1 h at room temperature. Immunoreactivity was detected with an enhanced chemiluminescence method (ECL detection reagent, GE Healthcare) using a ChemiDoc™ Touch Imaging System (Bio-Rad).

### Primary cultures of cortical neurons

Primary cultures of cortical neurons were prepared as previously described ^21^. Briefly, dissociated cells from the cerebral cortex of 17-day-old B6J mouse embryos were plated on glass coverslips placed on 24-well culture dishes (75,000 cells/coverslip) previously coated with poly-l-ornithine (mol. Wt. = 40,000; 15 μg/ml), and grown for 6 days for structural plasticity experiments or 14 days for colocalization and PLA experiments in Neurobasal A medium supplemented with B27 (2% v/v), 0.2 mM GlutaMAX, 0.2 mM glutamine and penicillin (100 u)/streptomycin (100 u).

### Immunohistochemistry and Sholl analysis

Cortical neurons were grown 14 days in vitro (DIV) and treated with either 5-HT_2A_ receptor agonists (DOI, 10 μM) or a mGlu_5_ receptor agonist (DHPG, 50 μM) for 15 min. Then, cells were placed on ice and washed once with cold PBS before to be fixed for 15 min at room temperature in paraformaldehyde (PFA, 4% in PBS). They were then washed three times for 5 min with PBS and permeabilized for 60 min at room temperature in PBS containing 5% goat serum and 0.3% Triton X-100. Cells were then incubated with primary antibodies (see the “Antibodies” section) overnight at 4°C in PBS containing 1% BSA and 0.3% Triton X-100 (antibody buffer). Cells were then washed three times for 5 min in PBS at room temperature and incubated for 1 h at room temperature with secondary antibodies (see the “Antibodies” section). They were incubated with DAPI (1 μg/ml) for 10 min, washed three times for 5 min in PBS at room temperature. Coverslips were mounted on glass slide in DAKO mounting media. For colocalization experiments, images were acquired on a Zeiss LSM980 confocal microscope equipped with a 63X objective. Single Z-sections were acquired, and settings were set in order to avoid pixel saturation and ensure proper colocalization. DAPI was used to unbiasedly select fields where images were acquired. Gain/offset settings were similarly adjusted between conditions. Linear modifications of fluorescence, cropping and resolution changes to 300 dpi were performed using Photoshop CC2017 for data presentation. Image colocalization was achieved with CellProfiler version 4.2.4 (https://cellprofiler.org). Pearson’s correlation was measured on original images using the measure correlation module. For structural plasticity experiments, DIV5 cortical neurons were treated with either MPEP (10 μM) or GF109203X (10 μM) for 15 min, in order to inhibits mGlu_5_ receptor or the protein kinase C (PKC), respectively. Cells were then exposed for 24 h ^22^ to either DOI (10 μM), lisuride (10 μM) or DHPG (50 μM). Cells were fixed and incubated with primary and secondary antibodies (see section “Antibodies”). Coverslips were mounted on glass slide in DAKO mounting media. Images were acquired with a 63X objective using an AxioImagerZ1 microscope equipped with Apotome (Zeiss). Branching number was counted manually and Sholl analysis was performed with Neuroanatomy plugin in ImageJ. All measurements were exported to Prism v10 for statistical analysis.

### Proximity Labelling Assay

DIV14 cortical neurons were treated with 5-HT_2A_ receptor agonists (DOI, 10 μM) for 15 min. Then, cells were placed on ice and washed once with cold PBS before to be fixed for 15 min at room temperature in paraformaldehyde (PFA, 4% in PBS). Proximity labelling was performed using the Duolink interaction kit (Sigma). Cells were washed on ice with 1X PBS and fixed for 15 min at room temperature with PFA 4% in PBS. Cells were blocked and permeabilized with the Duolink blocking solution for 60 min at room temperature. They were then incubated with primary antibody (see Antibodies section) overnight at 4°C. Probe incubation, ligation and amplification were performed according to the manufacturer’s instructions. Coverslips were mounted in PLA mounting media containing DAPI (1 μg/ml). Images were acquired with a 63X objective using the AxioImagerZ1 microscope equipped with Apotome (Zeiss). Fields were selected using DAPI labelling. Proximity Labelling Assay (PLA) dots were counted using the Cell profiler 4.2.4 primary object module and normalized to the number of nuclei. All measurements were exported to Prism for statistical analysis.

### Experimental design and statistics

Phosphoproteomic experiments were performed on three independent biological replicates. One to one Student’s t-tests with Benjamin-Hochberg correction were performed using the Prostar v1.34.5 software. For mGlu_5_ receptor, GRIN2B, Shank3 and LRRC7 phosphorylation intensities. Data from Skyline were analysed by two-way ANOVA followed by Fisher’s LSD tests using Prism v10 software. For colocalization and PLA experiments, two-way ANOVA followed by Fisher’s LSD test was performed on data acquired on at least 35 images containing a minimum of two cells from four biological replicates per condition. Statistics of Sholl analysis and branching counting were performed on a minimum of 30 neurons per condition originating from 4 independent biological replicates. To compare Sholl areas under the curve, Brown-Forsythe and Welch ANOVA followed by Dunnett’s test was performed. To compare branching numbers Gaussian distribution was first verified (Shapiro-Wilk normality test). Then, data were analysed either by two-way ANOVA followed by Fisher’s LSD test or ordinary one-way ANOVA followed by Tukey’s test in groups with normal distribution and by Kruskal-Wallis followed by Dunn’s test in groups with non-parametric distribution. The threshold for statistical significance was set at 0.05.

## Data availability

The raw mass spectrometry data have been deposited to the ProteomeXchange Consortium via the PRIDE partner repository with the dataset identifier PXD057142.

## Results

### DOI promotes the phosphorylation of synaptic proteins connected to mGlu_5_ receptor in prefrontal cortex

To characterize changes in the synaptic phosphoproteome elicited upon 5-HT_2A_ receptor stimulation by DOI, synapse-enriched fractions from PFC of WT and *htr2A^-/-^* mice, treated with DOI (5mg/kg) or vehicle (Veh) for one hour, were digested with trypsin and the resulting peptide samples were enriched in phosphorylated peptides by sequential metal oxide affinity chromatography (SMOAC). Both enriched (synaptic phosphoproteome) and non-enriched (total synaptic proteome) fractions were analysed by liquid chromatography coupled to tandem mass spectrometry (LC-MS/MS) (**Fig. S1A**). Proteins and phosphorylated peptides were quantified using a label-free approach and the Maxquant software. We identified a total of 2,149 proteins in synapse-enriched fractions. Most of them are related to the glutamatergic synapse, as assessed by GO cellular component analysis (**Fig. S1B**). Their relative quantification in Veh- and DOI-treated WT or *htr2A^-/-^* mice showed that a 1-h treatment with DOI does not alter the synaptic proteome in both strains (**Fig. 1A, 1B**). Analysis of fractions enriched in phosphorylated peptides identified a total of 980 phosphorylated residues on 351 unique proteins (**Fig. 1C, 1D**). These include 374 phosphorylated residues on 189 proteins whose phosphorylation level was regulated by DOI administration to WT mice (**Fig. 1C, 1D, Fig. S1C and Table S1)**. Only 18 of them (including 14 common to both strains) were also regulated by DOI in *htr2A^-/-^* mice (**Fig. S1D**), indicating that DOI-elicited changes in the synaptic phosphoproteome are mainly mediated by 5-HT_2A_ receptor activation. To further validate the observed quantitative changes in synaptic phosphoproteome elicited by DOI, we quantified the peak current intensities of phosphosites showing missing values in Maxquant analysis, using the Skyline software. This resulted in a list of 92 phosphorylated sites on 42 proteins exhibiting significant changes in their phosphorylation level in DOI-treated mice (**Fig. S2)**. Analysis of the biological processes associated with this set of proteins using the String enrichment plugin in Cytoscape showed enrichment in ‘Transsynaptic signaling’, ‘Cell-cell signaling’, ‘Nervous system development’, ‘Chemical synaptic transmission’, ‘Dendrites development’ and ‘Synapse organization’ terms, suggesting a role in synapse transmission and neuroplasticity. Analysis of previously described associations between the 42 proteins phosphorylated upon DOI treatment using String showed that 31 of them are part of a strongly interconnected complex, comprising the mGlu_5_ receptor (**Fig S3A and S3B**), the NMDA receptor subunits GRIN1 and GRIN2B (**Fig S3C and S3D**) and the synaptic scaffolding protein SH3 And Multiple Ankyrin Repeat Domains 3 (Shank3) (**Fig. 1E, S3E and S3F**). The mGlu_5_ receptor that interacts with 9 proteins, namely Solute Carrier Family 1 Member 2 (SLC1A2), GRIN1, GRIN2B, Shank3, Leucine Rich Repeat Containing 7 (LRRC7), Gamma-Aminobutyric Acid Type B Receptor Subunit 2 (GABBR2), Synapsin-1 and Synapsin-2, Erythrocyte membrane protein band 4.1 (EBP4L1) and Catenin Delta 2 (CTNND2) (**Fig. 1E**) seems to be a central hub in this network. In order to assess the functions directly related to the phosphorylation set induced by DOI treatment, we analysed the GO terms associated with each phosphorylated site using KEA2 software to reveal enriched pathways. This highlighted a link between 5-HT_2A_ receptor activity and Ca^2+^, PKC signalling and glutamate-related functions, consistent with the canonical receptor coupling to Gq, and the well-described crosstalk between 5-HT_2A_ and NMDA receptor signalling and modulation of glutamatergic transmission by 5-HT_2A_ receptors ^16,23,24^ (**Fig. S2B**).

**Figure 1:**
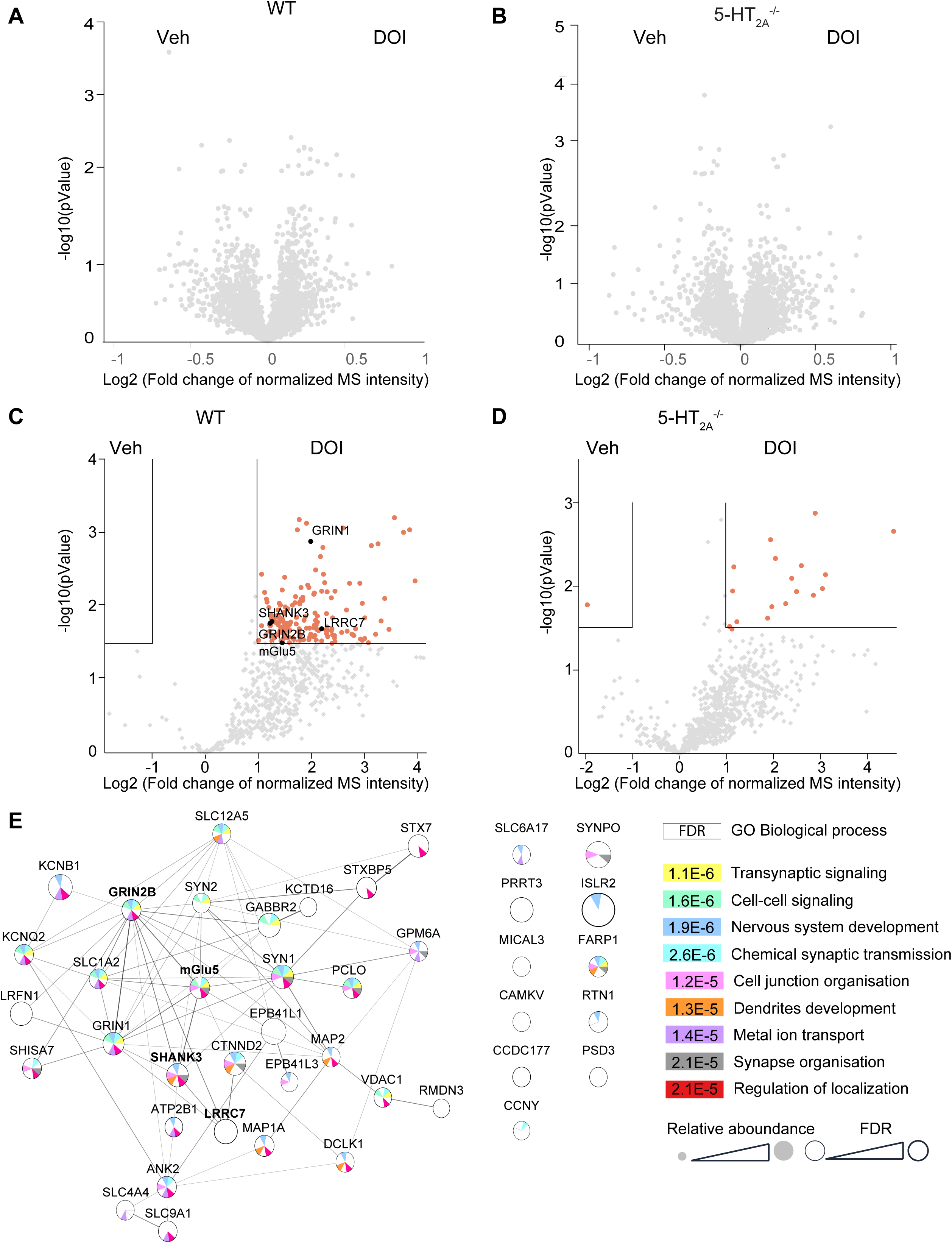
*S*ynaptic phosphoproteome changes induced upon 5-HT_2A_ receptor stimulation. **A and B**. Volcano plots comparing protein abundances in PFC synaptic-enriched fraction from WT (A) and *htr2a*^-/-^ (**B**) mice treated with vehicle or DOI (5mg/kg). N=3 biological replicates. **C and D**. Volcano plots comparing phosphopeptide abundances in PFC synaptic-enriched fraction from WT (A) and *htr2A*^-/-^ (B) mice treated with vehicle or DOI (5mg/kg). N=3 biological replicates. Peptides exhibiting significant changes in their phosphorylation state in DOI vs. vehicle conditions (Fold Change > 2 and p-value < 0.03) are in red. **E.** Synaptic protein network whose phosphorylation upon DOI administration to WT mice was confirmed by skyline (fold change > 2 and p-value < 0.05). Protein network was generated on Cytoscape with String database. Lines between proteins represents known interactions. Size of circle represent the relative abundance of the phosphorylation and thickness represents its significance. Circular diagram colors represent the GO biological processes associated to each protein.

### 5-HT2A receptor regulates mGlu_5_ receptor and Shank3 localization at the synapse

The connection of 5-HT_2A_ receptor to mGlu_5_ receptor signalosome prompted further investigation of the functional relationship between both receptors. Given the presence of proteins known to control the synaptic targeting of membrane receptors (Shank3, SHISA7, LRRC7, FARP1, PCLO, SYN1, SYNPO, STXBP5, or ANK2) among the proteins phosphorylated in response to DOI administration (**Fig1. E**), we investigated the effect of 5-HT_2A_ receptor deletion on the synaptic localization of mGlu_5_ receptor. We first compared mGlu_5_ receptor abundance in synaptosome-enriched fractions from WT and *htr2A^-/-^*mice by quantitative mass spectrometry. The mGlu_5_ receptor MS intensity signal was significantly diminished in *htr2A^-/-^* mice, compared to WT mice (**Fig. S4A**). These results were confirmed by Western blotting that also indicated that mGlu_5_ receptor level in total PFC lysates was not affected by the deletion of 5-HT_2A_ receptor (**Fig. S4B-D**). Likewise, Shank3 level was significantly decreased in the synaptic compartment in *htr2A^-/-^* mice, whereas it was similar in total PFC lysates from WT and *htr2A^-/-^* mice (**Fig. 3C and D**). In contrast, PSD95 abundance in PFC synaptosomes and PFC lysates was not significantly different in WT and *htr2A^-/-^* mice. Altogether, these results suggest that the expression of 5-HT_2A_ receptor controls the synaptic localization of both mGlu_5_ receptor and Shank3 without altering the general integrity of synapses.

### 5-HT2A receptor activation enhances mGlu_5_ receptor and Shank3 localization at the synapse

We next investigated whether 5-HT_2A_ receptor stimulation promotes mGlu_5_ receptor and Shank3 recruitment to the synapse. Primary cultures of cortical neurons from WT and *htr2A^-/-^*mice were treated with either vehicle or DOI (10 μM) ^22^ for 15 min (**Fig. 2A**) and the colocalization of mGlu_5_ receptor or Shank3 with PSD95 was assessed by immunohistochemistry and confocal microscopy. Both mGlu_5_-PSD95 and Shank3-PSD95 co- localization were reduced in *htr2A^-/-^* cultures (Pearson’s correlation score 0.173 ± 0.016 and 0.114 ± 0.010) compared to WT cultures (Pearson’s correlation score 0.263 ± 0.017 and 0.208 ± 0.014, p=0.0012 and p<0.0001, respectively), confirming that the absence of 5-HT_2A_ receptor affects the synaptic localization of mGlu_5_ receptor and Shank3 (**Fig. 2A and B)**. Exposure of neurons to DOI increased mGlu_5_ receptor and Shank3 colocalization with PSD95 in WT cultures (Pearson’s correlation score 0.358 ± 0.018 and 0.279 ± 0.018, p=0.0006 and p=0.0004, respectively) but not in *htr2A^-/-^* cultures (Pearson’s correlation score 0.216 ± 0.024 and 0.144 ± 0.011, p=0.117 and p=0.099 respectively, **Fig. 2A and B)**. Exposure of WT neurons to DOI also increased the interaction between mGlu_5_ receptor and Shank3, as shown by the increase in the number of fluorescent dots (6.256 ± 0.571 vs. 13.176 ± 1.071, in Vehicle and DOI-treated neurons, respectively, p<0.0001) in a proximity ligation assay (**Fig. 2E and F**) but not in *htr2A^-/-^*cultures (4.017 ± 0.479 vs 1.622 vs 0.241 in Vehicle and DOI-treated neurons, respectively, p=0.077). Collectively, these results demonstrate that 5-HT_2A_ receptor stimulation promotes the synaptic localization of Shank3 and mGlu_5_ receptor and enhances their physical proximity in primary cortical neurons.

**Figure 2:**
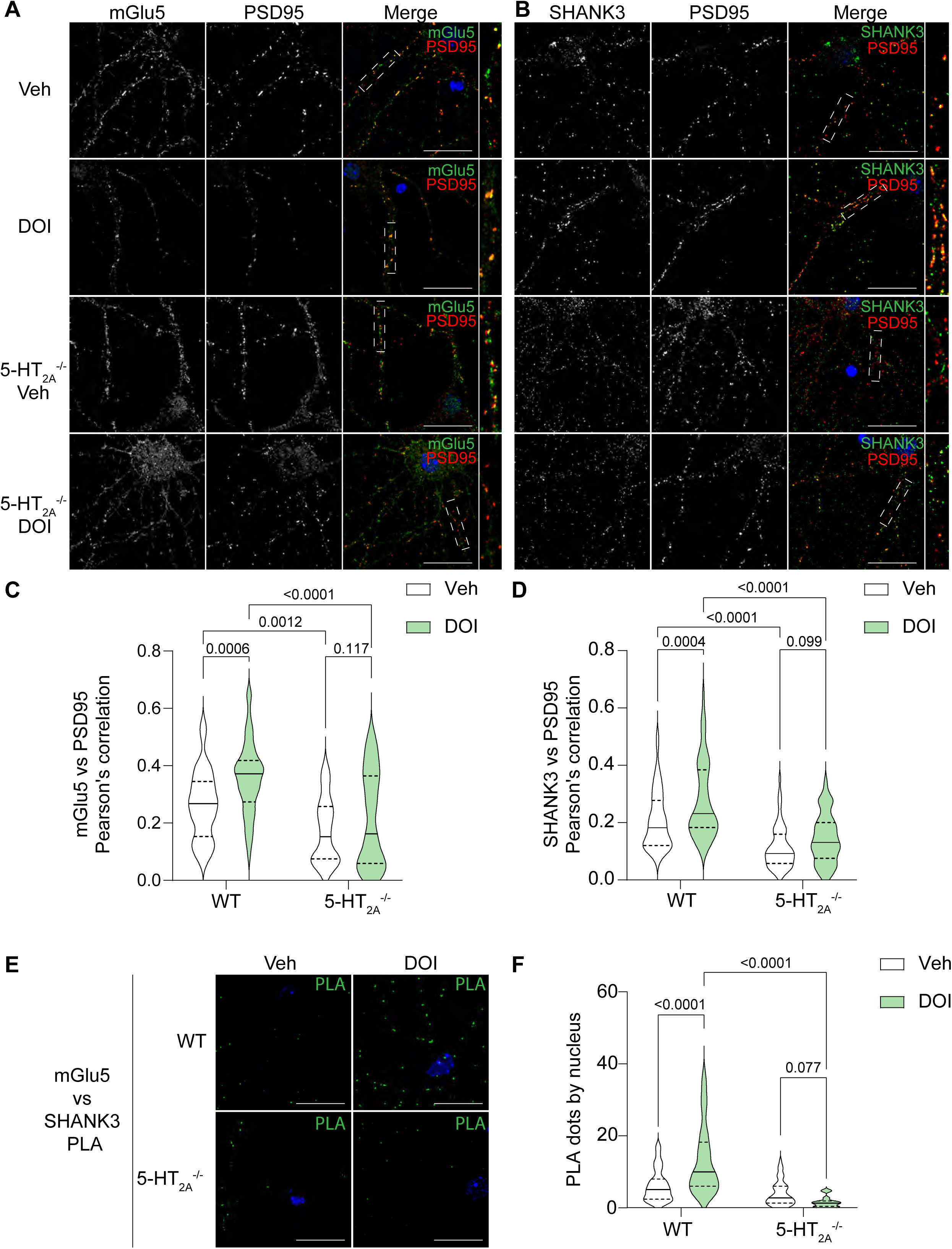
5-HT_2A_ receptor stimulation regulates mGlu_5_ receptor interaction with Shank3 interaction and their synaptic localization. **A and B.** Immunostaining of mGlu_5_ receptor (green, **A**) or Shank3 (green, **B**) and PSD95 (red) on primary cortical neurons (DIV14) from WT and *htr2A*^-/-^ mice treated for 15 min with vehicle or DOI (10 μM). Single-channel images are depicted and boxed region is magnified. Scale bars, 20 μm. **C and D.** Pearson’s correlation coefficient between PSD95 and mGlu_5_ receptor (**C**) or Shank3 (**D**) immunostainings, n > 45 images from 4 independent experiments. The Violin boxes represent the distribution, full line the mean and dashed line the quartiles of Pearson’s correlation coefficients, respectively. White violons correspond to vehicle condition in WT and *htr2A*^-/-^ mice and green violins to DOI-treated (10 μM) condition in both strains. Two-way ANOVA followed by Fisher’s LSD test was performed. The p-value is reported for each comparison. **E.** PLA signals (green) showing the interaction between mGlu_5_ receptor and Shank3 in primary cortical neurons from WT and *htr2A*^-/-^ mice treated with vehicle or DOI (10 μM). Scale bars, 20 μm. **F.** Violin box distribution of PLA dot numbers per nucleus in neurons from WT and *htr2A*^-/-^ mice treated for 15 min with vehicle (white) or DOI (10 μM, green), n > 35 images from 4 independent experiments. Two-way ANOVA followed by Fisher’s LSD test was performed. The p-value is reported for each comparison.

### Structural plasticity induced by 5-HT_2A_ receptor stimulation depends on mGlu_5_ receptor in primary cortical neurons

It was recently shown that serotonergic psychedelics, such as psilocybin or LSD, promote neuroplasticity, including dendritogenesis ^10,22,25–29^. Previous studies also demonstrated that both mGlu_5_ receptor activation ^30–32^ and Shank3 overexpression ^33^ induce neuroplasticity. First, we confirmed that 5-HT_2A_ and mGlu_5_ receptor stimulation promote dendritic growth by treating primary cortical neurons for 24 h with either DOI or the group I mGlu receptor agonist (S)-3,5-Dihydroxyphenylglycine (DHPG, 50 μM) and analysing the resulting changes in their morphological features. Sholl analysis ^34^ showed that exposing cortical cultures to DOI or DHPG increases dendritic arbor complexity of neurons, as assessed by the area under the curves in the Sholl plots and the number of dendrite branches (**Fig. 3A and Fig. 4C)**. DOI-induced increase in dendritic arbor complexity was not observed in primary cortical neurons from *htr2A^-/-^* mice (**Fig. S5A**) as well as in neurons from WT mice pre-treated with 2-Methyl-6- (phenylethynyl) pyridine (MPEP, 10 μM, 15 min, **Fig. 3A**), a mGlu_5_ receptor antagonist, indicating a functional crosstalk between both receptors to promote neuroplasticity. Interestingly, exposure to lisuride (10 μM, 24 h), a non-hallucinogenic 5-HT_2A_ receptor agonist that induces rapid and long-lasting antidepressant effects in a preclinical model of depression ^35^ (**Fig 3B**), also increased dendritic arbor complexity of cortical neurons from WT mice (**Fig. 3B**), but not *htr2A^-/-^* mice (**Fig. S5B**). Again, pre-treating WT cultures with MPEP abolished lisuride-induced neuroplasticity (**Fig. 3B**).

**Figure 3:**
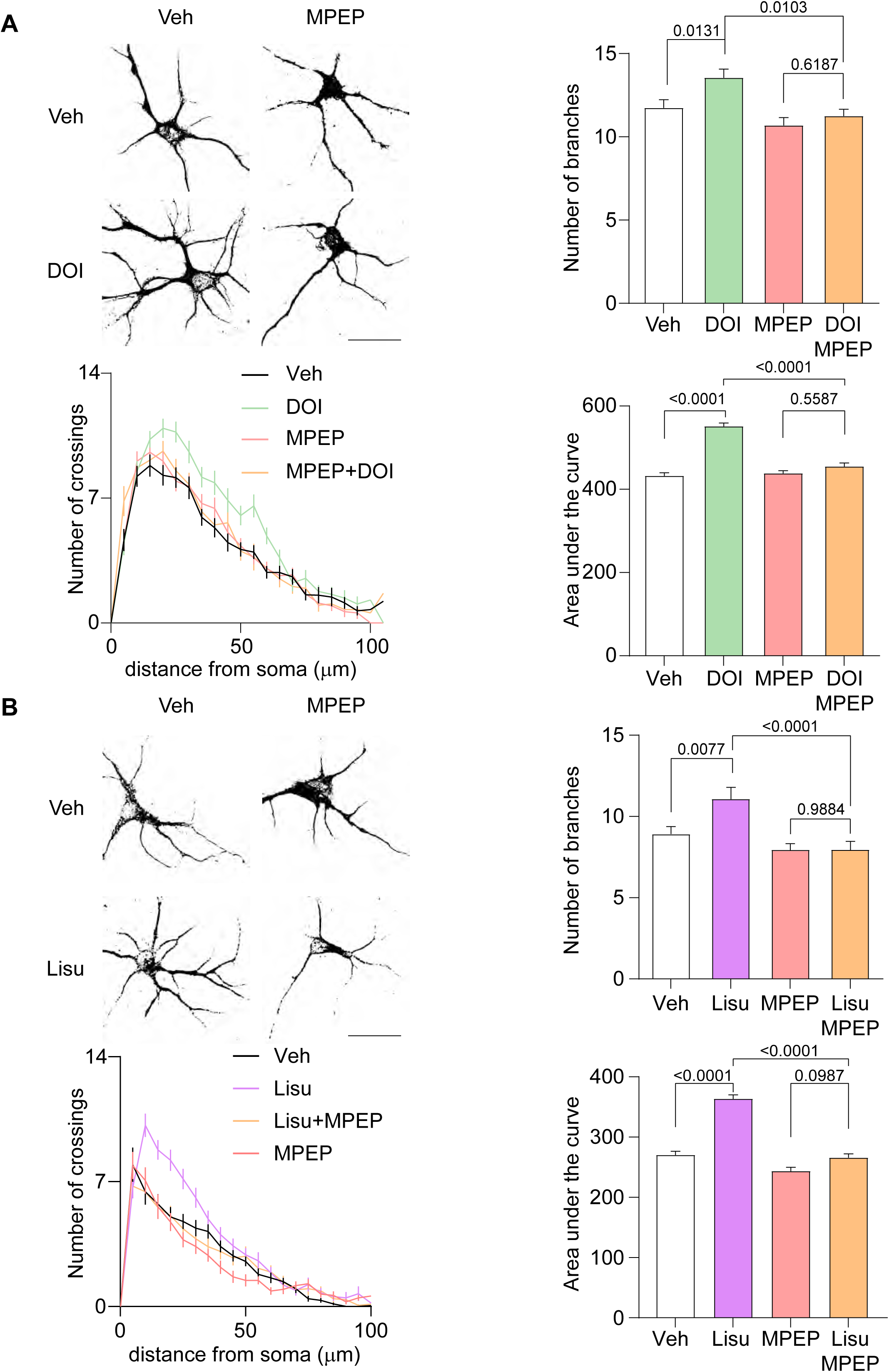
5-HT_2A_ receptor-mediated structural neuroplasticity depends on mGlu_5_ receptor in primary cortical neurons. **A.** Dendritogenesis induced in primary cortical neurons by 5-HT_2A_ receptor stimulation with DOI (10 μM) for 24 h and its inhibition by the mGlu_5_ receptor antagonist MPEP (10 μM). The upper left panel shows representative images of cortical neurons. The upper right panel shows a histogram representing number of branches per neuron in each condition, n > 60 neurons from 4 independent experiments. Kruskal-Wallis followed by Dunn’s test were performed and p-values are reported on the graph. The lower left panel illustrates the Sholl analysis of the number of dendrites crossing concentric circles from the soma every 5 μm, n > 30 neurons per condition. The lower right panel shows a histogram representing the area under the curve in the corresponding experiment. Brown-Forsythe and Welch ANOVA followed by the Dunnett’s test was performed and p-values are reported on the graph. **B.** Dendritogenesis induced in primary cortical neurons by 5-HT_2A_ receptor stimulation with lisuride (10 μM) for 24 h and its inhibition by MPEP (10 μM). The upper left panel shows representative images of cortical neurons. The upper right panel shows a histogram representing number of branches per neuron in each condition, n > 35 neurons from 4 independent experiments. The Kruskal-Wallis followed by Dunn’s test was performed and p-value reported on graph. The lower left panel illustrates the Sholl analysis of the number of dendrites crossing concentric circles from the soma every 5 μm, n > 30 neurons per condition. The lower right panel shows a histogram representing the area under the curve in the corresponding experiment. Brown-Forsythe and Welch ANOVA followed by the Dunnett’s test was performed and p-values are reported on the graph.

**Figure 4:**
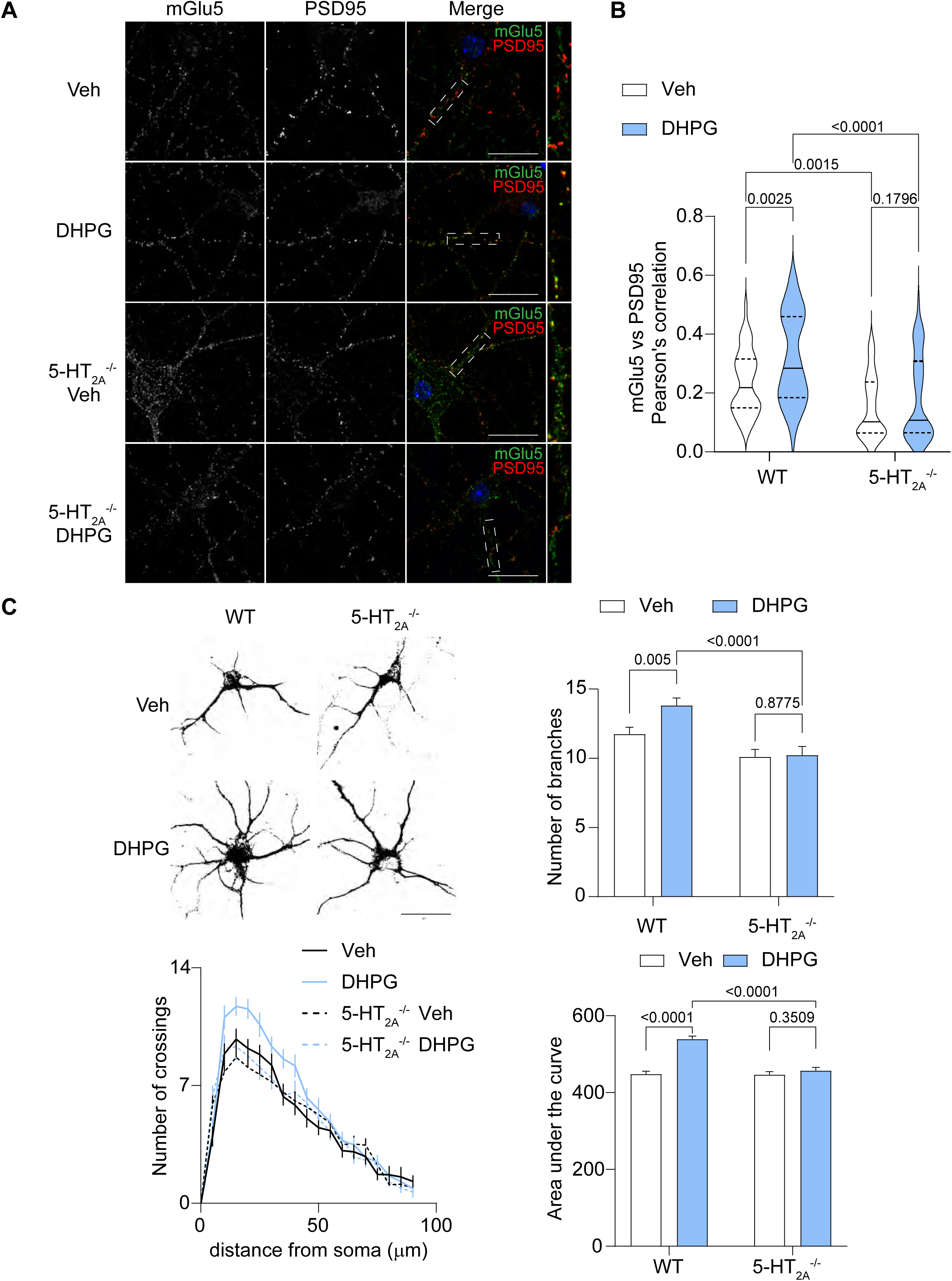
DHPG-induced structural neuroplasticity depends on 5-HT_2A_ receptor in primary cortical neurons. **A.** Immunostaining of mGlu_5_ receptor (green) and PSD95 (red) on primary cortical neurons from WT and *htr2A*^-/-^ mice treated for 15 min with vehicle or DHPG (50 μM). Single-channel images are depicted and boxed regions are magnified. Scale bars, 20 μm. **B**. Pearson’s correlation coefficient between PSD95 and mGlu_5_ receptor immunostainings, n > 45 images from 4 independent experiments. The violins represent Pearson’s correlation coefficient distribution, the full line represents the mean and dashed line the quartiles. White violons correspond to neurons from WT or *htr2A*^-/-^ mice exposed to vehicle, and green violins neurons from both strains exposed to DHPG (50 μM). Two-way ANOVA followed by Fisher’s LSD test was performed. The p-value is reported for each comparison. **C.** Dendritogenesis induced by mGlu_5_ receptor stimulation with DHPG (50 μM) for 24 h is inhibited by the deletion of the 5-HT_2A_ receptor. The upper left panel shows representative images of cortical neurons. The upper right panel shows a histogram representing the number of branches per neuron in each condition, n > 60 neurons from 4 independent experiments. Two-way ANOVA followed by Fisher’s LSD test was performed. The p-values are reported for each comparison. The lower left panel illustrates the Sholl analysis of the number of dendrites crossing concentric circles from the soma every 5 μm, n > 33 neurons per condition. The lower right panel shows a histogram representing area under the curve in the corresponding experiment. Brown-Forsythe and Welch ANOVA followed by Dunnett’s test was performed and p-values are reported on the graph.

### Structural plasticity elicited by mGlu_5_ receptor activation depends on 5-HT_2A_ receptor

Taken together, these results demonstrate that 5-HT_2A_ receptor stimulation by both hallucinogenic and non-hallucinogenic agonists promote structural plasticity in cortical neurons through a mechanism involving mGlu_5_ receptor activation. Previous studies have shown that mGlu_5_ receptors are mainly confined in perisynaptic domains ^36^ and that their activation increases receptor mobility ^37^. Consistent with these findings, a 15 min exposure of primary neurons from WT mice to DHPG increases mGlu_5_ receptor recruitment to the synapse in primary cortical neurons (**Fig.4A**), as assessed by colocalization with PSD95 (Pearson’s correlation score 0.237 ± 0.016 vs. 0.319 ± 0.022, in vehicle and DHPG-treated neurons, respectively, p<0.0025). DHPG-induced synaptic insertion of mGlu_5_ receptor was not seen in cultures from *htr2A^-/-^* mice (Pearson’s correlation score 0.150 ± 0.015 vs. 0.186 ± 0.021, in vehicle and DHPG-treated neurons, respectively, p=0.1796). Likewise, DHPG exposure did not increase dendritic arbor complexity (**Fig.4C**) in primary cortical neurons from *htr2A^-/-^* mice (**Fig. 4C**), demonstrating that neuroplasticity elicited by mGlu_5_ receptor activation depends on 5- HT_2A_ receptor. Collectively, these results suggest that a reciprocal interplay between both receptors gates structural plasticity in primary cortical neurons.

### Shank3 is essential for 5-HT_2A_ and mGlu_5_ receptor-dependent neuroplasticity

Given that 5-HT_2A_ receptor stimulation promotes Shank3 phosphorylation, synaptic localization and association with mGlu_5_ receptor, and the role of Shank3 in mGlu_5_ receptor signalling at the synapse ^38^, we next examined the role of Shank3 in 5-HT_2A_ and mGlu_5_ receptor- dependent neuroplasticity. Neither DOI nor DHPG treatment increased dendritic arbor complexity in primary cortical neurons from *Shank3^ΔC/ΔC^* mice that carry a deletion of exon 21, resulting in the truncation of the C-terminal tail of Shank3, the loss of its functionality ^39^ and the disruption of mGlu_5_ receptor signalosome ^40^ (**Fig. 3C and Fig. 4C**). This suggests that Shank3 might act as a functional molecular link between 5-HT_2A_ and mGlu_5_ receptors that contributes to neuroplasticity gating upon stimulation of each receptor.

### PKC activation mediates 5-HT_2A_ and mGlu_5_ receptor-gated structural plasticity in cortical neurons

We submitted our phosphoproteome dataset to the Kinase Enrichment Analysis (KEA) software to predict kinases that may phosphorylate the identified residues based on sequence analysis and knowledge of kinase substrate specificities (**Fig. 5A**). These analyses identified several protein kinases C (PKC) isoforms (PKCγ, PKCα and PKCγ2) as the best candidates contributing to DOI-induced phosphorylation set (**Fig. 5A**). These results are consistent with the canonical coupling of 5-HT_2A_ receptor (and mGlu_5_ receptor) to Gq/11 proteins and our phosphosite-associated terms analysis showing that PKC was among the most enriched terms (**Fig. S2B**). They are also reminiscent of a previous study indicating that PKC directly phosphorylates Ser^839^ on mGlu_5_ receptor, a site found to be upregulated upon 5-HT_2A_ receptor stimulation by DOI in the present study (Fig. S3A, S3B) and essential for Ca^2+^ oscillations produced by mGlu_5_ receptor activation ^41^. To investigate the role of PKC in structural plasticity gating by 5-HT_2A_ and mGlu_5_ receptor stimulation, cortical neurons were treated with either DHPG (50 μM) or DOI (10 μM) in the presence or absence of GF109203X (10 μM), a potent and selective PKC inhibitor. Sholl analysis showed that inhibition of PKC abolished the increase in dendritic arbor complexity elicited by DOI and DHPG treatments (**Fig 5B and 5C**), indicating that both 5-HT_2A_ and mGlu_5_ receptors gate structural plasticity in primary cortical neurons through the activation of PKC.

**Figure 5:**
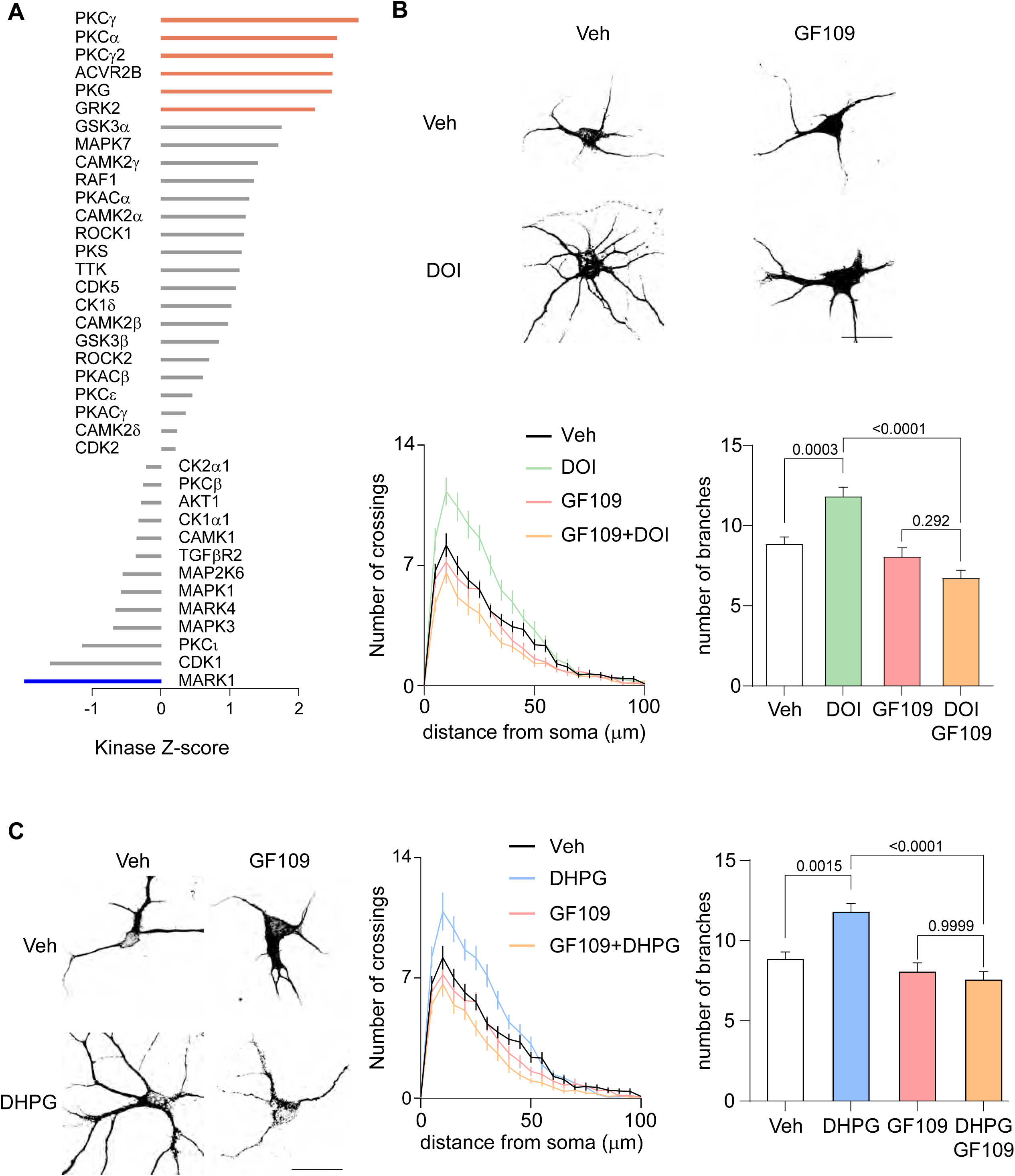
mGlu_5_ and 5-HT_2A_ receptor-dependent structural neuroplasticity require PKC activity. **A.** Normalized kinase prediction Z score, generated with KSEA software, for the phosphosites regulated by DOI treatment administration to WT mice. The kinases over-represented are in red and the under-represented ones are in blue. **B.** Dendritogenesis induced by 5-HT_2A_ receptor stimulation with DOI (10 μM) for 24 h and its inhibition by GF109203X (GF109, 10 μM), a ‘selective’ inhibitor of PKC activity. The upper left panel shows representative images of cortical neurons. The upper right panel shows a histogram representing number of branches per neuron in each condition, bars and error bars correspond to the mean ± SEM, n > 60 neurons from 4 independent experiments. One-way ANOVA followed by Tukey’s test was performed and p-value reported on graph. The lower left panel illustrates Sholl analysis representing the number of dendrites crossing concentric circles from the soma every 5 μm, error bars correspond to SEM, n > 30 neurons per condition. The lower right panel shows a histogram representing area under the curve of Sholl analysis. Brown-Forsythe and Welch ANOVA followed by Dunnett’s test were performed and p-value reported on graph. **C.** Dendritogenesis evoked by mGlu_5_ receptor stimulation with DHPG (50 μM) for 24 h and its inhibition by GF109203X (10 μm). The upper left panel shows representative images of cortical neurons. The upper right panel shows a histogram representing number of branches per neuron in each condition, bars and error bars correspond to the mean ± SEM, n > 35 neurons from 4 independent experiments. Kruskal-Wallis followed by Dunn’s test was performed and p-value reported on graph. The lower left panel illustrates Sholl analysis representing the number of dendrites crossing concentric circles from the soma every 5 μm, error bars correspond to SEM, n > 25 neurons per condition. The lower left panel shows a histogram representing area under the curve of Sholl analysis. Brown-Forsythe and Welch ANOVA followed by Dunnett’s test was performed and p-value reported on graph.

**Figure 6:**
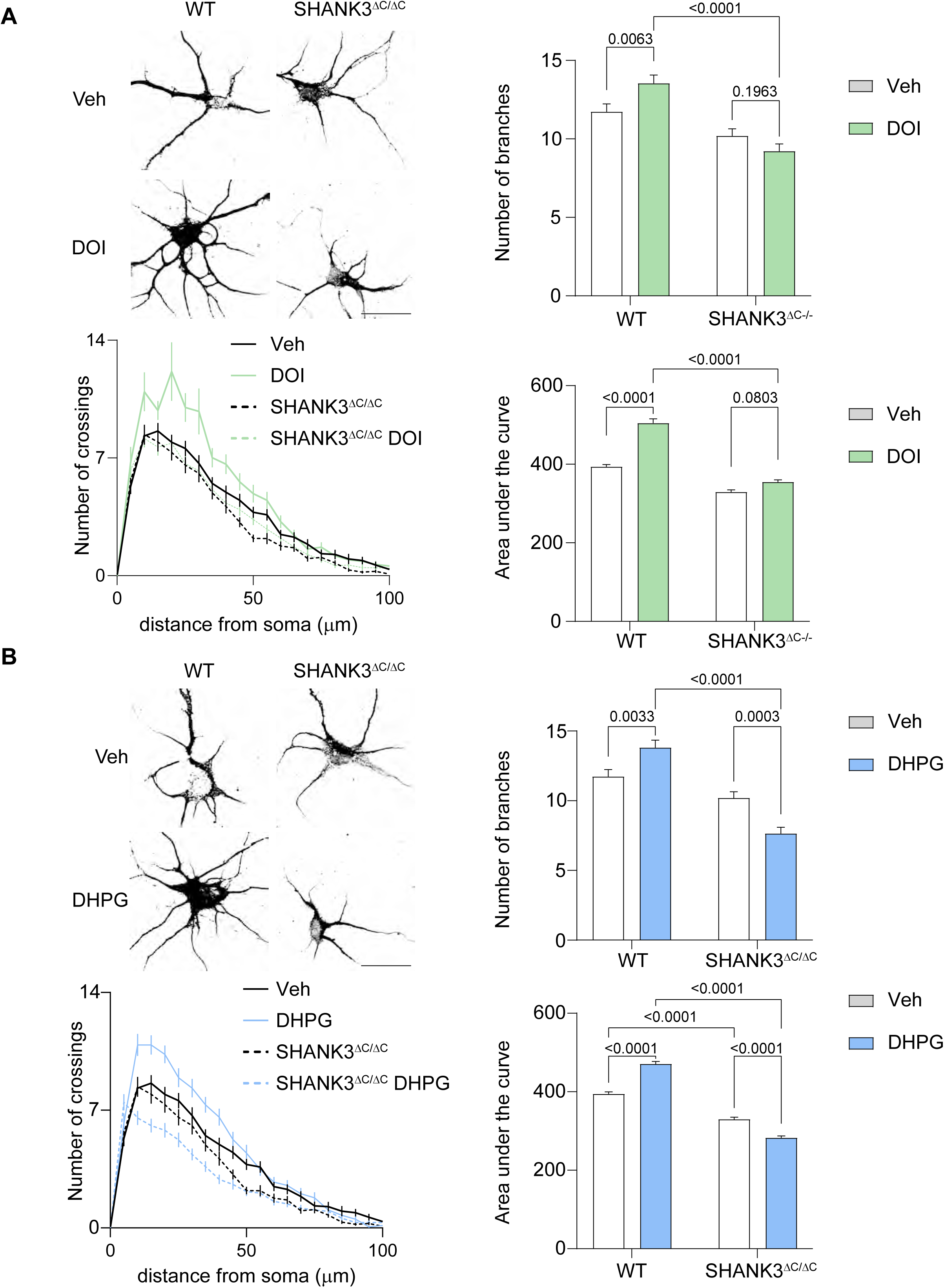
Shank3 mediates 5-HT_2A_ and mGlu_5_ receptor-dependent structural neuroplasticity. **A.** Dendritogenesis induced by 5-HT_2A_ receptor stimulation with DOI (10 μM) for 24 h and its inhibition by the deletion of Shank3 exon21. The upper left panel shows representative images of cortical neurons. The upper right panel shows a histogram representing number of branches per neuron in each condition, bars and error bars correspond to the mean ± SEM, n > 60 neurons from 4 independent experiments. Two-way ANOVA followed by Fischer’s LSD test was performed and p-value reported on graph. The lower left panel illustrates Sholl analysis representing the number of dendrites crossing concentric circles from the soma every 5 μm, error bars correspond to SEM, n > 30 neurons per condition. The lower right panel shows a histogram representing area under the curve of Sholl analysis. Brown-Forsythe and Welch ANOVA followed by Dunnett’s test were performed and p-value reported on graph. **B.** Dendritogenesis induced by mGlu_5_ receptor stimulation with DHPG (50 μM) for 24 h and its inhibition by the deletion of Shank3 exon21. The upper left panel shows representative images of cortical neurons. The upper right panel shows a histogram representing number of branches per neuron in each condition, bars and error bars correspond to the mean ± SEM, n > 35 neurons from 4 independent experiments. Kruskal-Wallis followed by Dunn’s test was performed and p-value reported on graph. The lower left panel illustrates Sholl analysis representing the number of dendrites crossing concentric circles from the soma every 5 μm error bars correspond to SEM, n > 25 neurons per condition. The lower right panel shows a histogram representing area under the curve of Sholl analysis. Brown-Forsythe and Welch ANOVA followed by Dunnett’s test was performed and p-values reported on graph.

## Discussion

The present study characterized for the first time changes in the synaptic phosphoproteome induced by 5-HT_2A_ receptor stimulation in PFC thanks to an unbiased quantitative phosphoproteomic approach. While we identified a total of 2,149 proteins and 980 phosphorylated residues on 351 unique proteins in synapse-enriched fractions, DOI administration induced changes in the phosphorylation state of 374 residues (i.e., 38% of the identified phosphorylated peptides) on 189 synaptic proteins that represent more than 50% of the identified synaptic phosphorylated proteins. Thus, stimulation of prefrontal 5-HT_2A_ receptors induces substantial changes in synaptic phosphoproteome that might contribute to 5-HT_2A_ receptor-dependent modulation of synaptic transmission and plasticity in this brain region ^16,18^. The low density of 5-HT_2A_ receptor (and any 5-HT receptor subtype) precludes their detection by mass spectrometry in brain tissue without applying a specific enrichment procedure such as immunoprecipitation ^42^. This explains why we did not detect in synapse- enriched fractions 5-HT_2A_ receptor phosphorylation on Ser^280^, a phosphorylation site previously identified in HEK293 cells overexpressing the receptor and further validated in prefrontal cortex using a phosphosite-specific antibody ^5^.

Phosphorylated protein analysis suggests that synaptic 5-HT_2A_ receptor targets are strongly embedded with glutamatergic signaling. For instance, 5-HT_2A_ receptor promotes the phosphorylation of the NMDA receptor subunits GRIN1 (on Ser^889^) and GRIN2B (on Ser^1303^). This is consistent with previous studies showing that 5-HT_2A_ receptor activation enhances NMDA transmission and gates the induction of temporal-dependent plasticity mediated by NMDA receptors through the phosphorylation of GRIN2B on Ser^1303^ ^16^. Furthermore, 5-HT_2A_ receptor stimulation promotes the phosphorylation of mGlu_5_ receptor (on Ser^839^) and several proteins that are part of the mGlu_5_ signalosome, such as GRIN1 and GRIN2B, LRRC7 (on Ser^1206^) and Shank3 (on Ser^781^). This suggests a functional connection between 5-HT_2A_ and mGlu_5_ receptors that are known to exhibit a predominant post-synaptic localization ^43,44^. Whether 5- HT_2A_ and mGlu_5_ receptors form heteromers and whether heteromerization is important for 5- HT_2A_ receptor-operated mGlu_5_ receptor phosphorylation remain to be established.

In fact, recent studies have shown that the mGlu_5_ receptor bulk is mostly located at the perisynaptic compartment ^36^ and is highly mobile ^45^, Furthermore, mGlu_5_ stabilization at the post-synapse is essential to promote its activity. Here, we show that stimulation of the 5-HT_2A_ receptor promotes mGlu_5_ receptor and Shank3 synaptic targeting, as assessed by the increase in their colocalization with PSD95, thereby favoring their physical association. These effects might reflect the increase in mGlu_5_ receptor phosphorylation on Ser^839^, a residue located in a domain essential for its synaptic localization, and the increased phosphorylation of Shank3 on Ser^781^, a residue previously shown to be crucial for the synaptic enrichment of the rat ortholog of Shank3 ^46^.

Analysis of 5-HT_2A_ receptor-associated synaptic phosphoproteome changes also identified numerous proteins involved in neurite growth. These include MAP1A (Microtubule-Associated Protein 1A) ^47^, MAP2 Microtubule-Associated Protein 2) ^48^, DCLK1 (Doublecortin Like Kinase 1) ^49^, CTNND2 (Catenin Delta 2) ^50^, FARP1 (FERM, ARH/RhoGEF And Pleckstrin Domain Protein 1) ^51^, mGlu_5_ receptor ^30–32^ and Shank3^52^. Consistent with these findings, previous studies have demonstrated that 5-HT_2A_ receptor agonists, including psychedelics, promote neurite growth ^22,53^ through a mechanism that remains to be elucidated. Here, we confirm these results and show that DOI as well as lisuride induce neurite growth in primary cortical neurons from WT but not *htr2A^-/-^* mice. These findings unequivocally demonstrate that structural plasticity induced by both drugs depends on 5-HT_2A_ receptor activation. This contrasts with previous studies indicating that both LSD and psilocybin promote structural plasticity through their direct binding to TrkB receptor in a 5-HT_2A_ receptor- independent manner, thereby acting as allosteric modulators of BDNF signaling ^29^. The discrepancy between these studies and the present work could be due to the different molecules tested that belong to different chemical families. Finally, the ability of the non-hallucinogenic 5-HT_2A_ receptor agonist lisuride to promote neurite growth, as previously demonstrated with other molecules devoid of hallucinogenic properties in mice ^54,55^, demonstrate that the hallucinogenic properties of 5- HT_2A_ receptor agonists are not necessary to their effects on structural plasticity.

The present study provides new insight into the mechanisms by which 5-HT_2A_ and mGlu_5_ receptor promote structural plasticity in cortical neurons. We show that neurite growth elicited by 5-HT_2A_ receptor agonists is prevented by mGlu_5_ receptor pharmacological inhibition while 5-HT_2A_ receptor deletion inhibits neurite growth induced by the mGlu_5_ receptor agonist DHPG, demonstrating a reciprocal functional crosstalk between both receptors to promote structural plasticity. Whether this crosstalk involves the formation of 5-HT_2A_/mGlu_5_ receptor heteromers or occurs intracellularly at the level of signaling pathways remains to be established. A recent study suggested that structural plasticity induced by psychedelics belonging to the tryptamine family depends on intracellular 5-HT_2A_ receptor ^56^. Other studies have also shown that intracellular mGlu_5_ receptors can activate ERK1/2, Elk-1 and Arc pathways ^57,58^ and mediate synaptic plasticity in neurons ^59^. This suggests that a crosstalk between intracellular 5-HT_2A_ and mGlu_5_ receptors might promote structural plasticity. Here, we used the mGlu_5_ receptor agonist DHPG which would only activate cell surface receptors ^59,60^, suggesting that structural plasticity induced by 5-HT_2A_ and mGlu_5_ receptor agonists rather results from a functional crosstalk between both receptors located on the cell surface, even though one cannot rule out internalization and signaling in early endosomes of DHPG-bound mGlu_5_ receptors. Importantly, a functional interaction between 5-HT2A and mGlu5 receptors has previously been shown to play a role in regulating locomotor behavior in mice^61^.

There is growing evidence that the mGlu_5_ receptor is an important molecular player in the pathophysiology of depression and the therapeutic effects of antidepressants ^62^. On one hand, mGlu_5_ receptor inhibition by antagonists or NAMs induce antidepressant-like effects in several depression paradigms in rodents. On the other hand, conventional antidepressant molecules, electroconvulsive therapy, sleep deprivation as well as ketamine ^63,64^ induce expression of the immediate early gene *Homer1a*, an effect leading to agonist-independent activation of mGlu_5_ receptor ^65^. These contrasting results might reflect the differential effects of receptors expressed on glutamatergic vs. GABAergic neurons: the specific knockout of mGlu_5_ receptor in glutamatergic neurons where 5-HT_2A_ receptors are predominantly expressed, causes depression-like behavior in the mouse, whereas its deletion in GABAergic neurons induces antidepressant-like effects ^66^. Furthermore, studies have shown that mGlu_5_ receptors might be essential for the antidepressant effects of sleep deprivation and ketamine ^67^. Given the role of mGlu_5_ receptors in structural plasticity induced by serotonergic psychedelics, their involvement in the antidepressant effects of these drugs certainly warrants further exploration.

## Acknowledgments

The authors thank Dr. Julie Perroy (IGF) for providing the Shank3^ΔC/ΔC^ mice. This study was supported by grants from CNRS, INSERM, University of Montpellier. Imaging was performed using the facilities of magnetic resonance imaging platform, MS experiments using facilities of the Functional Proteomics Platform (Biocampus Montpellier). T. Del Olmo received post- doctoral fellowships from the LabEx Epigenmed and from Fondation pour la Recherche Médicale.

## Author Contributions

T.D.O. designed, performed and analysed all the experiments. M.D and M.S. performed LC-MS/MS experiments. C.B. and P.M. designed the experiments and supervised the study; T.D.O, C.B., P.M. and J.B. wrote the manuscript.

## Supplemental information

Figures S1–S5

