## Supplementary material for "A reciprocal interplay between 5-HT_2A_ and mGlu_5_ receptors underlies neuroplasticity": supp data

### Supplemental figures

**Figure S1: Synaptosome enrichment and mass spectrometry analysis**

**A.** Workflow of the phosphoproteomic approach. **B.** GO cellular component analysis of proteins identified in PFC synaptosomes from WT and *htr2a* $^{-/-}$ mice performed on Cytoscape with the String database. **C.** Venn diagram comparing protein phosphorylated upon DOI treatment in PFC synaptosomes from WT and *htr2A*$^{-/-}$ mice.

**Figure S2: Synaptic phosphoproteome induced upon 5-HT$_{2A}$ receptor stimulation with DOI, is interconnected with glutamatergic signaling.**

**A.** Volcano plots comparing phosphopeptide peak intensities (assessed using Skyline) in synaptosomes preparation from vehicle (left) and DOI (right) treated mice, N=3 biological replicates. Phosphopeptides exhibiting significant changes in their phosphorylation state upon DOI treatment (Fold Change > 2 and p-value < 0.05) are in red. **B.** Biological terms analysis associated with the verified phosphosite dataset. The analysis was performed using the KEA2 software.

**Figure S3: Validation of mGlu$_5$ receptor, Shank3, LRRC7 and GRIN2B phosphorylation**

**A, C, E and G**. MS signal intensity plots generated on Skyline, represent phosphopeptide (corresponding to mGlu$_5$ receptor phosphorylated on Ser$^{839}$ (A), LRRC7 on Ser$^{1206}$ (C), Shank3 on Ser$^{781}$ (E) and GRIN2B on Ser$^{1303}$ (G), respectively) peak intensities in synaptosomes from WT and *htr2a*$^{-/-}$ mice treated with vehicle or DOI. **B, D, F and H**. Histograms representing the mean of phosphosite peak intensities from A, C, E and G, respectively. N = 3 biological replicates. Two-way ANOVA followed by Fisher's LSD test was performed, p-values are reported on the graph.

**Figure S4: 5-HT$_{2A}$ receptor promotes mGlu$_5$ receptor and Shank3 synaptic localization**

**A**. Histogram representing normalized mGlu$_5$ peak intensities between WT and *htr2a*$^{-/-}$ synaptosomes. An Unpaired t-test was performed and p-values were reported on graph.

**B**. Immunoblot analysis of 5-$HT_{2A}$ receptor, $mGlu_5$ receptor, Shank3, PSD95 and Actin expression in PFC lysates and synaptosomes from WT and *htr2A*$^{-/-}$ mice. **C**. Quantification of $mGlu_5$ receptor, Shank3 and PSD95 expression (normalized to Actin expression) in PFC synaptosomes from WT (grey histogram) and *htr2A*$^{-/-}$ (white histogram) mice. Two-way ANOVA followed by a Fisher's LSD test was performed and p-values reported on graph.

**D**. Quantification of $mGlu_5$ receptor, Shank3 and PSD95 expression (normalized to Actin expression) in PFC lysates from WT (grey histogram) and *htr2A*$^{-/-}$ (white histogram) mice. Bars and error bars correspond to the mean ± SEM. Each point represents individual values. Two-way ANOVA followed by Fischer's LSD test was performed and the p-values are reported on the graph.

**Figure S5: DOI and lisuride-induced dendritogenesis in cortical neurons is mediated by the 5-$HT_{2A}$ receptor**

**A**. Dendritogenesis induced by DOI (10 μM) treatment for 24 h is inhibited by the deletion of 5-$HT_{2A}$ receptor. The upper left panel shows representative images of cortical neurons. The upper right panel shows a histogram representing number of branches per neuron in each condition, bars and error bars correspond to the mean ± SEM, n > 35 neurons from 4 independent experiments. The lower left panel illustrates Sholl analysis representing the number of dendrites crossing concentric circles from the soma every 5 μm, error bars correspond to SEM. The lower right panel shows a histogram representing area under the curve of Sholl analysis. Two-way ANOVA followed by Fisher's LSD test was performed, p-value is reported for each comparison. **B**. Dendritogenesis induced by lisuride (10 μM) treatment for 24 h is inhibited by the deletion of 5-$HT_{2A}$ receptor. The upper left panel shows representative images of cortical neurons. The upper right panel shows a histogram representing number of branches per neuron in each condition, bars and error bars correspond to the mean ± SEM, n > 35 neurons from 4 independent experiments. The lower left panel illustrates Sholl analysis representing the number of dendrites crossing concentric circles from the soma every 5 μm, error bars correspond to SEM. The lower right panel shows a histogram representing area under the curve of Sholl analysis. Two-way ANOVA followed by Fisher's LSD test was performed, p-value is reported for each comparison.

**Table S1: List and statistical analysis of maxquant phosphopeptide MS peak intensities**

**Column A**: Unique phosphopeptide I.D generated on Prostar software. **Column B**: Protein Uniprot accession. **Column C**: Phosphorylation site. **Column D**: Protein name. **Column E**: Gene name. **Column F**: Phosphorylated peptide sequence. **Column G**: Phosphosite position probabilities. **Column H, I, J**: MS peak intensities of phosphopeptide issued from PFC synaptic enriched fraction of WT mice injected I.P with Vehicle solution. **Column K, L, M**: MS peak intensities of phosphopeptides purified from PFC synaptic-enriched fraction of WT mice injected with DOI (5mg/kg) solution. **Column N, O, P**: MS peak intensities of phosphopeptides purified from PFC synaptic-enriched fraction of htr2A-/- mice injected with Vehicle solution. **Column Q, R, S**: MS peak intensities of phosphopeptides issued from PFC synaptic-enriched fraction of htr2A-/- mice injected with DOI (5mg/kg) solution **Column T to AE**: Phosphopeptide MS peak normalized on total protein MS intensities for each condition, respectively. **Column AF**: Log2 of the fold change of MS normalized phosphopeptide peak intensities between WT mice injected with either Veh or DOI (5mg/kg) solution. **Column AG**: p-Value obtained from Student t-test analysis of difference between WT mice injected with either Veh or DOI (5mg/kg) solution. **Column AH**: Log2 of the fold change of MS normalized phosphopeptides peak intensities between htr2A-/- mice injected with either Veh or DOI (5mg/kg) solution. **Column AI**: p-Value obtained from Student t-test analysis of difference between htr2A-/- mice injected with either Veh or DOI (5mg/kg) solution.

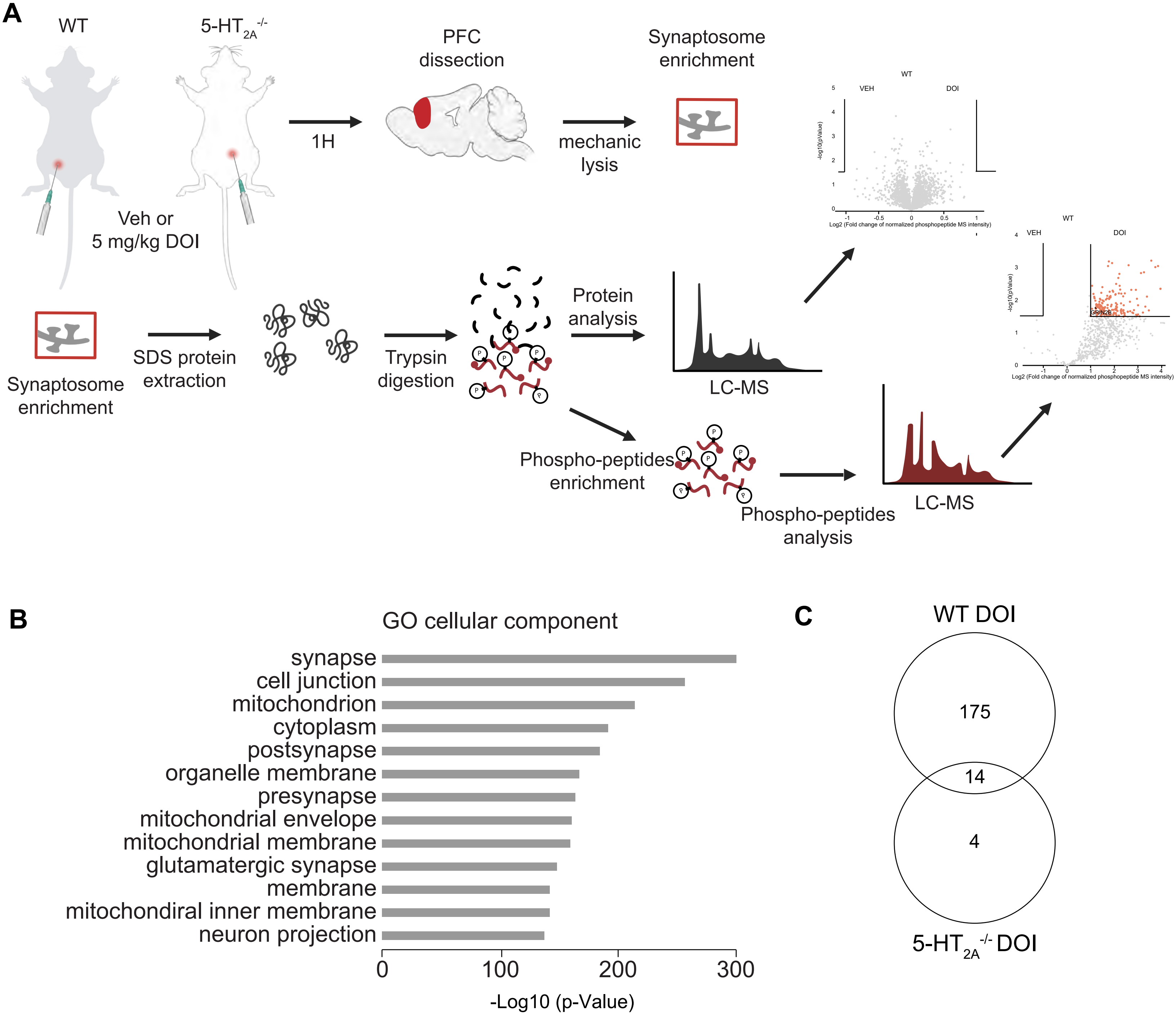
A
WT
5-HT2A -/-
Veh or
5 mg/kg DOI
1H
PFC
dissection
mechanic
lysis
Synaptosome
enrichment
Synaptosome
enrichment
SDS protein
extraction
Trypsin
digestion
Protein
analysis
LC-MS
Phospho-peptides
enrichment
Phospho-peptides
analysis
LC-MS
WT
VEH
DOI
-log10(pValue)
Log2 (Fold change of normalized phosphopeptide MS intensity)
B
GO cellular component
synapse
cell junction
mitochondrion
cytoplasm
postsynapse
organelle membrane
presynapse
mitochondrial envelope
mitochondrial membrane
glutamatergic synapse
membrane
mitochondiral inner membrane
neuron projection
0
100
200
300
-Log10 (p-Value)
C
WT DOI
175
14
4
5-HT2A -/- DOI

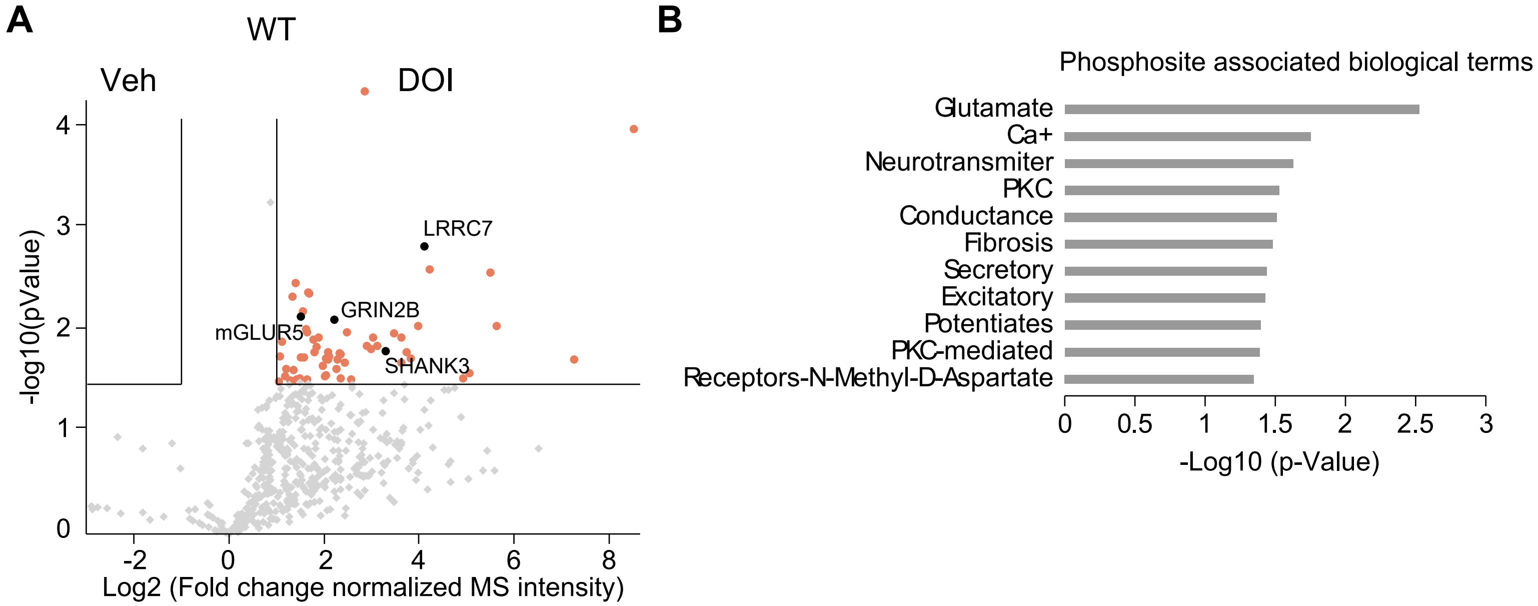
A
WT
Veh
DOI
LRRC7
GRIN2B
mGLUR5
SHANK3
-log10(pValue)
Log2 (Fold change normalized MS intensity)
B
Phosphosite associated biological terms
Glutamate
Ca+
Neurotransmiter
PKC
Conductance
Fibrosis
Secretory
Excitatory
Potentiates
PKC-mediated
Receptors-N-Methyl-D-Aspartate
-Log10 (p-Value)

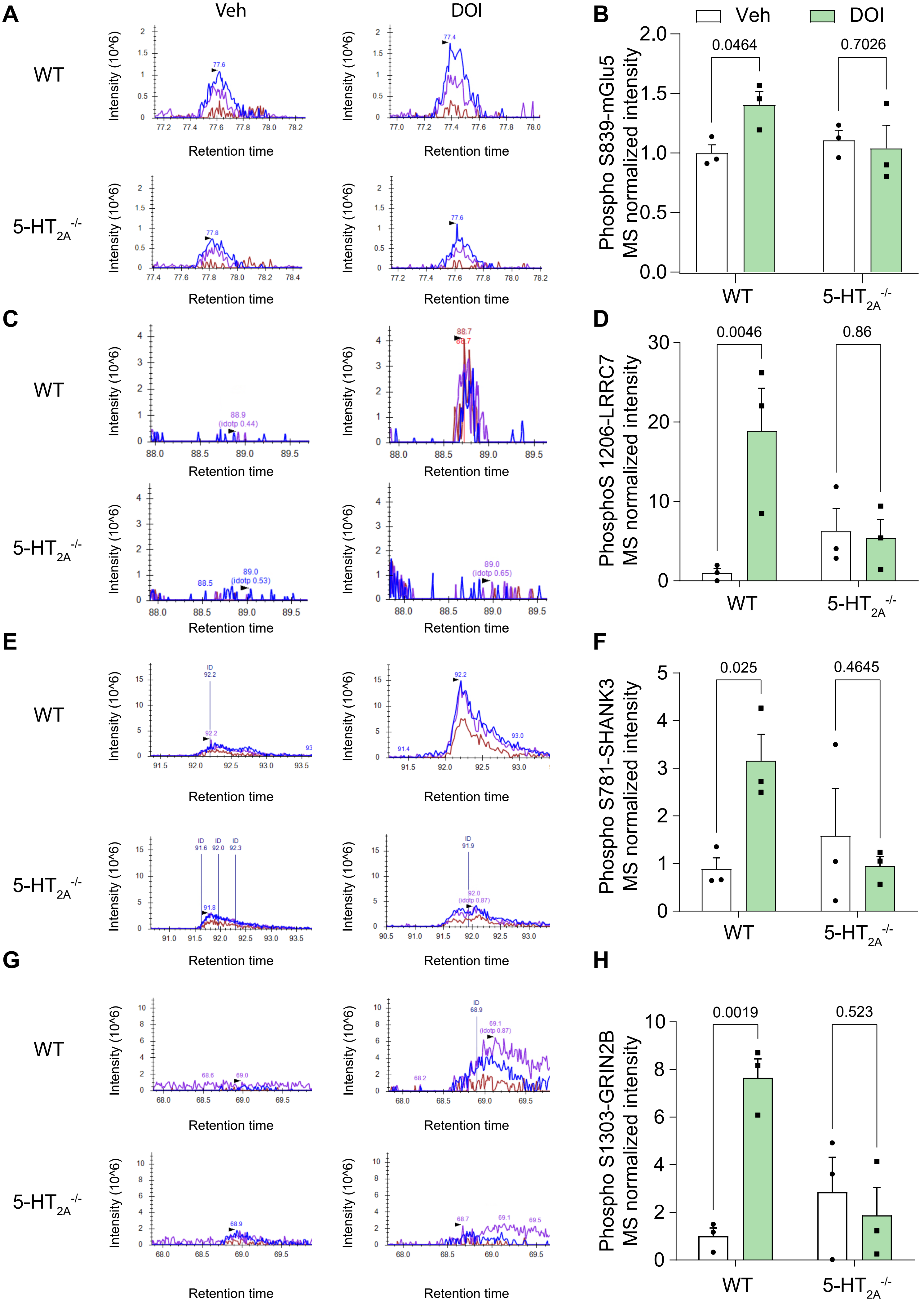

A
Veh
DOI
WT
5-HT2A-/-
Intensity (10^6)
Retention time
B
Veh
DOI
0.0464
0.7026
Phospho S839-mGlu5 MS normalized intensity
WT
5-HT2A-/-
C
D
0.0046
0.86
PhosphoS 1206-LRRC7 MS normalized intensity
E
F
0.025
0.4645
Phospho S781-SHANK3 MS normalized intensity
G
H
0.0019
0.523
Phospho S1303-GRIN2B MS normalized intensity

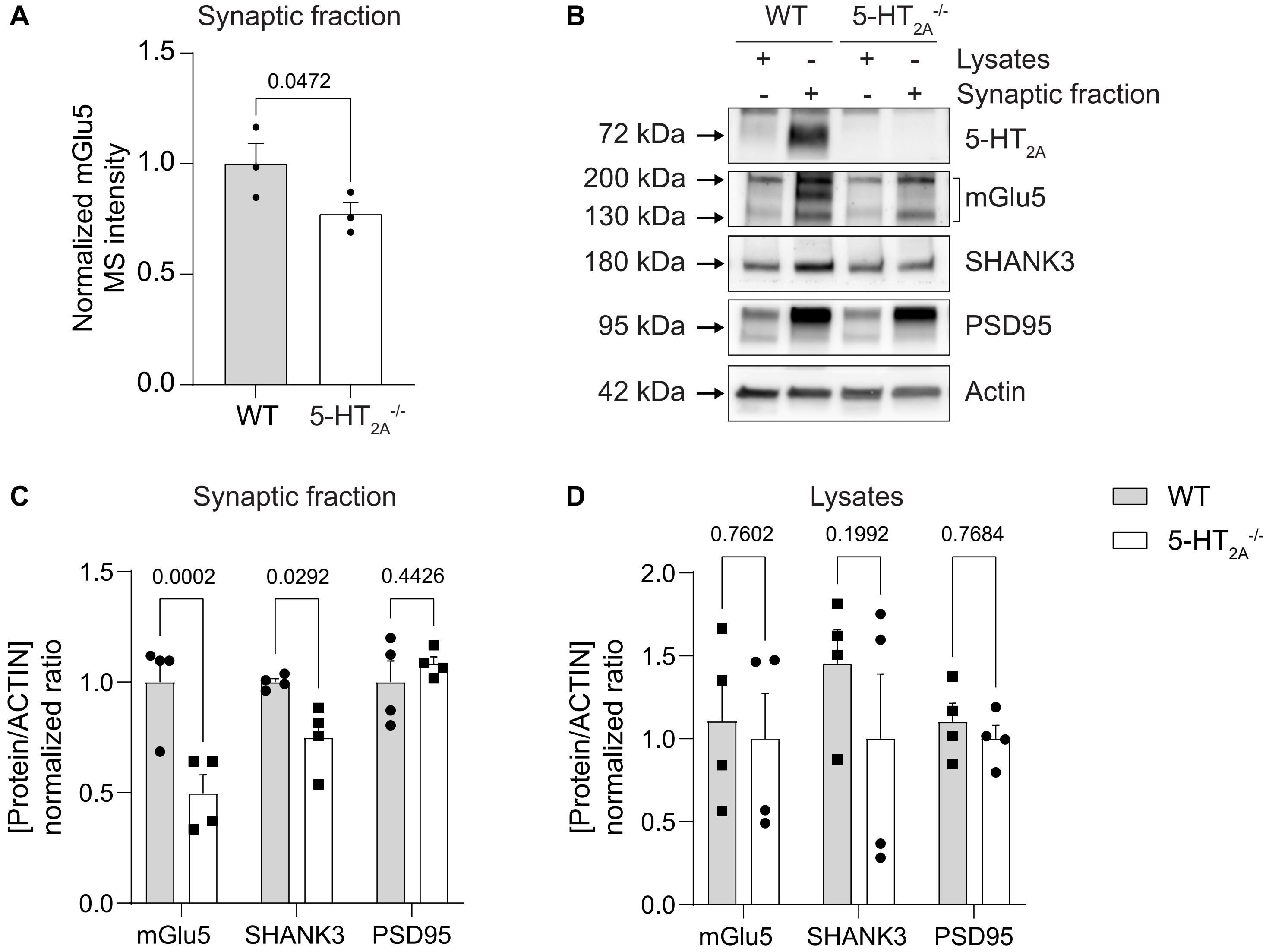
A
Synaptic fraction
0.0472
Normalized mGlu5 MS intensity
WT
5-HT2A-/-
B
WT
5-HT2A-/-
Lysates
Synaptic fraction
72 kDa
5-HT2A
200 kDa
130 kDa
mGlu5
180 kDa
SHANK3
95 kDa
PSD95
42 kDa
Actin
C
Synaptic fraction
0.0002
0.0292
0.4426
[Protein/ACTIN] normalized ratio
mGlu5
SHANK3
PSD95
D
Lysates
0.7602
0.1992
0.7684
WT
5-HT2A-/-

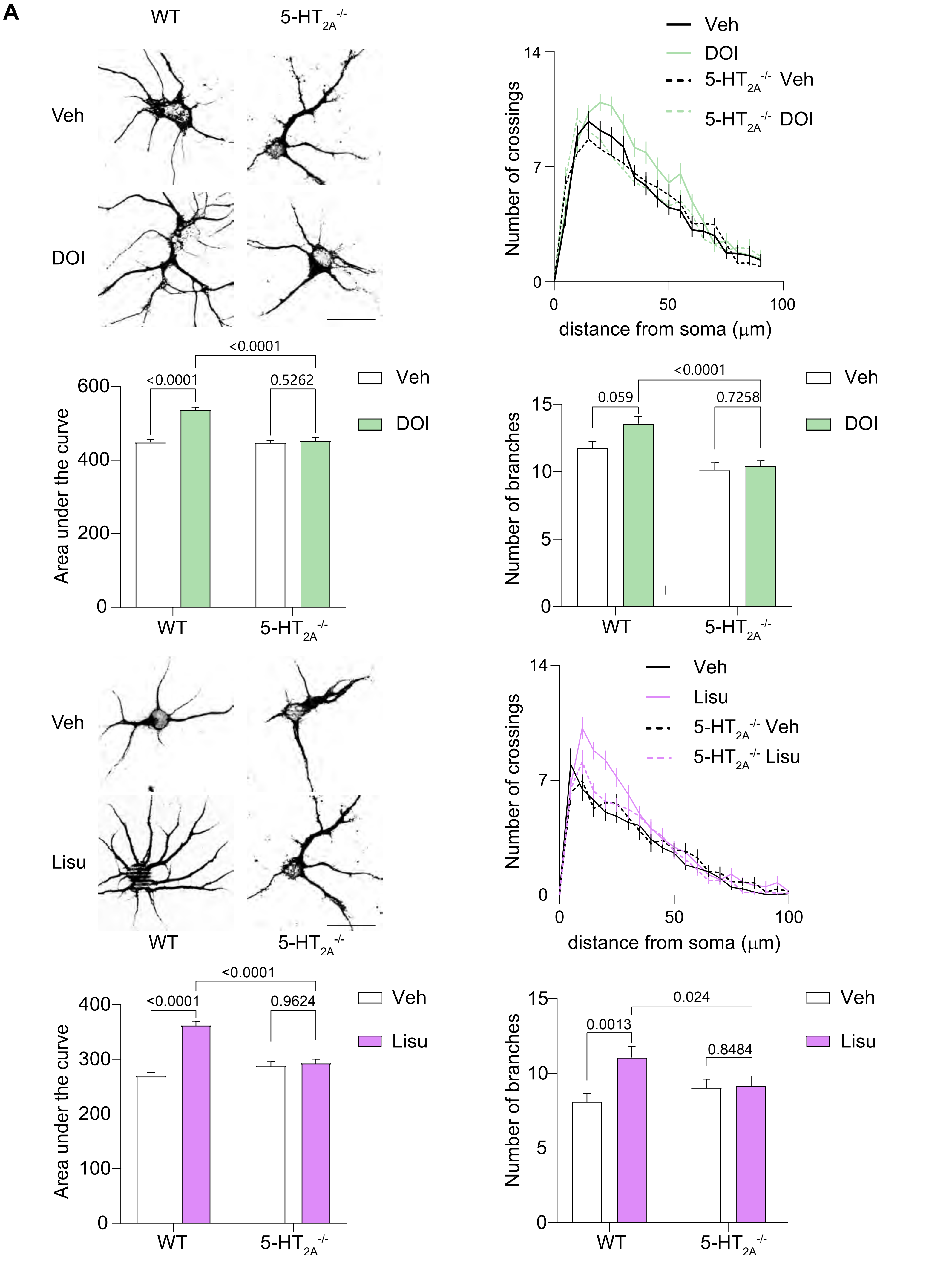

A
WT
5-HT2A-/-
Veh
DOI
Lisu
Number of crossings
distance from soma (μm)
5-HT2A-/- Veh
5-HT2A-/- DOI
5-HT2A-/- Lisu
Area under the curve
Number of branches
<0.0001
0.5262
0.059
0.7258
0.9624
0.0013
0.024
0.8484

| peptide ID | Protein | Phosphosite position | Protein name | Gene name | Peptide sequence | Phosphosite Probabilities | Peptide Intensity WT Veh_1 | Peptide Intensity WT Veh_2 | Peptide Intensity WT Veh_3 | Peptide Intensity WT DOI_1 | Peptide Intensity WT DOI_2 | Peptide Intensity WT DOI_3 | Peptide Intensity HTR2A-/- Veh_1 | Peptide Intensity HTR2A-/- Veh_2 | Peptide Intensity HTR2A-/- Veh_3 | Peptide Intensity HTR2A-/- DOI_1 | Peptide Intensity HTR2A-/- DOI_2 | Peptide Intensity HTR2A-/- DOI_3 | Peptide Normalized intensity WT Veh_1 | Peptide Normalized intensity WT Veh_2 | Peptide Normalized intensity WT Veh_3 | Peptide Normalized intensity WT DOI_1 | Peptide Normalized intensity WT DOI_2 | Peptide Normalized intensity WT DOI_3 | Peptide Normalized intensity HTR2A-/- Veh_1 |
|---|---|---|---|---|---|---|---|---|---|---|---|---|---|---|---|---|---|---|---|---|---|---|---|---|---|
| 760 | P14873 | 1438 | crotubule-associated protein 1B;MAP1B heavy chain;MAP1 light chain L | Map1b | PFEGKNGKQGFPDRESPVSDLTSTGLYQDKQ | QGFPDRES(0.999)PVS(0.001)DLTSTGLYQDKQEEK | 0 | 0 | 0 | 17798000 | 21833000 | 10812000 | 20064000 | 9333900 | 0 | 12505000 | 22121000 | 10312000 | 0 | 0 | 0 | 0,153355686 | 0,166215524 | 0,108511586 | 0,178791481 |
| 1337 | O88935 | 67 | Synapsin-1 | Syn1 | ERASTAAPVASPAAPSPGSSGGGGFFSSLSN | ASTAAPVASPAAPS(0.997)PGS(0.002)SGGGGFFSSLSNAVK | 24079000 | 19845000 | 23084000 | 65337000 | 71779000 | 61763000 | 65035000 | 0 | 8286200 | 43809000 | 46401000 | 39824000 | 0,19574548 | 0,132573286 | 0,184406627 | 0,562973393 | 0,546456469 | 0,619866914 | 0,5795307 |
| 575 | Q3UNH4 | 691 | G protein-regulated inducer of neurite outgrowth 1 | Gprin1 | ASTSQKDLAAAAAQKSPSAEAAAPPPGPRTR | S(0.961)PS(0.039)AEAAAPPPGPR | 12286000 | 10531000 | 11349000 | 34391000 | 38256000 | 38808000 | 36440000 | 20667000 | 8441100 | 28284000 | 35752000 | 26796000 | 0,099876613 | 0,070351689 | 0,090661532 | 0,296328542 | 0,291244496 | 0,389485536 | 0,324718978 |
| 49 | Q9QYC0 | 610 | Alpha-adducin | Add1 | EQKEKSPPDQSAVPNTPPSTPVKLEEDLPQE | SPPDQS(0.001)AVPNT(0.996)PPS(0.528)T(0.476)PVKLEEDLPQEPTSR | 0 | 2061300 | 0 | 7563500 | 11170000 | 11270000 | 8754500 | 5880000 | 11734000 | 11259000 | 21587000 | 5387500 | 0 | 0,013770386 | 0 | 0,065170566 | 0,085037668 | 0,113108173 | 0,078011863 |
| 620 | B5TVM2 | 487 | Immunoglobulin-like domain-containing receptor 2 | Ildr2 | ERSESRAHGAFYQDGSLDEYYGRGRSREPPG | AHGAFYQDGS(1)LDEYYGR | 0 | 1977400 | 0 | 5952800 | 6021400 | 4215500 | 10199000 | 6294900 | 1944800 | 7906500 | 7318000 | 4717700 | 0 | 0,013209898 | 0 | 0,05129204 | 0,045841165 | 0,042307676 | 0,090883887 |
| 1109 | Q9ES97 | 31 | Reticulon-3 | Rtn3 | SSSGAEPSALGGGGGSPGACPALGAKSCGSS | AESSAATQSPSVSSSSSGAEPSALGGGGGS(1)PGACPALGAK | 0 | 0 | 0 | 21179000 | 15286000 | 23849000 | 25698000 | 15303000 | 9548200 | 20113000 | 26857000 | 14297000 | 0 | 0 | 0 | 0,182487924 | 0,116372945 | 0,239353756 | 0,228996386 |
| 779 | P20357 | 626 | Microtubule-associated protein 2 | Map2 | NQHDEKELQAKASQPSPPAQEAGYSTLAQSY | AS(0.154)QPS(0.795)PPAQEAGY(0.029)S(0.004)T(0.004)LAQS(0.005)YE | 0 | 0 | 0 | 22126000 | 13783000 | 19740000 | 22247000 | 20198000 | 18223000 | 20521000 | 15110000 | 16011000 | 0 | 0 | 0 | 0,190647708 | 0,104930544 | 0,198114937 | 0,198244322 |
| 904 | P13595 | 958 | Neural cell adhesion molecule 1 | Ncam1 | SKASPAPTPTPAGAASPLAAVAAPATDAPQA | ASPAPT(0.001)PT(0.007)PAGAAS(0.991)PLAAVAAPATDAPQAK | 3664800 | 3218900 | 4636700 | 12692000 | 13703000 | 12745000 | 16605000 | 5353000 | 0 | 5999100 | 9614300 | 8281600 | 0,029792269 | 0,021503661 | 0,037040297 | 0,109360061 | 0,104321501 | 0,127911595 | 0,147968129 |
| 360 | Q71M36 | 543 | Chondroitin sulfate proteoglycan 5 | Cspg5 | SRLKEEESFNIQNSMSPKLEGGKGDQDDLGV | LKEEESFNIQNS(0.013)MS(0.987)PK | 0 | 0 | 0 | 10684000 | 12607000 | 16369000 | 27681000 | 6366700 | 0 | 14681000 | 10705000 | 8804300 | 0 | 0 | 0 | 0,092058217 | 0,095977608 | 0,164282847 | 0,246667015 |
| 5 | P28661 | 68 | Septin-4 | sept-04 | KTRVARPQILEPRPQSPDLCDDDVEFRGSLW | VARPQILEPRPQS(1)PDLCDDDVEFR | 0 | 3931000 | 0 | 19868000 | 15040000 | 17955000 | 14239000 | 5493200 | 0 | 6951600 | 8113400 | 8832600 | 0 | 0,026260801 | 0 | 0,17119175 | 0,114500136 | 0,180200289 | 0,126884564 |
| 964 | A0A0A6YY83 | 1011 | | Pcdh7 | YKSSSPLPTVQLHPQSPTAGKKHQAVQDLPP | SSSPLPT(0.016)VQLHPQS(0.784)PT(0.199)AGKK | 0 | 2793100 | 0 | 6744000 | 12029000 | 7093000 | 0 | 5566400 | 0 | 7046200 | 5969300 | 0 | 0 | 0,018659131 | 0 | 0,05810938 | 0,09157727 | 0,071186892 | 0 |
| 249 | O88737 | 2860 | Protein bassoon | Bsn | QQTLPRPMKTLQRSLSDPKPLSPTAEESAKE | S(0.012)LS(0.988)DPKPLS(0.998)PT(0.002)AEESAK | 5744900 | 4281200 | 5992500 | 20049000 | 19273000 | 22586000 | 16290000 | 32982000 | 6593300 | 15562000 | 22432000 | 12938000 | 0,046702031 | 0,0286029 | 0,04787111 | 0,172751329 | 0,146718526 | 0,226678013 | 0,145161146 |
| 1350 | O88935 | 553 | Synapsin-1 | Syn1 | GGPGAPPAARPPASPSPQRQAGAPQATRQAS | PVAGGPGAPPAARPPAS(1)PS(1)PQR | 16272000 | 12449000 | 14764000 | 56361000 | 79495000 | 32766000 | 9745700 | 64488000 | 3350100 | 83030000 | 65716000 | 40117000 | 0,132280014 | 0,083164769 | 0,117942274 | 0,485632083 | 0,605198693 | 0,328846709 | 0,086844504 |
| 164 | Q6PIE5 | 439 | Sodium/potassium-transporting ATPase subunit alpha-2 | Atp1a2 | GLCNRAVFKAGQENISVSKRDTAGDASESAL | AGQENIS(0.908)VS(0.092)KR | 7156200 | 7856400 | 7593700 | 16038000 | 12549000 | 12626000 | 22230000 | 8460600 | 5362900 | 18165000 | 10461000 | 10496000 | 0,058174916 | 0,052484191 | 0,060662303 | 0,138190723 | 0,095536051 | 0,126717285 | 0,198092834 |
| 250 | O88737 | 2866 | Protein bassoon | Bsn | PMKTLQRSLSDPKPLSPTAEESAKERFSLYQ | S(0.012)LS(0.988)DPKPLS(0.998)PT(0.002)AEESAK | 1954500 | 0 | 0 | 6262300 | 7794500 | 7491500 | 6033500 | 14794000 | 3225200 | 6854900 | 9332100 | 6219200 | 0,015888722 | 0 | 0 | 0,053958833 | 0,059339848 | 0,075186325 | 0,053764872 |
| 1375 | Q8CHC4 | 1160 | Synaptojanin-1 | Synj1 | PPSPGVARREIEAPKSPGTARKDNIGRNQPS | EIEAPKS(0.986)PGT(0.014)AR | 0 | 2130100 | 1780500 | 3515800 | 4427200 | 5072900 | 7217600 | 4409500 | 2715500 | 0 | 7054200 | 3989500 | 0 | 0,01423 | 0,014223531 | 0,030293736 | 0,033704455 | 0,050912729 | 0,064316457 |
| 770 | P14873 | 1395 | crotubule-associated protein 1B;MAP1B heavy chain;MAP1 light chain L | Map1b | PDSESPVEKVLSPLRSPPLLGSESPYEDFLS | VLS(1)PLRS(1)PPLLGSESPYEDFLSADSK | 5175000 | 0 | 0 | 32449000 | 32951000 | 32503000 | 1178000 | 21527000 | 0 | 25404000 | 53484000 | 33147000 | 0,042069142 | 0 | 0 | 0,279595384 | 0,250857314 | 0,326207184 | 0,010497227 |
| 769 | P14873 | 1391 | crotubule-associated protein 1B;MAP1B heavy chain;MAP1 light chain L | Map1b | DEPVPDSESPVEKVLSPLRSPPLLGSESPYE | VLS(1)PLRS(1)PPLLGSESPYEDFLSADSK | 5175000 | 0 | 0 | 32449000 | 32951000 | 32503000 | 1178000 | 21527000 | 0 | 25404000 | 53484000 | 33147000 | 0,042069142 | 0 | 0 | 0,279595384 | 0,250857314 | 0,326207184 | 0,010497227 |
| 606 | Q640R3 | 280 | Hepatocyte cell adhesion molecule | Hepacam | PSKKSRKKRKLEKQNSLEYMDQNDDRLKSEA | QNS(1)LEYMDQNDDRLK | 0 | 3114300 | 0 | 13002000 | 9609400 | 5432800 | 22433000 | 8904200 | 13355000 | 31366000 | 0 | 10564000 | 0 | 0,020804219 | 0 | 0,112031162 | 0,073156756 | 0,054524764 | 0,199901779 |
| 93 | Q8C8R3 | 846 | Ankyrin-2 | Ank2 | HKLNVPETMTEVLDVSDEEGDDTVTGDGGEY | HKLNVPETMTEVLDVS(1)DEEGDDTVTGDGGEYLRPEDLK | 2922400 | 0 | 0 | 18622000 | 11851000 | 10190000 | 30445000 | 16008000 | 7989600 | 16390000 | 0 | 5930300 | 0,023757074 | 0 | 0 | 0,160455646 | 0,090222149 | 0,102269058 | 0,271297181 |
| 280 | Q61290 | 948 | Voltage-dependent R-type calcium channel subunit alpha-1E | Cacna1e | DSASASRSRSASQERSLDEGVSVEGEKEHEP | S(1)LDEGVSVEGEKEHEPHSSHR | 0 | 5377800 | 0 | 14752000 | 15983000 | 13278000 | 31511000 | 16913000 | 9321800 | 4993000 | 14078000 | 10226000 | 0 | 0,035926058 | 0 | 0,127109961 | 0,121679234 | 0,133260899 | 0,28079637 |
| 732 | Q9QYR6 | 1648 | crotubule-associated protein 1A;MAP1A heavy chain;MAP1 light chain L | Map1a | TSPTRGEPVPAWEGKSPEQEVRYWRDRDITL | ET(0.372)S(0.487)PT(0.141)RGEPVPAWEGKS(1)PEQEVR | 2728600 | 3521800 | 4399600 | 13217000 | 9040600 | 12816000 | 21048000 | 17497000 | 7999900 | 18564000 | 20473000 | 8779000 | 0,022181615 | 0,023527165 | 0,035146222 | 0,1138837 | 0,068826458 | 0,128624166 | 0,187559963 |
| 1353 | O88935 | 666 | Synapsin-1 | Syn1 | AAAGGPPHPQLNKSQSLTNAFNLPEPAPPRP | SQS(1)LTNAFNLPEPAPPRPSLSQDEVK | 54816000 | 47163000 | 92257000 | 213290000 | 232350000 | 183290000 | 255240000 | 99689000 | 40510000 | 148190000 | 255120000 | 137130000 | 0,445615858 | 0,315069483 | 0,736995417 | 1,837803925 | 1,768890074 | 1,839538342 | 2,274458614 |
| 453 | Q9Z2H5 | 489 | Band 4.1-like protein 1 | Epb41l1 | RTPTKIKELKPEQETTPRHKQEFLDKPEDVL | ELKPEQET(0.177)T(0.823)PR | 5484300 | 5892000 | 9867700 | 17813000 | 25844000 | 19391000 | 18797000 | 11770000 | 3588400 | 23363000 | 20809000 | 19283000 | 0,044583535 | 0,039361139 | 0,078828161 | 0,153484933 | 0,196751431 | 0,194612297 | 0,16750117 |
| 665 | Q08460 | 711 | Calcium-activated potassium channel subunit alpha-1 | Kcnma1 | KCGCRRLEDEQPPTLSPKKKQRNGGMRNSPN | RLEDEQPPTLS(1)PK | 8553200 | 9554100 | 12197000 | 25869000 | 20760000 | 24785000 | 27023000 | 16698000 | 6365500 | 20711000 | 24296000 | 16375000 | 0,069531552 | 0,06382557 | 0,097435784 | 0,222899103 | 0,158046731 | 0,248747656 | 0,240803538 |
| 1242 | Q61165 | 697 | Sodium/hydrogen exchanger 1 | Slc9a1 | QKITNYLTVPAHKLDSPTLSRARIGSDPLAY | ITNYLTVPAHKLDS(0.772)PT(0.076)LS(0.152)R | 0 | 3478700 | 0 | 5974800 | 13146000 | 8629500 | 9690900 | 2543200 | 0 | 6525100 | 4531300 | 6612800 | 0 | 0,023239239 | 0 | 0,051481602 | 0,100081037 | 0,086607541 | 0,086356178 |
| 1401 | Q9CPQ3 | 15 | Mitochondrial import receptor subunit TOM22 homolog | Tomm22 | _MAAAVAAAGAGEPLSPEELLPKAEAEKAEE | AAAVAAAGAGEPLS(1)PEELLPK | 0 | 1505700 | 0 | 4632900 | 11961000 | 9046000 | 5129100 | 1746900 | 4613000 | 2012200 | 5325900 | 4572400 | 0 | 0,010058735 | 0 | 0,03991918 | 0,091059583 | 0,090787625 | 0,045705711 |
| 248 | O88737 | 789 | Protein bassoon | Bsn | KRRPHSLSITPEAFDSDEELGDILEEDDSLA | RPHSLSIT(0.001)PEAFDS(0.999)DEELGDILEEDDSLAWGR | 0 | 4511400 | 0 | 10813000 | 9447700 | 10778000 | 5001000 | 0 | 0 | 4180800 | 0 | 7552100 | 0 | 0,030138127 | 0 | 0,09316974 | 0,071925727 | 0,108170354 | 0,044564204 |
| 605 | O88703 | 842 | ssium/sodium hyperpolarization-activated cyclic nucleotide-gated chan | Hcn2 | TARTAAPSPDRRDSASPGAAGGLDPLDSARS | DS(0.034)AS(0.966)PGAAGGLDPLDSAR | 0 | 1684300 | 0 | 0 | 0 | 0 | 4752300 | 0 | 0 | 0 | 0 | 0 | 0 | 0,011251861 | 0 | 0 | 0 | 0 | 0,042348024 |
| 38 | P51830 | 610 | Adenylate cyclase type 9 | Adcy9 | DDSRESSGPRGQGTASPGSVSDLAQTVKTFD | GQGT(0.017)AS(0.983)PGSVSDLAQTVK | 4975300 | 4712600 | 0 | 13698000 | 12461000 | 14256000 | 0 | 0 | 2418100 | 14494000 | 0 | 12105000 | 0,04044572 | 0,031482231 | 0 | 0,118028216 | 0,094866104 | 0,143076319 | 0 |
| 1420 | Q3UWE6 | 357 | | Wdr20 | RDRANSTQSRLSKRNSTDSRPVSVTYRFGSV | RNS(0.857)T(0.142)DS(0.001)RPVSVTYR | 0 | 0 | 0 | 7970900 | 0 | 5710000 | 10378000 | 0 | 2058200 | 6463800 | 5493800 | 3289900 | 0 | 0 | 0 | 0,06868091 | 0,043470464 | 0,039376097 | 0,092478967 |
| 961 | Q8CFX3 | 951 | | Pcdh1 | YPPGSPDLGRHYRSNSPLPSIQLQPQSPSAS | S(0.062)NS(0.937)PLPS(0.002)IQLQPQS(0.779)PS(0.17)AS(0.05)KK | 2753400 | 2725700 | 4979300 | 9142000 | 11176000 | 8334900 | 0 | 10940000 | 1681100 | 0 | 15284000 | 10095000 | 0,022383222 | 0,018208869 | 0,039777158 | 0,078771642 | 0,085083346 | 0,083650871 | 0 |
| 1182 | P43006 | 562 | Excitatory amino acid transporter 2 | Slc1a2 | ECKVTLAANGKSADCSVEEEPWKREK_____ | SADCS(1)VEEEPWKR | 3484400 | 5409200 | 4175100 | 15627000 | 9436400 | 15993000 | 12773000 | 11849000 | 5837300 | 12835000 | 10527000 | 10057000 | 0,028325742 | 0,036135824 | 0,033552803 | 0,13464936 | 0,0718397 | 0,16050923 | 0,113820952 |
| 621 | Q5DU25 | 402 | IQ motif and SEC7 domain-containing protein 2 | Iqsec2 | YEKAQNPAYFEGKPASLDEGAMAGARSHRLE | AQNPAYFEGKPAS(1)LDEGAMAGAR | 3598800 | 5057800 | 0 | 13520000 | 12200000 | 22383000 | 0 | 0 | 3000100 | 0 | 9161700 | 6930600 | 0,029255735 | 0,033788318 | 0 | 0,116494487 | 0,0928791 | 0,224640661 | 0 |
| [illegible] | | | | | | | | | | | | | | | | | | | | | | | | | |

| | | | | | | | | | | | | | | | | | | | | | | | | | |
|---|---|---|---|---|---|---|---|---|---|---|---|---|---|---|---|---|---|---|---|---|---|---|---|---|---|
| 1309 | O70439 | 79 | Syntaxin-7 | Stx7 | KETDKYIKEFGSLPTTPSEQRQRKIQKDRLV | EFGSLPT(0.138)T(0.724)PS(0.138)EQR | 0 | 0 | 4341600 | 6207100 | 12004000 | 8221900 | 10351000 | 3422100 | 2634600 | 6428000 | 11690000 | 6423600 | 0 | 0 | 0,034682889 | 0,053483205 | 0,091386944 | 0,082516778 | 0,092238368 |
| 1143 | B1AWN6;A2ASI5 | 610;610 | Sodium channel protein | Scn2a1;Scn3a | DDEHSTFEDNDSRRDSLFVPHRHGERRPSNV | RDS(1)LFVPHR | 0 | 2629700 | 0 | 12516000 | 8369800 | 0 | 16549000 | 1584900 | 2432800 | 3689300 | 10761000 | 0 | 0 | 0,017567547 | 0 | 0,107843565 | 0,06371963 | 0 | 0,14746911 |
| 730 | Q9QYR6 | 499 | crotubule-associated protein 1A;MAP1A heavy chain;MAP1 light chain L | Map1a | LKVDKGRAARGEKELSSEPRTPPAQKGAAPP | ELS(0.78)S(0.079)EPRT(0.141)PPAQK | 0 | 2482600 | 0 | 8321300 | 6109900 | 2642400 | 8169600 | 3986600 | 2080400 | 6412200 | 0 | 5979100 | 0 | 0,016584855 | 0 | 0,071700116 | 0,046514919 | 0,026519702 | 0,07279785 |
| 182 | Q9R0K7 | 18 | Plasma membrane calcium-transporting ATPase 2 | Atp2b2 | DMTNSDFYSKNQRNESSHGGEFGCTMEELRS | NES(0.978)S(0.022)HGGEFGCTMEELR | 0 | 0 | 0 | 14173000 | 0 | 7825400 | 0 | 0 | 0 | 0 | 5249600 | 2792200 | 0 | 0 | 0 | 0,122121033 | 0 | 0,078537418 | 0 |
| 1008 | Q6R891 | 100 | Neurabin-2 | Ppp1r9b | RASDRGVRLSLPRASSLNENVDHSALLKLGT | AS(0.01)S(0.99)LNENVDHSALLK | 4469900 | 4285700 | 5309100 | 10430000 | 7442400 | 12763000 | 12476000 | 5875200 | 4074300 | 9301300 | 10354000 | 11009000 | 0,03633717 | 0,028630352 | 0,042411767 | 0,089869637 | 0,056659296 | 0,128092246 | 0,111174368 |
| 636 | Q03717 | 484 | Potassium voltage-gated channel subfamily B member 1 | Kcnb1 | GEGVAKKDKVQDNHLSPNKWKWTKRALSETS | DKVQDNHLS(1)PNKWK | 0 | 0 | 0 | 0 | 0 | 5395200 | 15402000 | 8930100 | 0 | 12497000 | 6833800 | 2029900 | 0 | 0 | 0 | 0 | 0 | 0,054147402 | 0,137248126 |
| 1394 | Q8BG50 | 20 | Transmembrane protein 169 | Tmem169 | APVESQGQLPSPHHGSLRRAVAAVLALDGES | MEES(0.048)APVES(0.051)QGQLPS(0.955)PHHGS(0.946)LRR | 0 | 0 | 0 | 0 | 8179400 | 0 | 12072000 | 13522000 | 7624500 | 14156000 | 12313000 | 10811000 | 0 | 0 | 0 | 0 | 0,062270107 | 0 | 0,1075743 |
| 1393 | Q8BG50 | 15 | Transmembrane protein 169 | Tmem169 | _MEESAPVESQGQLPSPHHGSLRRAVAAVLA | MEES(0.048)APVES(0.051)QGQLPS(0.955)PHHGS(0.946)LRR | 0 | 0 | 0 | 0 | 8179400 | 0 | 12072000 | 13522000 | 7624500 | 14156000 | 12313000 | 10811000 | 0 | 0 | 0 | 0 | 0,062270107 | 0 | 0,1075743 |
| 311 | Q3UHB8 | 310 | Coiled-coil domain-containing protein 177 | Ccdc177 | VPLTARSFSLGDLSHSPQTAQHVERIVRQVR | S(0.027)FS(0.949)LGDLS(0.175)HS(0.836)PQT(0.014)AQHVER | 0 | 1552600 | 0 | 0 | 0 | 5856800 | 7836200 | 10729000 | 4593000 | 10514000 | 9362400 | 6417000 | 0 | 0,010372048 | 0 | 0 | 0 | 0,05878012 | 0,069828838 |
| 1039 | Q6PE13 | 804 | Proline-rich transmembrane protein 3 | Prrt3 | SLSELDLRPPSPINLSRSIDAALFREHLVRE | GS(0.001)VGPAPS(0.957)LS(0.042)ELDLRPPS(0.207)PINLS(0.793)R | 0 | 0 | 0 | 18990000 | 23065000 | 0 | 13971000 | 11628000 | 5529400 | 0 | 0 | 1150400 | 0 | 0 | 0 | 0,163626502 | 0,17559479 | 0 | 0,124496401 |
| 60 | Q01338 | 360 | Alpha-2A adrenergic receptor | Adra2a | DSLPRGPGAAGPGASGSGHGEERGGGAKAS | RGPGAAGPGAS(0.887)GS(0.113)GHGEER | 0 | 0 | 0 | 7266400 | 0 | 0 | 11134000 | 0 | 0 | 0 | 0 | 0 | 0 | 0 | 0 | 0,062610617 | 0 | 0 | 0,099215727 |
| 759 | P14873 | 1497 | crotubule-associated protein 1B;MAP1B heavy chain;MAP1 light chain L | Map1b | QSALALDERKLGGDVSPTQIDVSQFGSFKED | LGGDVS(1)PTQIDVSQFGSFK | 6234100 | 0 | 0 | 27184000 | 21572000 | 8459300 | 20549000 | 0 | 2392600 | 11793000 | 16102000 | 8097500 | 0,050678886 | 0 | 0 | 0,234229743 | 0,16422852 | 0,084899376 | 0,183113345 |
| 572 | Q3UNH4 | 764 | G protein-regulated inducer of neurite outgrowth 1 | Gprin1 | AGGPAFSFQAAPRAPSPAPRPPSRRDAGLQV | APS(1)PAPRPPS(1)RR | 2828500 | 3399800 | 8483300 | 12649000 | 13718000 | 17777000 | 17307000 | 7919300 | 3720600 | 4546200 | 14332000 | 13322000 | 0,022993733 | 0,022712152 | 0,067768876 | 0,108989553 | 0,104435696 | 0,178413842 | 0,154223692 |
| 338 | Q9JKC6 | 90 | Cell cycle exit and neuronal differentiation protein 1 | Cend1 | HSNLKPAPTVPAAPSSPDATSEPKGPGDGAE | AOPVLLNNHSNLKPAPTVPAAPS(0.064)S(0.928)PDAT(0.004)S(0.004)EPK | 40612000 | 33193000 | 85792000 | 195040000 | 119640000 | 153680000 | 218020000 | 0 | 53729000 | 62113000 | 106640000 | 49071000 | 0,330147242 | 0,221743769 | 0,685349738 | 1,680553601 | 0,910824224 | 1,542365936 | 1,94278901 |
| 1104 | P62071 | 186 | Ras-related protein R-Ras2 | Rras2 | LVRVIRKFQEQECPPSPEPTRKEKDKKGCHC | KFQEQECPPS(0.998)PEPT(0.002)RK | 17791000 | 13546000 | 15520000 | 33333000 | 22758000 | 26264000 | 43316000 | 19499000 | 10111000 | 38615000 | 22512000 | 15626000 | 0,144628425 | 0,090493209 | 0,123981583 | 0,287212332 | 0,173257587 | 0,263591222 | 0,385991417 |
| 187 | Q0VF55 | 8 | Calcium-transporting ATPase | Atp2b3 | ________MGDMANSSIEFHPKPQQQREVPH | GDMANS(0.033)S(0.967)IEFHPKPQQQR | 0 | 0 | 2190600 | 3898100 | 0 | 7316700 | 12180000 | 5963800 | 0 | 9236600 | 0 | 7788500 | 0 | 0 | 0,017499617 | 0,033587808 | 0 | 0,073431994 | 0,108536695 |
| 162 | Q8VDN2 | 47 | Sodium/potassium-transporting ATPase subunit alpha-1 | Atp1a1 | MDELKKEVSMDDHKLSLDELHRKYGTDLSRG | EVSMDDHKLS(1)LDELHRK | 2275600 | 2294400 | 4941900 | 14559000 | 6544200 | 13594000 | 24522000 | 5117700 | 3841300 | 17500000 | 1823900 | 6956800 | 0,018499041 | 0,015327596 | 0,039478388 | 0,125446985 | 0,049821263 | 0,136432343 | 0,218516981 |
| 1411 | Q8JZK6 | 248 | | Trim61 | ALLASVKDWYERYKQSGWPEMALVTLKGYDY | QS(0.997)GWPEMALVT(0.003)LK | 4648300 | 13327000 | 32620000 | 51626000 | 118360000 | 103220000 | 81290000 | 12065000 | 9452800 | 17459000 | 36879000 | 0 | 0,037787438 | 0,089030193 | 0,260585002 | 0,444833163 | 0,901079531 | 1,035938391 | 0,724379959 |
| 517 | Q80T41 | 883 | Gamma-aminobutyric acid type B receptor subunit 2 | Gabbr2 | TEPSRTCKDPIEDINSPEHIQRRLSLQLPIL | TCKDPIEDINS(1)PEHIQR | 53814000 | 37634000 | 55120000 | 138500000 | 83044000 | 85948000 | 120890000 | 64488000 | 17695000 | 83231000 | 84042000 | 64539000 | 0,437470296 | 0,251411592 | 0,440326343 | 1,193379172 | 0,632217376 | 0,862592839 | 1,077257882 |
| 47 | Q9QYC0 | 613 | Alpha-adducin | Add1 | EKSPPDQSAVPNTPPSTPVKLEEDLPQEPTS | SPPDQS(0.003)AVPNT(0.995)PPS(0.53)T(0.472)PVKLEEDLPQEPTSR | 0 | 2061300 | 0 | 7563500 | 0 | 0 | 8754500 | 5880000 | 11734000 | 11259000 | 21587000 | 5387500 | 0 | 0,013770386 | 0 | 0,065170566 | 0 | 0 | 0,078011863 |
| 413 | Q8BJ42 | 1047 | Disks large-associated protein 2 | Dlgap2 | RAASFRQNSATERADSIEIYIPEAQTRL___ | ADS(1)IEIYIPEAQTR | 1924900 | 4754100 | 8227000 | 15368000 | 14479000 | 21143000 | 10818000 | 17322000 | 1934400 | 4602100 | 17694000 | 7209100 | 0,015648095 | 0,031759469 | 0,065721423 | 0,132417698 | 0,11022922 | 0,212195751 | 0,096399833 |
| 483 | Q3TY60 | 322 | Protein FAM131B | Fam131b | ALSRKVSDVTSSGVQSFDEEEGDANN_____ | KVSDVTSS(0.001)GVQS(0.999)FDEEEGDANN | 2690100 | 0 | 4562200 | 12291000 | 8182900 | 15981000 | 24436000 | 23914000 | 0 | 17434000 | 24689000 | 19688000 | 0,021868637 | 0 | 0,036445153 | 0,105904862 | 0,062296753 | 0,160388795 | 0,21775063 |
| 448 | Q9Z2H5 | 684 | Band 4.1-like protein 1 | Epb41l1 | DESGGLEDSPDRGACSTPEMPQFESVKAETM | GACS(0.5)T(0.5)PEMPQFESVK | 0 | 0 | 0 | 7785100 | 0 | 0 | 0 | 0 | 0 | 0 | 0 | 0 | 0 | 0 | 0 | 0,067079973 | 0 | 0 | 0 |
| 510 | P39688 | 531 | Tyrosine-protein kinase Fyn | Fyn | LQGFLEDYFTATEPQYQPGENL_________ | KDPEERPTFEYLQGFLEDY(0.007)FT(0.008)AT(0.008)EPQY(0.978)QPGENL | 1620000 | 0 | 5840500 | 8982800 | 16874000 | 30815000 | 0 | 4154000 | 2536600 | 8274200 | 0 | 4812400 | 0,01316947 | 0 | 0,046656858 | 0,077399902 | 0,128462454 | 0,309266048 | 0 |
| 959 | Q8CFX3 | 927 | | Pcdh1 | QKSLKFNLMSDAPGDSPRIHLPLNYPPGSPD | FNLMSDAPGDS(1)PR | 0 | 0 | 0 | 0 | 0 | 5997500 | 6127800 | 2372500 | 0 | 4588700 | 4634500 | 3336000 | 0 | 0 | 0 | 0 | 0 | 0,060192216 | 0,054605185 |
| 165 | Q6PIC6 | 218 | Sodium/potassium-transporting ATPase subunit alpha-3 | Atp1a3 | VDNSSLTGESEPQTRSPDCTHDNPLETRNIT | S(1)PDCTHDNPLETR | 0 | 1752700 | 0 | 0 | 0 | 6284500 | 8916900 | 4124600 | 0 | 4919200 | 0 | 4529400 | 0 | 0,011708803 | 0 | 0 | 0 | 0,06307261 | 0,079459019 |
| 408 | Q9JLM8 | 337 | Serine/threonine-protein kinase DCLK1 | Dclk1 | LSTPRSGKSPSPSPTSPGSLRKQRISQHGGS | SPSPSPT(0.153)S(0.845)PGS(0.002)LRK | 0 | 0 | 0 | 0 | 8314600 | 13616000 | 0 | 8349600 | 0 | 0 | 10401000 | 0 | 0 | 0 | 0 | 0 | 0,063297868 | 0,13665314 | 0 |
| 530 | P06837 | 96 | Neuromodulin | Gap43 | KEGDGSATTDAAPATSPKAEEPSKAGDAPSE | KEGDGSATTDAAPAT(0.001)S(0.999)PK | 637130000 | 746450000 | 787460000 | 1872700000 | 996850000 | 1372300000 | 2566100000 | 1316300000 | 1101800000 | 2647700000 | 2386100000 | 1780600000 | 5,17942264 | 4,98661272 | 6,290627389 | 16,13603737 | 7,589059909 | 13,77270155 | 22,86666764 |
| 734 | Q9QYR6 | 991 | crotubule-associated protein 1A;MAP1A heavy chain;MAP1 light chain L | Map1a | EERCLSPDDSTVKMASPPPSGPPSAAHTPFH | MAS(0.998)PPPS(0.001)GPPSAAHTPFHQSPVEEK | 0 | 0 | 0 | 0 | 17885000 | 17122000 | 0 | 0 | 0 | 26011000 | 0 | 12830000 | 0 | 0 | 0 | 0 | 0,136159238 | 0,171840119 | 0 |
| 1373 | F6SEU4 | 1201 | Ras/Rap GTPase-activating protein SynGAP | Syngap1 | ESRLDRVKEYEEEIHSLKERLHMSNRKLEEY | VKEYEEEIHS(1)LKER | 0 | 0 | 2791800 | 11907000 | 7505600 | 0 | 11849000 | 7859500 | 0 | 0 | 0 | 0 | 0 | 0 | 0,022302306 | 0,102596143 | 0,05714044 | 0 | 0,105587134 |
| 1436 | A0A2R8VKG2 | 1753 | | | DLLVTRERPRRGIRNSGHDTEPEIIEETNVD | NS(0.577)GH(0.013)DT(0.41)EPEIIEETNVDR | 0 | 0 | 0 | 8070600 | 0 | 0 | 0 | 0 | 0 | 0 | 0 | 0 | 0 | 0 | 0 | 0,069539971 | 0 | 0 | 0 |
| 908 | P13595 | 926 | Neural cell adhesion molecule 1 | Ncam1 | SNLSSTVLANQGAVLSPSTPASAGETSKAPP | VAPLVDLS(0.001)DT(0.001)PT(0.006)S(0.006)APS(0.052)AS(0.052)NLS( | 0 | 0 | 0 | 19412000 | 0 | 14200000 | 0 | 0 | 0 | 8896000 | 0 | 0 | 0 | 0 | 0 | 0,167262646 | 0 | 0,142514291 | 0 |
| 874 | P04370 | 146 | Myelin basic protein | Mbp | LDVMASQKRPSQRSKYLATASTMDHARHGFL | S(0.225)KY(0.719)LAT(0.054)AS(0.001)T(0.001)MDHAR | 0 | 0 | 0 | 16164000 | 0 | 17221000 | 20029000 | 0 | 9106700 | 0 | 0 | 0 | 0 | 0 | 0 | 0,139276397 | 0 | 0,172833705 | 0,178479594 |
| 1289 | Q9QWI6 | 1110 | SRC kinase signaling inhibitor 1 | Srcin1 | VVTSKKDSVFIKKAESEELEVQKPQVKLRRA | KAES(1)EELEVQKPQVK | 35756000 | 20343000 | 18718000 | 82750000 | 47894000 | 46970000 | 66688000 | 51408000 | 21767000 | 89464000 | 66859000 | 33898000 | 0,290671348 | 0,202704521 | 0,149528819 | 0,713011744 | 0,364618985 | 0,471401145 | 0,59426068 |
| 569 | Q8C419 | 1059 | Probable G-protein coupled receptor 158 | Gpr158 | HKSKATEGLYQANHKSIDKTEVCPWEIHSQS | S(1)IDKTEVCPWEIHSQSLLEDENR | 3218600 | 10727000 | 20994000 | 38311000 | 48778000 | 53899000 | 36194000 | 16957000 | 5807200 | 23153000 | 25283000 | 17119000 | 0,026164974 | 0,071661055 | 0,167710654 | 0,33010505 | 0,371348913 | 0,540942098 | 0,322526857 |
| [illegible] | [illegible] | [illegible] | [illegible] | [illegible] | [illegible] | [illegible] | [illegible] | [illegible] | [illegible] | [illegible] | [illegible] | [illegible] | [illegible] | [illegible] | [illegible] | [illegible] | [illegible] | [illegible] | [illegible] | [illegible] | [illegible] | [illegible] | [illegible] | [illegible] | [illegible] |

| | | | | | | | |
|---|---|---|---|---|---|---|---|
| 1037 | Q6PE13 | 791 | Proline-rich transmembrane protein 3 | Prrt3 | GQRMSRGSVGPAPSLSELDLRPPSPINLSRS | GS(0.022)VGPAPS(0.136)LS(0.843)ELDLRPPS(0.884)PINLS(0.115)R | 2126900 |
| 1407 | Q7TQD2 | 31 | Tubulin polymerization-promoting protein | Tppp | PPKSPGDPARAAKRLSLESEGANEGATAAPE | RLS(1)LESEGANEGATAAPELSALEEAFRR | 37736000 |
| 28 | Q9R1V6 | 832 | Disintegrin and metalloproteinase domain-containing protein 22 | Adam22 | KHISDICENGRPRSNSWQGNMGGNKKKIRGK | S(0.035)NS(0.965)WQGNMGGNK | 0 |
| 463 | Q9WV92 | 96 | Band 4.1-like protein 3;Band 4.1-like protein 3, N-terminally processed | Epb41l3 | AAKQLEYQQFEDDKLSQRSSSSKLSRSPLKI | QLEYQQFEDDKLS(1)QR | 6919500 |
| 797 | P20357 | 1560 | Microtubule-associated protein 2 | Map2 | SLPRPSSILPPRRGVSGDREENSFSLNSSIS | RGVS(0.973)GDREENS(0.02)FS(0.006)LNSSISSAR | 4786700 |
| 320 | Q8BGU5 | 25 | Cyclin-Y | Ccny | SSSPKLRRNAHSRLESYRPDTDLSREDTGCN | LES(1)YRPDTDLSR | 1458100 |
| 1405 | Q9CZW5 | 108 | Mitochondrial import receptor subunit TOM70 | Tomm70a | ASPALGSGHHDGSGDSLEMSSLDRAQAAKNK | ASPALGS(0.018)GHHDGS(0.219)GDS(0.755)LEMS(0.006)S(0.002)LDR | 0 |
| 723 | Q9QYR6 | 667 | crotubule-associated protein 1A;MAP1A heavy chain;MAP1 light chain L | Map1a | VIEKAELEEMEEVHPSDEEEEETKAESFYQK | AELEEMEEVHPS(0.992)DEEEEET(0.008)KAESFYQK | 0 |
| 715 | P0C192 | 700 | Leucine-rich repeat-containing protein 4B | Lrrc4b | PGLNSIHEPLLFKSGSKENVQETQI______ | S(0.5)GS(0.5)KENVQETQI | 7072500 |
| 714 | P0C192 | 698 | Leucine-rich repeat-containing protein 4B | Lrrc4b | KGPGLNSIHEPLLFKSGSKENVQETQI____ | S(0.586)GS(0.414)KENVQETQI | 7072500 |
| 269 | P97445 | 2303 | Voltage-dependent P/Q-type calcium channel subunit alpha-1A | Cacna1a | PEGPRPRGADYTEPDSPREPPGGAHDPAPRS | GADY(0.002)T(0.027)EPDS(0.971)PREPPGGAHDPAPR | 3627300 |
| 391 | P97821 | 193 | peptidase 1 exclusion domain chain;Dipeptidyl peptidase 1 heavy chain; | Ctsc | INTVQKSWTATAYKEYEKMSLRDLIRRSGHS | EY(1)EKMS(1)LRDLIR | 5290800 |
| 390 | P97821 | 197 | peptidase 1 exclusion domain chain;Dipeptidyl peptidase 1 heavy chain; | Ctsc | QKSWTATAYKEYEKMSLRDLIRRSGHSQRIP | EY(1)EKMS(1)LRDLIR | 5290800 |
| 185 | Q9R0K7 | 1117 | Plasma membrane calcium-transporting ATPase 2 | Atp2b2 | FRSSLYEGLEKPESRTSIHNFMAHPEFRIED | T(0.5)S(0.5)IHNFMAHPEFR | 23659000 |
| 183 | Q9R0K7 | 1118 | Plasma membrane calcium-transporting ATPase 2 | Atp2b2 | RSSLYEGLEKPESRTSIHNFMAHPEFRIEDS | T(0.5)S(0.5)IHNFMAHPEFR | 23659000 |
| 905 | P13595 | 1005 | Neural cell adhesion molecule 1 | Ncam1 | PGTVKNPPEAATAPASPKSKAATTNPSQGED | NPPEAATAPAS(1)PK | 46349000 |
| 708 | Q7TME0 | 647 | Lipid phosphate phosphatase-related protein type 4 | Lppr4 | NRPIIQIPSSTEGEGSGSWKWKAPEKSSLRQ | VEAHPENNRPIIQIPS(0.008)S(0.008)T(0.008)EGEGS(0.525)GS(0.451)WK | 0 |
| 278 | Q61290 | 873 | Voltage-dependent R-type calcium channel subunit alpha-1E | Cacna1e | KGDIGGLTSALDNQRSPLSLGKREPPWLPRS | GGS(1)LKGDIGGLT(0.001)S(0.001)ALDNQRS(0.981)PLS(0.017)LGK | 0 |
| 18 | Q3UHJ0 | 729 | AP2-associated protein kinase 1 | Aak1 | FAKLGEGKLPEKLGGSAESLIPGFQPTQGDA | LGGS(0.809)AES(0.191)LIPGFQPTQGDAFTTPSFSAGTAEK | 0 |
| 107 | Q8C8R3 | 3686 | Ankyrin-2 | Ank2 | GYSTFQDCLPKTEGDSPAAALSPQMHQEPVQ | T(0.004)EGDS(0.995)PAAALS(0.001)PQMHQEPVQQDFSGK | 0 |
| 282 | Q61290 | 29 | Voltage-dependent R-type calcium channel subunit alpha-1E | Cacna1e | GSGDGDSDQSRNRQGTPVPASGPAAAYKQSK | QGT(1)PVPASGPAAAYK | 0 |
| 1329 | Q9JIS5 | 127 | Synaptic vesicle glycoprotein 2A | Sv2a | RMADGAPLAGVRGGLSDGEGPPGGRGEAQRR | GGLS(1)DGEGPPGGRGEAQR | 0 |
| 1301 | O35526 | 14 | Syntaxin-1A | Stx1a | __MKDRTQELRTAKDSDDDDDVTVTVDRDRF | TAKDS(1)DDDDVTVTVDR | 3209300 |
| 765 | P14873 | 829 | crotubule-associated protein 1B;MAP1B heavy chain;MAP1 light chain L | Map1b | SGPVKELEAERSLMSSPEDLTKDFEELKAEE | S(0.032)LMS(0.977)S(0.986)PEDLT(0.005)KDFEELKAEEIDVAK | 0 |
| 764 | P14873 | 828 | crotubule-associated protein 1B;MAP1B heavy chain;MAP1 light chain L | Map1b | ASGPVKELEAERSLMSSPEDLTKDFEELKAE | S(0.032)LMS(0.977)S(0.986)PEDLT(0.005)KDFEELKAEEIDVAK | 0 |
| 76 | Q7TQF7 | 500 | Amphiphysin | Amph | EAEAEKAALPAGEGGSPEGAKIDGESTELAI | AALPAGEGGS(1)PEGAK | 2770900 |
| 197 | Q80ZF8 | 1220 | Brain-specific angiogenesis inhibitor 3 | Bai3 | GPCRAATITGTLSRISLNDDEEEKGTNPEGL | IS(0.891)LNDDEEEKGT(0.105)NPEGLS(0.002)Y(0.001)STLPGNVISK | 6208100 |
| 1413 | E9Q6P5 | 160 | | Ttc7b | YATKGLCLEKLPVSSSTSNLHVDREQDVITC | LPVSS(0.106)S(0.736)T(0.131)S(0.028)NLHVDR | 0 |
| 393 | Q6WQJ1 | 808 | Sn1-specific diacylglycerol lipase alpha | Dagla | SFDSRRSSGFRSIRGSPSLHAVLERDEGHLF | GS(0.992)PS(0.008)LHAVLER | 0 |
| 1192 | Q14BI1 | 430 | | Slc24a2 | HTLDPLAEGRFREKASILHKIAKKKCQVDEN | EKAS(1)ILHK | 6149000 |
| 345 | Q9ERZ4 | 232 | Muscarinic acetylcholine receptor M2 | Chrm2 | IKKEKKEPVANQDPVSPSLVQGRIVKPNNNN | EKKEPVANQDPVS(0.9)PS(0.1)LVQGR | 0 |
| 373 | Q61301 | 640 | Catenin alpha-2 | Ctnna2 | KAVLMIRTPEELEDDSDFEQEDYDVRSRTSV | TPEELEDDS(1)DFEQEDYDVR | 0 |
| 1087 | O54829 | 247 | Regulator of G-protein signaling 7 | Rgs7 | GLQNDIRSHSPTHTPTPETKPPTEDELHQQI | S(0.144)HS(0.783)PT(0.078)HT(0.088)PT(0.901)PET(0.005)KPPTEDELH | 6844700 |
| 1054 | Q2PFD7 | 759 | PH and SEC7 domain-containing protein 3 | Psd3 | EKANGTHPKTISRIGSTTNPFLDIPHDPNAA | IGS(0.723)T(0.131)T(0.145)NPFLDIPHDPNAAVYK | 5860300 |
| 1066 | B9EKR1 | 572 | Receptor-type tyrosine-protein phosphatase zeta | Ptprz1 | AESLNTVPITEYKEVSADVSEEENFLTDFKL | EVS(0.792)ADVS(0.208)EEENFLTDFK | 0 |
| 410 | Q91XM9 | 65 | Disks large homolog 2 | Dlg2 | SQIENVHGYVLQSHISPLKASPAPIIVNTDT | NLSQIENVHGYVLQS(0.017)HIS(0.983)PLK | 2275200 |
| 95 | Q8C8R3 | 1889 | Ankyrin-2 | Ank2 | PPVPSSGRTEKHPPVSPGKTEKHLPGSPSIR | HPPVS(1)PGKTEK | 11867000 |
| 1256 | Q3UTJ2 | 829 | Sorbin and SH3 domain-containing protein 2 | Sorbs2 | LKDICDQIKAEKRRGSLPDNSILHRLISELL | RGS(1)LPDNSILHR | 2764600 |
| 820 | P20357 | 1609 | Microtubule-associated protein 2 | Map2 | TSTPTTPGSTAITPGTPPSYSSRTPGTPGTP | SGTSTPTTPGSTAIT(0.039)PGT(0.955)PPS(0.002)YS(0.001)S(0.001)R | 5944200 |
| 1339 | O88935 | 427 | Synapsin-1 | Syn1 | NKMTQALPRQPQRDASPGRGSHSQSSSPGAL | DAS(1)PGRGSHSQSSSPGALTLGR | 229590000 |
| 138 | P84091 | 156 | AP-2 complex subunit mu | Ap2m1 | QHQTKEEQSQITSQVTGQIGWRREGIKYRRN | EEQSQITSQVT(1)GQIGWR | 0 |
| 1160 | Q4ACU6 | 781 | SH3 and multiple ankyrin repeat domains protein 3 | Shank3 | EKLPGSLRKGIPRTKSVGEDEKLASLLEGRF | T(0.025)KS(0.969)VGEDEKLAS(0.007)LLEGR | 6537400 |
| 159 | Q9D5T0 | 322 | ATPase family AAA domain-containing protein 1 | Atad1 | ALLCVREYVNSTSEESHDEDEIRPVQQQDLH | EYVNST(0.001)S(0.003)EES(0.996)HDEDEIRPVQQQDLHR | 0 |
| 981 | Q01063 | 515 | cAMP-specific 3',5'-cyclic phosphodiesterase 4D | Pde4d | SLRKMVIDIVLATDMSKHMNLLADLKTMVET | MVIDIVLAT(1)DMS(1)K | 0 |
| 199 | Q8BKX1 | 326 | Brain-specific angiogenesis inhibitor 1-associated protein 2 | Baiap2 | YNPWADRKAAQPKSLSPPQSQSKLSDSYSNT | SLS(1)PPQSQSK | 6873600 |
| 1294 | P54227 | 38 | Stathmin | Stmn1 | ILSPRSKESVPDFPLSPPKKKDLSLEEIQKK | SKESVPDFPLS(1)PPK | 0 |
| 1205 | Q6PSF6 | 593 | Zinc transporter ZIP10 | Slc39a10 | RLNETELTDLEAQQESPPKNYLGVEEEKIMD | LNETELTDLEAQQES(1)PPKNYLGVEEEK | 0 |
| 689 | Q6P9L6 | 622 | Kinesin-like protein KIF15 | Kif15 | ELTRKKQLELESELQSLQKANLNLENLLEAT | KKQLELES(0.5)LQS(0.5)LQK | 10872000 |
| 688 | Q6P9L6 | 618 | Kinesin-like protein KIF15 | Kif15 | EEFKELTRKKQLELESELQSLQKANLNLENL | KKQLELES(0.5)LQS(0.5)LQK | 10872000 |
| 1249 | P60879 | 29 | Synaptosomal-associated protein 25 | Snap25 | MQRRADQLADESLESTRRMLQLVEESKDAGI | RADQLADES(0.5)T(0.5)RR | 0 |
| 1248 | P60879 | 28 | Synaptosomal-associated protein 25 | Snap25 | EMQRRADQLADESLESTRRMLQLVEESKDAG | RADQLADES(0.001)LES(0.579)T(0.419)RR | 0 |
| 261 | Q80XB0 | 468 | C2 domain-containing protein 2-like | C2cd2l | QSRPRVDGKLDSPSRSPSKVEVTEKMTTVLS | LDS(0.994)PS(0.013)RS(0.895)PS(0.098)KVEVTEK | 2217800 |
| 1243 | Q61165 | 790 | Sodium/hydrogen exchanger 1 | Slc9a1 | SSPGTDDVFTPGSSDSPSSQRIQRCLSDPGP | SKEPSSPGTDDVFTPGS(0.006)DS(0.741)PS(0.209)S(0.045)QR | 8891100 |
| 146 | P55088 | 285 | Aquaporin-4 | Aqp4 | AQQTKGSYMEVEDNRSQVETEDLILKPGVVH | S(1)QVETEDLILKPGVVHVIDIDR | 3169600 |
| 519 | Q80T41 | 818 | Gamma-aminobutyric acid type B receptor subunit 2 | Gabbr2 | ELDKDLEEVTMQLQDTPEKTTYIKQNHYQEL | ITELDKDLEEVTMQLQDT(1)PEK | 1454500 |
| 442 | O70251 | 106 | Elongation factor 1-beta | Eef1b | GAADAKDDDDIDLFGSDDEEESEEAKKLREE | YGPSSVEDTTGSGAADAKDDDDIDLFGS(1)DDEEESEEAKK | 0 |
| 1272 | Q62261 | 2318 | Spectrin beta chain, non-erythrocytic 1 | Sptbn1 | SAISSDKHDTSASTQSTPASSRAQTLPTSVV | DDEEMNTWIQAS(0.006)S(0.006)AS(0.058)S(0.061)DKHDT(0.22)S(0.2 | 2016400 |
| 851 | P10637 | 523 | Microtubule-associated protein tau | Mapt | PTPTREPKKVAVVRTPPKSPSASKSRLQTA | VAVVRT(0.999)PPKS(0.802)PS(0.171)AS(0.028)K | 8640100 |
| 846 | P10637 | 527 | Microtubule-associated protein tau | Mapt | TREPKKVAVVRTPPKSPSASKSRLQTAPVPM | VAVVRT(0.998)PPKS(0.848)PS(0.126)AS(0.028)K | 8640100 |
| 1355 | O88935 | 684 | Synapsin-1 | Syn1 | NAFNLPEPAPPRPSLSQDEVKAETIRSLRKS | S(0.113)QS(0.816)LT(0.071)NAFNLPEPAPPRPS(0.051)LS(0.949)QDEVK | 0 |
| 1068 | Q9JKF6 | 421 | Nectin-1 | Pvrl1 | QHHPPMAQNLQYPDDSDDEKKAGPLGGSSYE | AGIPQHHPPMAQNLQYPDDS(1)DDEKK | 10130000 |
| 1115 | Q99P72 | 857 | Reticulon-4 | Rtn4 | EIIDEFPTFVSAKDDSPKEYTDLEVSNKSEI | DDS(1)PKEYTDLEVSNK | 0 |
| 539 | P23242 | 372 | Gap junction alpha-1 protein | Gja1 | AIVDQRPSSRASSRASSRPRPDDLEI_____ | AS(0.533)S(0.467)RPRPDDLEI | 0 |
| 1122 | Q99P72 | 167 | Reticulon-4 | Rtn4 | APPSTPAAPKRRGSGSVDETLFALPAASEPV | RRGS(0.358)GS(0.625)VDET(0.018)LFALPAASEPVIPSSAEK | 0 |
| 780 | P20357 | 654 | Microtubule-associated protein 2 | Map2 | QSYTPDHPSELPEEPSSPQERMFTIDPKVYG | AS(0.434)QPS(0.562)PPAQEAGY(0.004)STLAQSYTPDHPS(0.006)ELPEEPS | 20676000 |
| 1103 | P99027;A0A5F8MPY2 | 105;105 | 60S acidic ribosomal protein P2 | Rplp2 | AAEEKKDEKKEESEESDDDMGFGLFD_____ | KEES(1)EES(1)DDDMGFGLFD | 0 |
| 1102 | P99027;A0A5F8MPY2 | 102;102 | 60S acidic ribosomal protein P2 | Rplp2 | APAAAEEKKDEKKEESEESDDDMGFGLFD__ | KEES(1)EES(1)DDDMGFGLFD | 0 |
| 294 | O35887 | 47 | Calumenin | Calu | PQLSDKVHNDAQNFDYDHDAFLGAEEAKSFD | VHNDAQNFDY(1)DHDAFLGAEEAK | 0 |
| 458 | O70318 | 543 | Band 4.1-like protein 2 | Epb41l2 | PAPQFERASSKRVSRSLDGAPIGVVDQSPPG | S(0.736)LDGAPIGVVDQS(0.221)PPGEGS(0.043)VPGPGVISYTTIQDGR | 0 |
| 1315 | Q64324 | 48 | Syntaxin-binding protein 2 | Stxbp2 | MDHPSMRILSSCCKMSDILAEGITIVEDINK | MS(0.951)DILAEGIT(0.049)IVEDINKR | 4334800 |
| 299 | Q3UHL1 | 446 | CaM kinase-like vesicle-associated protein | Camkv | PATDRSATPATDGRATPATEESTVPATQSSA | S(0.2)AT(0.787)PAT(0.074)DGRAT(0.927)PAT(0.011)EESTVPATQSSALPA | 0 |
| 705 | Q7TME0 | 547 | Lipid phosphate phosphatase-related protein type 4 | Lppr4 | SSQLVHIPEETQENISTSPKSSSARAKWLKA | PGSSQLVHIPEETQENIS(0.605)T(0.258)S(0.137)PK | 4750000 |
| 1300 | O35526 | 64 | Syntaxin-1A | Stx1a | ENVEEVKRKHSAILASPNPDEKTKEELEELM | HSAILAS(1)PNPDEK | 11268000 |
| 302 | Q3UHL1 | 438 | CaM kinase-like vesicle-associated protein | Camkv | PATDGSVTPATDRSATPATDGRATPATEEST | S(0.2)AT(0.787)PAT(0.074)DGRAT(0.927)PAT(0.011)EESTVPATQSSALPA | 15963000 |
| 1038 | Q6PE13 | 799 | Proline-rich transmembrane protein 3 | Prrt3 | VGPAPSLSELDLRPPSPINLSRSIDAALFRE | GSVGPAPS(0.807)LS(0.196)ELDLRPPS(0.969)PINLS(0.028)R | 2126900 |
| 1055 | Q2PFD7 | 998 | PH and SEC7 domain-containing protein 3 | Psd3 | ELLTTDGNEPVGLKKSHSSPSLNPDASPVTA | KS(0.597)HS(0.281)S(0.092)PS(0.031)LNPDASPVTAK | 13000000 |
| 464 | Q9WV92 | 63 | Band 4.1-like protein 3;Band 4.1-like protein 3, N-terminally processed | Epb41l3 | KQQPALEQFPEAAAHSTPVKREIGDKDRDFA | QQPALEQFPEAAAHS(0.611)T(0.389)PVKR | 11565000 |
| 1235 | P63115 | 5 | Asc-type amino acid transporter 1 | Slc7a10 | __________MRRDSDMASHIQQPGGHGNP | RDS(0.937)DMAS(0.062)HIQQPGGHGNPGPAPSPSPGPGPGPGASER | 5247800 |
| 300 | Q3UHL1 | 449 | CaM kinase-like vesicle-associated protein | Camkv | DRSATPATDGRATPATEESTVPATQSSALPA | AT(0.069)PAT(0.931)EESTVPATQSSALPAAK | 0 |
| 133 | Q8C8R3 | 3718 | Ankyrin-2 | Ank2 | DFSGKTQDQQEYYVTTPGAEVEDPQKATAVP | TQDQQEY(0.002)Y(0.002)VT(0.046)T(0.95)PGAEVEDPQK | 0 |
| 75 | Q3TZM9 | 181 | GDP-Man:Man(3)GlcNAc(2)-PP-Dol alpha-1,2-mannosyltransferase | Alg11 | QRVPDVYIDSMGYAFTLPLFKYVGGCRVGSY | VPDVYIDS(0.001)MGY(0.468)AFT(0.531)LPLFK | 0 |
| 833 | P10637 | 692 | Microtubule-associated protein tau | Mapt | KTDHGAEIVYKSPVVSGDTSPRHLSNVSSTG | AKTDHGAEIVYKS(1)PVVS(1)GDT(0.329)S(0.671)PR | 0 |
| 528 | P06837 | 142 | Neuromodulin | Gap43 | GSAETESAAKATTDNSPSSKAEDGPAKEEPK | ATTDNS(0.999)PSSKAEDGPAKEEPK | 230930000 |
| 303 | Q3UHL1 | 441 | CaM kinase-like vesicle-associated protein | Camkv | DGSVTPATDRSATPATDGRATPATEESTVPA | S(0.496)AT(0.522)PAT(0.569)DGRAT(0.364)PAT(0.032)EES(0.009)T(0.0 | 15963000 |
| 1030 | E9PUL5 | 72 | Proline-rich transmembrane protein 2 | Prrt2 | APVDSGPKAELAPETTETPVETPETVQATDL | AELAPET(0.159)T(0.64)ET(0.097)PVET(0.097)PET(0.008)VQATDLSLNPEE | 0 |
| 1120 | Q99P72 | 839 | Reticulon-4 | Rtn4 | SKEDKMKESETFSDSSPIEIIDEFPTFVSAK | MKES(0.019)ET(0.05)FS(0.523)DS(0.714)S(0.694)PIEIIDEFPTFVSAK | 0 |
| 454 | Q9Z2H5 | 685 | Band 4.1-like protein 1 | Epb41l1 | ESGGLEDSPDRGACSTPEMPQFESVKAETMT | GACS(0.073)T(0.927)PEMPQFESVK | 0 |
| 1331 | Q8BG39 | 33 | Synaptic vesicle glycoprotein 2B | Sv2b | GYRSNEQNQEEDAQSDVTEGHDEEDEIYEG | SNEQNQEEDAQS(0.974)DVT(0.026)EGHDEEDEIYEGEYQGIPHPDDVK | 0 |
| 471 | Q9WV92 | 64 | Band 4.1-like protein 3;Band 4.1-like protein 3, N-terminally processed | Epb41l3 | QQPALEQFPEAAAHSTPVKREIGDKDRDFAA | QQPALEQFPEAAAHS(0.142)T(0.858)PVKR | 14806000 |
| 1229 | Q8BJI1 | 13 | Sodium-dependent neutral amino acid transporter SLC6A17 | Slc6a17 | ___MPKNSKVTQREHSNEHVTESVADLLALE | EHS(0.993)NEHVT(0.007)ESVADLLALEEPVDYK | 0 |
| 785 | P20357 | 136 | Microtubule-associated protein 2 | Map2 | AALPLAAEETANLPPSPPPSPASEQTATVEE | DQPAALPLAAEET(0.005)ANLPPS(0.992)PPPS(0.834)PAS(0.166)EQT(0.0 | 0 |
| 533 | P06837 | 171 | Neuromodulin | Gap43 | PKQADVPAAVTDAAATTPAAEDAATKAAQPP | QADVPAAVTDAAAT(0.82)T(0.18)PAAEDAATK | 67657000 |
| 952 | Q9Z0P4 | 141 | Paralemmin-1 | Palm | AKEEQKSETLVNAQQTPLGTPKENRTSTPVR | SETLVNAQQT(1)PLGT(1)PK | 1915100 |
| 919 | P08553 | 610 | Neurofilament medium polypeptide | Nefm | IKVEKPEKAKSPMPKSPVEEVKPKPEAKAGK | S(1)PVEEVKPKPEAK | 12433000 |
| 180 | Q9R0K7 | 1155 | Plasma membrane calcium-transporting ATPase 2 | Atp2b2 | IDDTDLEEDAALKQNSSPPSSLNKNNSAIDS | IEDSQPHIPLIDDTDLEEDAALKQNS(0.603)S(0.395)PPS(0.002)SLNK | 0 |
| 340 | Q9JKC6 | 9 | Cell cycle exit and neuronal differentiation protein 1 | Cend1 | ______MESRGKSASSPKPDTKVPQATAEA | SAS(0.599)S(0.401)PKPDTKVPQATAEAK | 0 |
| 1178 | P43006 | 530 | Excitatory amino acid transporter 2 | Slc1a2 | MTKTQSIYDDKNHRESNSNQCVYAAHNSVVI | TQS(1)YDDKNHRES(0.994)NS(0.006)NQCVYAAHNSVVIDECK | 0 |
| 425 | P60904 | 15 | DnaJ homolog subfamily C member 5 | Dnajc5 | _MADQRQRSLSTSGESLYHVLGLDKNATSDD | S(0.458)LS(0.518)T(0.026)S(0.041)GES(0.951)LY(0.006)HVLGLDK | 0 |
| 231 | O08539 | 298 | Myc box-dependent-interacting protein 1 | Bin1 | AQPSDNAPEKGNKSPSPPPDGSPAATPEIRV | GNKS(1)PS(1)PPPDGSPAATPEIR | 0 |
| 230 | O08539 | 296 | Myc box-dependent-interacting protein 1 | Bin1 | VKAQPSDNAPEKGNKSPSPPPDGSPAATPEI | GNKS(1)PS(1)PPPDGSPAATPEIR | 0 |
| 407 | Q9JLM8 | 392 | Serine/threonine-protein kinase DCLK1 | Dclk1 | LGRRHSLQRGWREESEEGFQIPATITERYK | REES(1)EEGFQIPATITER | 0 |
| 927 | Q8BYM5 | 722 | Neuroligin-3 | Nlgn3 | YYRKDKRRQEPLRQPSPQRGTGAPELGTAPE | RQEPLRQPS(1)PQR | 21982000 |
| 562 | Q8C419 | 944 | Probable G-protein coupled receptor 158 | Gpr158 | AKSQKPKDRETIRKYSNSDNVETIPNSGHME | KYS(0.714)NS(0.286)DNVETIPNSGHMEEPR | 7235300 |
| 592 | Q8BWS5 | 327 | G protein-regulated inducer of neurite outgrowth 3 | Gprin3 | EVQAVASVESRSVSTSPSILPAYLKENPAPE | S(0.933)VS(0.059)T(0.06)S(0.909)PS(0.04)ILPAYLK | 0 |
| 591 | Q8BWS5 | 323 | G protein-regulated inducer of neurite outgrowth 3 | Gprin3 | WQDAEVQAVASVESRSVSTSPSILPAYLKEN | S(0.933)VS(0.059)T(0.06)S(0.909)PS(0.04)ILPAYLK | 0 |
| 1203 | O35633 | 97 | Vesicular inhibitory amino acid transporter | Slc32a1 | DIHYQRGGAPLPPSGSKDQAVGAGGEFGGHD | GGAPLPPS(0.5)GS(0.5)KDQAVGAGGEFGGHDKPK | 5442800 |
| 1202 | O35633 | 95 | Vesicular inhibitory amino acid transporter | Slc32a1 | EGDIHYQRGGAPLPPSGSKDQAVGAGGEFGG | GGAPLPPS(0.5)GS(0.5)KDQAVGAGGEFGGHDKPK | 5442800 |
| 467 | Q9WV92 | 547 | Band 4.1-like protein 3;Band 4.1-like protein 3, N-terminally processed | Epb41l3 | ERTDTAADGETSATESDQEEDAEIKAQDLDK | TDTAADGET(0.486)S(0.514)AT(0.333)ES(0.667)DQEEDAEIKAQDLDKTQD | 0 |
| 466 | Q9WV92 | 543 | Band 4.1-like protein 3;Band 4.1-like protein 3, N-terminally processed | Epb41l3 | KTDSERTDTAADGETSATESDQEEDAEIKAQ | TDTAADGET(0.486)S(0.514)AT(0.333)ES(0.667)DQEEDAEIKAQDLDKTQD | 0 |
| 1359 | Q64332 | 546 | Synapsin-2 | Syn2 | PAQKPQPHPQLNKSQSLTNAFSFSESSFFRS | S(0.007)QS(0.993)LTNAFS(0.993)FS(0.007)ESSFFR | 5154600 |
| 1310 | O08599 | 594 | Syntaxin-binding protein 1 | Stxbp1 | LDTLKKLNKTDEEISS______________ | KLNKTDEEIS(0.027)S(0.973) | 0 |
| 1258 | P70663 | 155 | SPARC-like protein 1 | Sparcl1 | YVDPNQRANIITKGKESQEQPVSDSHQQPNES | GKES(0.992)QEQPVS(0.007)DSHQQPNESSK | 2506700 |
| 78 | Q7TQF7 | 272 | Amphiphysin | Amph | IAKTPSPEEPSPLPSPTASPNHTLAPASPA | T(0.485)PS(0.511)PEEPS(0.038)PLPS(0.841)PT(0.121)AS(0.004)PNHTL | 2507500 |
| 77 | Q7TQF7 | 262 | Amphiphysin | Amph | GAPSDSGPLRIAKTPSPEEPSPLPSPTASP | T(0.485)PS(0.511)PEEPS(0.038)PLPS(0.841)PT(0.121)AS(0.004)PNHTL | 2507500 |
| 571 | Q8C419 | 1063 | Probable G-protein coupled receptor 158 | Gpr158 | ATEGLYQANHKSIDKTEVCPWEIHSQSLLED | S(0.5)IDKT(0.5)EVCPWEIHSQSLLEDENR | 0 |
| 901 | P13595 | 770 | Neural cell adhesion molecule 1 | Ncam1 | PGAKGKDMEEGKAAFSKDESKEPIVEVRTEE | AAFS(1)KDES(1)KEPIVEVR | 6862400 |
| 1325 | Q8K400 | 786 | Syntaxin-binding protein 5 | Stxbp5 | VKDNSFSRSRSSSVTSIDKESRETISALHFC | S(0.1)S(0.131)S(0.768)VT(0.009)S(0.99)IDKESR | 3443500 |
| 271 | P97445 | 433 | Voltage-dependent P/Q-type calcium channel subunit alpha-1A | Cacna1a | PFDGALRRATLKKSKTDLLNPEEAEDQLADI | S(0.465)KT(0.535)DLLNPEEAEDQLADIASVGSPFAR | 0 |
| 1409 | Q7TQD2 | 15 | Tubulin polymerization-promoting protein | Tppp | _MADSKAKPAKAANKTPPKSPGDPARAAKRL | AANKT(1)PPKS(1)PGDPAR | 74325000 |
| 291 | Q9JJV5 | 302 | Voltage-dependent calcium channel gamma-3 subunit | Cacng3 | AFLQFHNSTPKEFKESLHNNPANRRTTPV__ | EFKES(1)LHNNPANR | 5417600 |
| 1308 | Q9JKK1 | 2 | Syntaxin-6 | Stx6 | _____________MSMEDPFFVVKGEVQKA | S(1)MEDPFFVVK | 0 |
| 934 | Q9D8N4 | 183 | Putative monooxygenase p33MONOX | P33monox | HPASAQSTPSSTPHASPKQKSRGWFPSGSST | HPASAQSTPS(0.004)S(0.021)T(0.021)PHAS(0.952)PK | 4049600 |
| 1183 | P43006 | 520 | Excitatory amino acid transporter 2 | Slc1a2 | SQHRMQEHEMTKTQSIYDDKNHRESNSNQC | TQS(1)YDDKNHR | 23881000 |
| 235 | O55003 | 60 | BCL2/adenovirus E1B 19 kDa protein-interacting protein 3 | Bnip3 | QHESGRSSSKSSHCDSPPRSQTPQDTNRAEI | S(0.007)S(0.007)HCDS(0.87)PPRS(0.058)QT(0.058)PQDT(0.002)NR | 2910700 |
| 1131 | Q99P72 | 488 | Reticulon-4 | Rtn4 | EKKIEEKKAQIITEKTSPKTSNPFLVAIHDS | AQIIT(0.217)EKT(0.511)S(0.272)PK | 7690200 |
| 1010 | P05132;P68181 | 196;196 | rotein kinase catalytic subunit alpha;cAMP-dependent protein kinase cat | Prkaca;Prkacb | IQVTDFGFAKRVKGRTWTLCGTPEYLAPEII | T(0.525)WT(0.463)LCGT(0.012)PEYLAPEIILSK | 2970000 |
| 444 | Q9D8Y0 | 74 | EF-hand domain-containing protein D2 | Efhd2 | LLRRADLNQGIGEPQSPSRRVFNPYTEFKEF | RADLNQGIGEPQS(0.963)PS(0.037)RR | 0 |
| 53 | Q9QYB8 | 618 | Beta-adducin | Add2 | PVQSPSKAGTKSPAVSPSKTSEDTKKTEVSE | AGT(0.972)KS(0.028)PAVS(0.978)PS(0.019)KT(0.002)SEDTK | 0 |
| 52 | Q9QYB8 | 614 | Beta-adducin | Add2 | TPASPVQSPSKAGTKSPAVSPSKTSEDTKKT | AGT(0.031)KS(0.969)PAVS(0.937)PS(0.06)KT(0.003)SEDTK | 0 |
| 457 | Q9Z2H5 | 475 | Band 4.1-like protein 1 | Epb41l1 | ESGGRRSEAEEGEVRTPTKIKELKPEQETTP | SEAEEGEVRT(0.976)PT(0.024)K | 10859000 |
| 836 | P10637 | 704 | Microtubule-associated protein tau | Mapt | PVVSGDTSPRHLSNVSSTGSIDMVDSPQLAT | HLS(0.174)NVS(0.665)S(0.687)T(0.285)GS(0.188)IDMVDS(0.001)PQLAT | 0 |
| 885 | Q80WJ7 | 565 | Protein LYRIC | Mtdh | NSVPPSQTKSETNWESPKQIKKKKKARRET_ | SETNWES(1)PKQIK | 3206500 |
| 811 | P20357 | 146 | Microtubule-associated protein 2 | Map2 | ANLPPSPPPSPASEQTATVEEDLLTASKMEF | DQPAALPLAAEET(0.045)ANLPPS(0.467)PPPS(0.467)PAS(0.467)EQT(0.5 | 3219500 |
| 703 | Q7TPB0 | 346 | Lipid phosphate phosphatase-related protein type 3 | Lppr3 | EGVPRPVAREKTSLGSLKRASVDVDLLAPRS | EKT(0.003)S(0.022)LGS(0.974)LKR | 11734000 |
| 974 | Q9QYX7 | 4774 | Protein piccolo | Pclo | SIEHGKSHSSQNSQQSPKPSVIKSRSHGIFP | SHS(0.004)S(0.026)QNS(0.14)QQS(0.8)PKPS(0.029)VIK | 4884200 |
| 135 | Q8C8R3 | 3363 | Ankyrin-2 | Ank2 | SSLDAKTKCPVKARSYIETETESRERAEGFE | ARS(0.475)Y(0.525)IETETESR | 3807800 |
| 1009 | P05132 | 339 | cAMP-dependent protein kinase catalytic subunit alpha | Prkaca | DTSNFDDYEEEEIRVSINEKCGKEFTEF___ | FKGPGDTSNFDDYEEEEIRVS(1)INEK | 14534000 |
| 560 | P35803 | 318 | Neuronal membrane glycoprotein M6-b | Gpm6b | QDIKAKEEQELQDIQSRSKEQLNSYT_____ | AKEEQELQDIQS(1)R | 12096000 |
| 670 | Q6WVG3 | 153 | BTB/POZ domain-containing protein KCTD12 | Kctd12 | GPPPPHSRRGVHKEGSLGDELLPLGYAEPEP | EGS(1)LGDELLPLGYAEPEPQEGAS(0.018)AGAPS(0.823)PT(0.15)LELAS(0.0 | 0 |
| 1209 | Q5DTL9 | 1115 | Sodium-driven chloride bicarbonate exchanger | Slc4a10 | TADNSKEKESRFPSKSSPS_____________ | ESRFPSKS(0.885)S(0.115)PS | 3555400 |
| 1193 | Q14BI1 | 396 | | Slc24a2 | DGPTLPSKPRLQRGGSSASLHNSLMRNSIFQ | GGS(0.979)S(0.021)ASLHNSLMR | 7625300 |
| 659 | Q9ER47 | 174 | Potassium voltage-gated channel subfamily H member 7 | Kcnh7 | GFKFPGLRVLTYRKQSLPQEDPDVVVIDSSK | KQS(1)LPQEDPDVVVIDSSK | 3961600 |
| 1007 | H3BJD6 | 160 | | Ppp1r9a | MFERSVHESGQNHRYSPKKEKGGGTEPQDEW | SVHES(0.039)GQNHRYS(0.961)PK | 3286500 |
| 967 | Q9QYX7 | 1763 | Protein piccolo | Pclo | ELRAQRRRERPKTPPSNLSPIEDASPTEELR | ERPKT(0.977)PPS(0.512)NLS(0.512)PIEDAS(0.815)PT(0.185)EELR | 0 |
| 565 | Q8C419 | 869 | Probable G-protein coupled receptor 158 | Gpr158 | SKKLTQKLKEDSEAESTESVPLVCKSASAHN | LKEDS(0.996)EAES(0.799)T(0.192)ES(0.013)VPLVCK | 4028900 |
| 671 | Q6WVG3 | 178 | BTB/POZ domain-containing protein KCTD12 | Kctd12 | YAEPEPQEGASAGAPSPTLELASRSPSGGAA | EGS(1)LGDELLPLGYAEPEPQEGAS(0.018)AGAPS(0.823)PT(0.15)LELAS(0.0 | 0 |
| 616 | O88866 | 344 | Hormonally up-regulated neu tumor-associated kinase | Hunk | PCNVTYPNRISLEDLSPSVVLHMTEKLGYKN | IS(0.001)LEDLS(0.845)PS(0.077)VVLHMT(0.077)EK | 3759400 |
| 991 | O70161 | 552 | Phosphatidylinositol 4-phosphate 5-kinase type-1 gamma | Pip5k1c | RSPSDTSEQPRYRRRTQSSGQDGRPQEEPHA | RT(0.521)QS(0.409)S(0.07)GQDGRPQEEPHAEDLQK | 0 |
| 445 | Q6ZQ18 | 689 | Protein EFR3 homolog B | Efr3b | LCLPYIPQLTDEDRLSKRKSIGETISLQVEV | LCLPYIPQLT(0.192)DEDRLS(0.807)K | 0 |
| 63 | Q3TY86 | 36 | Apoptosis-inducing factor 3 | Aifm3 | EKERGKEELSASGKGSPRGYQGNGTARHFHA | GKEELSAS(0.025)GKGS(0.975)PR | 2871400 |
| 561 | P35803 | 320 | Neuronal membrane glycoprotein M6-b | Gpm6b | IKAKEEQELQDIQSRSKEQLNSYT______ | S(1)KEQLNSYT | 70347000 |
| 462 | Q9WV92 | 704 | Band 4.1-like protein 3;Band 4.1-like protein 3, N-terminally processed | Epb41l3 | KVLQETILVEERHVMSVHASGDASHTARDEV | HVMS(0.998)VHAS(0.001)GDASHTAR | 0 |
| 1059 | Q2PFD7 | 1009 | PH and SEC7 domain-containing protein 3 | Psd3 | GLKKSHSSPSLNPDASPVTAKVKRNVSERKD | SHS(0.153)S(0.847)PSLNPDAS(1)PVTAK | 2151100 |
| 576 | Q3UNH4 | 693 | G protein-regulated inducer of neurite outgrowth 1 | Gprin1 | TSQKDLAAAAAQKSPSAEAAAPPPGPRTRDN | S(0.256)PS(0.744)AEAAAPPPGPR | 0 |
| 468 | Q9WV92 | 488 | Band 4.1-like protein 3;Band 4.1-like protein 3, N-terminally processed | Epb41l3 | EVGTGQYATTKGISQTNLITTVTPEKKAEEE | GIS(0.438)QT(0.561)NLIT(0.001)T(0.001)VTPEK | 4000600 |
| 690 | Q9QXL2 | 855 | Kinesin-like protein KIF21A | Kif21a | RPMSDKVAGKVTRKLSSSESPAPDTGSSAAS | KLS(0.594)S(0.192)S(0.192)ES(0.022)PAPDTGSSAASGEADTSRPGTQQK | 4005900 |
| 925 | Q810U3 | 1230 | Neurofascin | Nfasc | KKDKEETEGNESSEATSPVNAIYSLA_____ | DKEET(0.006)EGNES(0.442)S(0.552)EAT(0.552)S(0.44)PVNAIY(0.006)S( | 0 |
| 920 | Q810U3 | 1226 | Neurofascin | Nfasc | QYTVKKDKEETEGNESSEATSPVNAIYSLA_ | DKEET(0.001)EGNES(0.831)S(0.168)EAT(0.527)S(0.473)PVNAIYSLA | 0 |
| 61 | Q01338 | 362 | Alpha-2A adrenergic receptor | Adra2a | LPRRGPGAAGPGASGSGHGEERGGGAKASRW | RGPGAAGPGAS(0.022)GS(0.978)GHGEER | 6105300 |
| 476 | Q8CHU3 | 443 | Epsin-2 | Epn2 | AESGASVPPQDSRTTSPDLFESQSLTSASSK | T(0.099)T(0.099)S(0.802)PDLFESQSLTSASSKPSSAR | 0 |
| 970 | Q9QYX7 | 1378 | Protein piccolo | Pclo | LVGEKAEKKTQPQKVSPEQPQDQQKTQTPSE | KT(0.004)QPQKVS(0.996)PEQPQDQQK | 0 |
| 824 | P27546 | 667 | Microtubule-associated protein 4 | Map4 | TGNDITTPPNKEPPPSPEKKAKPLATTQPAK | AAVGVTGNDIT(0.001)T(0.001)PPNKEPPPS(0.999)PEKK | 4692400 |
| 1299 | O35526 | 59 | Syntaxin-1A | Stx1a | IDKIAENVEEVKRKHSAILASPNPDEKTKEE | KHS(0.998)AILAS(0.002)PNPDEK | 0 |
| 729 | Q9QYR6 | 527 | crotubule-associated protein 1A;MAP1A heavy chain;MAP1 light chain L | Map1a | APPPAAGGHRELALSSPEDLTQDFEELKREE | ELALS(1)S(1)PEDLTQDFEELKR | 0 |
| 728 | Q9QYR6 | 526 | crotubule-associated protein 1A;MAP1A heavy chain;MAP1 light chain L | Map1a | AAPPPAAGGHRELALSSPEDLTQDFEELKRE | ELALS(1)S(1)PEDLTQDFEELKR | 0 |
| 609 | Q640R3 | 385 | Hepatocyte cell adhesion molecule | Hepacam | SPATGRTHTSPPRAPSSPGRSRSSSRSLRTA | THT(0.069)S(0.931)PPRAPS(0.747)S(0.253)PGR | 37604000 |
| 613 | P58406 | 332 | Histamine H3 receptor | Hrh3 | PRSLKRGSKPSASSASLEKRMKMVSQSITQR | GSKPS(0.001)AS(0.007)S(0.032)AS(0.96)LEKR | 3827600 |
| 1069 | Q8R361 | 486 | Rab11 family-interacting protein 5 | Rab11fip5 | RRGSVGEKGSPSLGASPHHSSTGEEKAKSSW | GS(0.008)PS(0.045)LGAS(0.913)PHHS(0.011)S(0.011)T(0.011)GEEK | 4676500 |
| 1336 | O88935 | 62 | Synapsin-1 | Syn1 | ATASAERASTAAPVASPAAPSPGSSGGGGFF | AS(0.014)T(0.014)AAPVAS(0.972)PAAPSPGSSGGGGFFSSLSNAVK | 0 |
| 1086 | O54829 | 245 | Regulator of G-protein signaling 7 | Rgs7 | VHGLQNDIRSHSPTHTPTPETKPPTEDELHQ | S(0.143)HS(0.845)PT(0.103)HT(0.891)PT(0.014)PET(0.003)KPPTEDELH | 13216000 |
| 619 | B5TVM2 | 594 | Immunoglobulin-like domain-containing receptor 2 | Ildr2 | SYIAWSPPTTYKAGASEGEDEDDAADEDALP | AGAS(1)EGEDEDDAADEDALPPYSELELSR | 0 |
| 1004 | Q61469 | 274 | Lipid phosphate phosphohydrolase 1 | Ppap2a | RKEEDPHTTLHETASSRNYSTNHEP______ | KEEDPHTTLHET(0.085)AS(0.393)S(0.522)R | 5002200 |
| 555 | P63216 | 5 | Guanine nucleotide-binding protein G(I)/G(S)/G(O) subunit gamma-3 | Gng3 | __________MKGETPVNSTMSIGQARKMV | MKGET(1)PVNSTMSIGQAR | 8215500 |
| 525 | Q9D2N8 | 615 | Polypeptide N-acetylgalactosaminyltransferase 15 | Galnt15 | GKCMEAGVQPSNKDLYLRQCDGKTSQLWRFD | DLY(1)LRQCDGKTSQLWR | 69392000 |
| 161 | Q8VDN2 | 16 | Sodium/potassium-transporting ATPase subunit alpha-1 | Atp1a1 | MGKGVGRDKYEPAAVSEHGDKKGKKAKKERD | DKYEPAAVS(1)EHGDKK | 4091600 |
| 743 | Q9QYR6 | 1789 | crotubule-associated protein 1A;MAP1A heavy chain;MAP1 light chain L | Map1a | PEMTGQRVPSAPGQESPVPDTKSTPPTRNEP | VPSAPGQES(1)PVPDTK | 0 |
| 1245 | P70441 | 275 | Na(+)/H(+) exchange regulatory cofactor NHE-RF1 | Slc9a3r1 | QKESSREALVEPASESPRPALARSASSDTSE | EALVEPAS(0.036)ES(0.964)PRPALAR | 2434100 |
| 1244 | P70441 | 273 | Na(+)/H(+) exchange regulatory cofactor NHE-RF1 | Slc9a3r1 | EIQKESSREALVEPASESPRPALARSASSDT | EALVEPAS(0.5)ES(0.5)PRPALAR | 2434100 |
| 806 | P20357 | 1743 | Microtubule-associated protein 2 | Map2 | SVKLDFKEKAQAKVGSLDNAHHVPGGGNVKI | VGS(1)LDNAHHVPGGGNVK | 5185300 |
| 1107 | Q8K0T0 | 92 | Reticulon-1 | Rtn1 | TASTGMAAVPDALDHSPSSTLKDGEGACYTS | SPPVAMET(0.002)AS(0.007)T(0.007)GMAAVPDALDHS(0.686)PS(0.181) | 0 |
| 990 | O70161 | 555 | Phosphatidylinositol 4-phosphate 5-kinase type-1 gamma | Pip5k1c | SDTSEQPRYRRRTQSSGQDGRPQEEPHAEDL | T(0.036)QS(0.449)S(0.516)GQDGRPQEEPHAEDLQK | 0 |
| 1318 | Q8K400 | 903 | Syntaxin-binding protein 5 | Stxbp5 | KDEKEKLKKRRPVSVSPSSSQEISENQYAVI | RRPVS(0.156)VS(0.718)PS(0.041)S(0.041)S(0.041)QEIS(0.002)ENQYAVI | 5765500 |
| 1326 | Q5DQR4 | 792 | Syntaxin-binding protein 5-like | Stxbp5l | VNRWGPGRPPFRKAQSAACMEISLPVTTEET | KAQS(0.996)AACMEIS(0.004)LPVTTEETR | 0 |
| 1204 | Q8K2P7 | 52 | Sodium-coupled neutral amino acid transporter 1 | Slc38a1 | GQINSKFISDRESRRSLTNSHLEKRKCDEYI | S(0.953)LT(0.047)NSHLEK | 5177400 |
| 59 | Q9QYB5 | 681 | Gamma-adducin | Add3 | ERTEEVLSPDGSPSKSPSKKKKKFRTPSFLK | TEEVLSPDGS(0.005)PS(0.138)KS(0.759)PS(0.098)K | 0 |
| 1260 | P70663 | 411 | SPARC-like protein 1 | Sparcl1 | KAESSPNAEPSDEGNSREHSAGSCTNFQCKR | KAESSPNAEPS(0.001)DEGNS(0.799)REHS(0.195)AGS(0.003)CT(0.002)N | 0 |
| 1298 | Q9WUD1 | 15 | STIP1 homology and U box-containing protein 1 | Stub1 | _MKGKEEKEGGARLGTGGGGSPDKSPSAQEL | LGT(0.974)GGGGS(0.026)PDKSPSAQELK | 0 |
| 1444 | Q8K207 | 95 | Uncharacterized protein C1orf21 homolog | | PGLVHQPRANMHISESQQEFFRMLDEKIEKG | ANMHIS(0.005)ES(0.995)QQEFFR | 1987700 |
| 1406 | Q7TQD2 | 19 | Tubulin polymerization-promoting protein | Tppp | SKAKPAKAANKTPPKSPGDPARAAKRLSLES | AANKT(1)PPKS(1)PGDPAR | 6746500 |
| 298 | Q3UHL1 | 470 | CaM kinase-like vesicle-associated protein | Camkv | PATQSSALPAAKAATPEPAVAQPDSTALEG | AAAT(0.998)PEPAVAQPDS(0.001)T(0.001)ALEGATGQAPPSSKGEEATGCA | 5304800 |
| 1307 | P61264 | 10 | Syntaxin-1B | Stx1b | _____MKDRTQELRSAKDSDDEEEVVHVDR | S(0.645)AKDS(0.355)DDEEEVVHVDR | 8663300 |
| 337 | Q9JKC6 | 89 | Cell cycle exit and neuronal differentiation protein 1 | Cend1 | NHSNLKPAPTVPAAPSSPDATSEPKGPGDGA | ADPVLLNNHSNLKPAPTVPAAPS(0.562)S(0.433)PDAT(0.002)S(0.002)EPK | 18313000 |
| 529 | P06837 | 86 | Neuromodulin | Gap43 | DAPVADGVEKKEGDGSATTDAAPATSPKAEE | EGDGS(0.999)ATTDAAPATSPKAEEPSK | 0 |
| 378 | O35927 | 264 | Catenin delta-2 | Ctnnd2 | LYSSSTLPAPPRGGSPLTTTQGGSPTKLQR | GGS(1)PLTT(0.004)T(0.041)QGGS(0.945)PT(0.01)K | 5493200 |
| 608 | Q640R3 | 379 | Hepatocyte cell adhesion molecule | Hepacam | PRSPARSPATGRTHTSPPRAPSSPGRSRSSS | THT(0.025)S(0.975)PPRAPS(0.288)S(0.712)PGR | 8156800 |
| 578 | Q3UNH4 | 468 | G protein-regulated inducer of neurite outgrowth 1 | Gprin1 | EVTEGKGDPQPWKKASLPASEKTDPLASSKA | KAS(0.969)LPAS(0.031)EKTDPLASSK | 8694800 |
| 946 | Q9Z0P4 | 157 | Paralemmin-1 | Palm | PLGTPKENRTSTPVRSPGGSTMMKAAMYSVE | T(0.272)S(0.364)T(0.364)PVRS(1)PGGSTMMK | 3115900 |
| 586 | Q3UNH4 | 220 | G protein-regulated inducer of neurite outgrowth 1 | Gprin1 | KTENVMHSRRERPGSTGEGDLVSLRENDMKP | ERPGS(0.439)T(0.561)GEGDLVSLR | 5750200 |
| 794 | P20357 | 608 | Microtubule-associated protein 2 | Map2 | LSDSRGSAQESLDTISPKNQHDEKELQAKAS | GS(0.008)AQES(0.987)LDT(0.01)IS(0.995)PK | 4089100 |
| 868 | P04370 | 144 | Myelin basic protein | Mbp | GGLDVMASQKRPSQRSKYLATASTMDHARHG | S(0.996)KY(0.003)LAT(0.002)ASTMDHAR | 7700800 |
| 42 | P51830 | 608 | Adenylate cyclase type 9 | Adcy9 | VIDDSRESSGPRGQGTASPGSVSDLAQTVKT | GQGT(0.973)AS(0.037)PGS(0.93)VS(0.06)DLAQTVK | 0 |
| 1027 | E9PUL5 | 255 | Proline-rich transmembrane protein 2 | Prrt2 | GHPGSPRGSLSRHPSSQLAGPGVEGGEGTQK | HPS(0.5)S(0.5)QLAGPGVEGGEGTQKPR | 8904300 |
| 1026 | E9PUL5 | 254 | Proline-rich transmembrane protein 2 | Prrt2 | GGHPGSPRGSLSRHPSSQLAGPGVEGGEGTQ | HPS(0.831)S(0.169)QLAGPGVEGGEGTQKPR | 8904300 |
| 611 | Q640R3 | 378 | Hepatocyte cell adhesion molecule | Hepacam | YPRSPARSPATGRTHTSPPRAPSSPGRSRSS | THT(0.875)S(0.123)PPRAPS(0.001)S(0.001)PGR | 56114000 |
| 1012 | P68181 | 339 | cAMP-dependent protein kinase catalytic subunit beta | Prkacb | DTSNFDDYEEEEIRVSITEKCGKEFCEF___ | GSGDTSNFDDYEEEEIRVS(0.975)IT(0.025)EK | 5469800 |
| 315 | Q6VGS5 | 905 | Protein Daple | Ccdc88c | NRDLTKQVTMHTRTLTTLREDLVLEKLSQQ | QVT(0.014)MHT(0.627)RT(0.627)LT(0.627)T(0.105)LR | 0 |
| 314 | Q6VGS5 | 903 | Protein Daple | Ccdc88c | KDNRDLTKQVTMHTRTLTTLREDLVLEKLSS | QVT(0.014)MHT(0.627)RT(0.627)LT(0.627)T(0.105)LR | 0 |
| 313 | Q6VGS5 | 901 | Protein Daple | Ccdc88c | LEKDNRDLTKQVTMHTRTLTTLREDLVLEKL | QVT(0.014)MHT(0.627)RT(0.627)LT(0.627)T(0.105)LR | 0 |
| 223 | P59017 | 387 | Bcl-2-like protein 13 | Bcl2l13 | GAAQLSEERAGSRKKSHTGEAAAVRGAKSGL | KKS(1)HTGEAAAVR | 62347000 |
| 188 | Q6Q477 | 329 | Calcium-transporting ATPase | Atp2b4 | AKTQDGVALEIQPLNSQEGLDSEDKEKKIAR | TQDGVALEIQPLNS(0.999)QEGLDS(0.001)EDKEK | 7818100 |
| 922 | Q810U3 | 1231 | Neurofascin | Nfasc | KDKEETEGNESSEATSPVNAIYSLA______ | DKEETEGNES(0.015)S(0.015)EAT(0.11)S(0.582)PVNAIY(0.128)S(0.149)L | 4787800 |
| 1085 | O54829 | 243 | Regulator of G-protein signaling 7 | Rgs7 | KSVHGLQNDIRSHSPTHTPTPETKPPTEDEL | S(0.126)HS(0.858)PT(0.609)HT(0.39)PT(0.016)PET(0.001)KPPTEDELHQ | 11837000 |
| 1251 | Q61548 | 600 | Clathrin coat assembly protein AP180 | Snap91 | DLFGTDAFSSPPRGASPVPESSLTADLLSVD | GAS(0.962)PVPES(0.011)S(0.011)LT(0.011)ADLLS(0.004)VDAFAAPSPAS | 0 |
| 1141 | B1AWN6 | 497 | Sodium channel protein | Scn2a1 | VFSESSSVASKLSSKSEKELKNRRKKKKQKE | LSS(0.003)KS(0.997)EKELK | 10988000 |
| 427 | Q6IL44 | 290 | Protein Dos | Dos | SGTVLQFFTRLRRHASLDGASPYFKVKKWKL | HAS(1)LDGASPYFK | 3842100 |
| 745 | Q9QYR6 | 504 | crotubule-associated protein 1A;MAP1A heavy chain;MAP1 light chain L | Map1a | GRAARGEKELSSEPRTPPAQKGAAPPPAASG | ELSS(0.005)EPRT(0.995)PPAQK | 10224000 |
| 786 | P20357 | 140 | Microtubule-associated protein 2 | Map2 | LAAEETANLPPSPPPSPASEQTATVEEDLLT | DQPAALPLAAEET(0.01)ANLPPS(0.989)PPPS(0.982)PAS(0.018)EQT(0.00 | 0 |
| 838 | P10637 | 708 | Microtubule-associated protein tau | Mapt | GDTSPRHLSNVSSTGSIDMVDSPQLATLADE | HLSNVS(0.001)S(0.005)T(0.026)GS(0.968)IDMVDSPQLATLADEVSASLAK | 4092500 |
| 930 | Q9CR00 | 123 | OCIA domain-containing protein 1 | Ociad1 | SPLGEALRSGELRRSSPPGHYTQKPKFDSNV | RS(0.103)S(0.897)PPGHYTQKPK | 12140000 |
| 96 | Q8C8R3 | 1793 | Ankyrin-2 | Ank2 | PSSKSSTSSKAERHSSLSSSAKPERHTPSVP | HS(0.002)S(0.992)LS(0.004)S(0.001)S(0.001)AKPER | 5931600 |
| 872 | P04370 | 229 | Myelin basic protein | Mbp | ENPVVHFFKNIVTPRTPPPSQGKGGRDSRSG | NIVT(0.01)PRT(0.98)PPPS(0.01)QGK | 6883500 |
| 818 | P20357 | 606 | Microtubule-associated protein 2 | Map2 | YELSDSRGSAQESLDTISPKNQHDEKELQAK | GSAQESLDT(0.748)S(0.251)PK | 0 |
| 359 | Q71M36 | 541 | Chondroitin sulfate proteoglycan 5 | Cspg5 | LKSRLKEEESFNIQNSMSPKLEGGKGDQDDL | LKEEESFNIQNS(0.659)MS(0.341)PK | 0 |
| 275 | O55017 | 895 | Voltage-dependent N-type calcium channel subunit alpha-1B | Cacna1b | AREERARPRRSHSKETPGADTQVRCERSRRH | RS(0.078)HS(0.274)KET(0.648)PGADTQVR | 6991800 |
| 460 | Q9WV92 | 470 | Band 4.1-like protein 3;Band 4.1-like protein 3, N-terminally processed | Epb41l3 | ASVSENHDIYMKDSVSAAEVGTGQYATTKGI | DS(0.152)VS(0.848)AAEVGTGQYATTK | 0 |
| 1418 | D3Z7M3 | 126 | | Vmn2r73 | KTGLGLKHEEFIPNYYCANERRYLIVLTAPI | HEEFIPNY(0.004)Y(0.996)CANERR | 0 |
| 94 | Q8C8R3 | 1925 | Ankyrin-2 | Ank2 | APGSATGKHEKHLPVSPGKTEKQPPISPTSK | HLPVS(0.996)PGKT(0.004)EK | 7265200 |
| 847 | P10637 | 695 | Microtubule-associated protein tau | Mapt | HGAEIVYKSPVVSGDTSPRHLSNVSSTGSID | TDHGAEIVY(0.052)KS(0.893)PVVS(0.599)GDT(0.771)S(0.685)PR | 0 |
| 672 | Q5DTY9 | 130 | BTB/POZ domain-containing protein KCTD16 | Kctd16 | LPDLVKLLAPEDVKQSPDEFCHSDFEDASQG | LLAPEDVKQS(1)PDEFCHS(1)DFEDASQGSDTR | 3428500 |
| 1072 | Q8BIV3 | 696 | Ran-binding protein 6 | Ranbp6 | SGLEAKATACQMLVYAKELREGFVEYTEQV | AT(0.719)ACQMLVY(0.64)Y(0.64)AKELR | 0 |
| 1071 | Q8BIV3 | 695 | Ran-binding protein 6 | Ranbp6 | TSGLEAKATACQMLVYAKELREGFVEYTEQ | AT(0.719)ACQMLVY(0.64)Y(0.64)AKELR | 0 |
| 1070 | Q8BIV3 | 688 | Ran-binding protein 6 | Ranbp6 | QQSFGIKTSGLEAKATACQMLVYAKELREG | AT(0.719)ACQMLVY(0.64)Y(0.64)AKELR | 0 |
| 214 | Q91XV3 | 31 | Brain acid soluble protein 1 | Basp1 | DKAKDKDKKAEGAGTEEEGTPKESEPQAAA | AEGAGT(1)EEEGT(1)PKESEPQAAADATEVK | 0 |
| 948 | Q9Z0P4 | 335 | Paralemmin-1 | Palm | QDTIKAELVVIEDAATPREPAPLNGSAAELP | AELVVIEDAAT(1)PR | 0 |
| 1344 | O88935 | 438 | Synapsin-1 | Syn1 | QRDASPGRGSHSQSSSPGALTLGRQTSQQPA | GSHSQS(0.008)S(0.039)S(0.953)PGALTLGR | 15382000 |
| 170 | G5E829 | 1140 | Plasma membrane calcium-transporting ATPase 1 | Atp2b1 | RSSLYEGLEKPESRSSIHNFMTHPEFRIEDS | S(0.5)S(0.5)IHNFMTHPEFR | 3347000 |
| 169 | G5E829 | 1139 | Plasma membrane calcium-transporting ATPase 1 | Atp2b1 | FRSSLYEGLEKPESRSSIHNFMTHPEFRIED | S(0.5)S(0.5)IHNFMTHPEFR | 3347000 |
| 1349 | O88935 | 551 | Synapsin-1 | Syn1 | VAGGPGAPPAARPPASPSPQRQAGAPQATRQ | PVAGGPGAPPAARPPAS(1)PS(1)PQR | 10210000 |
| 550 | A2APB9 | 1077 | | Gm14025 | VQKDLLVSLESILTLTKEVVRKVPVLQEVQS | DLLVS(0.992)LES(0.352)ILT(0.828)LT(0.828)K | 0 |
| 549 | A2APB9 | 1075 | | Gm14025 | ETVQKDLLVSLESILTLTKEVVRKVPVLQEV | DLLVS(0.992)LES(0.352)ILT(0.828)LT(0.828)K | 0 |
| 548 | A2APB9 | 1069 | | Gm14025 | AGGQESETVQKDLLVSLESILTLTKEVVRKV | DLLVS(0.992)LES(0.352)ILT(0.828)LT(0.828)K | 0 |
| 386 | O35927 | 269 | Catenin delta-2 | Ctnnd2 | STLPAPPRGGSPLTTTQGGSPTKLQRGGSAP | GGSPLT(0.004)T(0.018)T(0.614)QGGS(0.3)PT(0.064)K | 8802700 |
| 807 | P20357 | 938 | Microtubule-associated protein 2 | Map2 | AAGRVKDEFTAEKEASPPTSADKSRLSREFD | VKDEFT(0.001)AEKEAS(0.996)PPT(0.002)S(0.001)ADK | 7587900 |
| 449 | Q9Z2H5 | 546 | Band 4.1-like protein 1 | Epb41l1 | DWDRERRLPSSPASPSPKGTPEKASERAGLR | RLPSSPAS(0.001)PS(0.999)PK | 7618500 |
| 635 | Q8K0T4 | 471 | Katanin p60 ATPase-containing subunit A-like 1 | Katnal1 | RGDLELALKKIAKSVSAADLEKYEKWMVEFG | KIAKS(0.5)VS(0.5)AADLEK | 5577800 |
| 634 | Q8K0T4 | 469 | Katanin p60 ATPase-containing subunit A-like 1 | Katnal1 | VTRGDLELALKKIAKSVSAADLEKYEKWMVE | KIAKS(0.5)VS(0.5)AADLEK | 5577800 |
| 1228 | P31650 | 23 | Sodium- and chloride-dependent GABA transporter 3 | Slc6a11 | PLGNGKAAEEARGSETLGGGGGGAAGTREAR | AAEEARGS(0.5)ET(0.5)LGGGGGGAAGTR | 8968800 |
| 673 | Q5DTY9 | 137 | BTB/POZ domain-containing protein KCTD16 | Kctd16 | LAPEDVKQSPDEFCHSDFEDASQGSDTRICP | LLAPEDVKQS(1)PDEFCHS(1)DFEDASQGSDTR | 0 |
| 1106 | Q8K0T0 | 352 | Reticulon-1 | Rtn1 | GTEPSAAESQGKGSVSEDELIAAIKEAKGLS | GS(0.033)VS(0.967)EDELIAAIK | 1768200 |
| 1119 | Q99P72 | 838 | Reticulon-4 | Rtn4 | SSKEDKMKESETFSDSSPIEIIDEFPTFVSA | MKES(0.019)ET(0.05)FS(0.523)DS(0.714)S(0.694)PIEIIDEFPTFVSAK | 0 |
| 1118 | Q99P72 | 836 | Reticulon-4 | Rtn4 | LLSSKEDKMKESETFSDSSPIEIIDEFPTFV | MKES(0.015)ET(0.053)FS(0.863)DS(0.565)S(0.504)PIEIIDEFPTFVSAK | 0 |
| 409 | Q9JLM8 | 336 | Serine/threonine-protein kinase DCLK1 | Dclk1 | QLSTPRSGKSPSPSPTSPGSLRKQRISQHGG | SPSPSPT(0.929)S(0.069)PGS(0.002)LRK | 9907800 |
| 426 | P60904 | 11 | DnaJ homolog subfamily C member 5 | Dnajc5 | _____MADQRQRSLSTSGESLYHVLGLDKNA | SLS(0.414)T(0.577)S(0.009)GESLYHVLGLDK | 57103000 |

| | | | | | | | |
|---|---|---|---|---|---|---|---|
| 0 | 0,170138076 | 0,100501402 | 0,123669267 | -3,248732208 | 0,001381644 | -1,405155625 | 0,390812678 |
| 0 | 0,080562077 | 0,076170769 | 0,124066782 | -3,125253466 | 0,001450507 | -1,30453481 | 0,339710347 |
| 0 | 0,081658396 | 0,056041385 | 0 | -2,215937885 | 0,001554914 | -1,214811022 | 0,249662758 |
| 0,057994636 | 0,180347983 | 0,210597614 | 0,181733128 | -2,16316876 | 0,002046383 | -0,473983601 | 0,534202845 |
| 0,029467464 | 0,962234482 | 0,61695938 | 0,563501925 | -2,072989142 | 0,003160218 | -2,590478385 | 0,115580256 |
| 0,047172043 | 0,210514144 | 0,098210665 | 0,147431668 | -1,058365037 | 0,003565093 | -0,685028417 | 0,373384345 |
| 0,028368844 | 0,079441421 | 0,08761225 | 0,087357758 | -2,230844241 | 0,003622206 | -0,512743649 | 0,481910609 |
| 0,023885525 | 0 | 0,066226716 | 0,056038361 | -1,492135685 | 0,003935742 | -0,261815733 | 0,657934457 |
| 0 | 0,294406899 | 0,502122093 | 0,465598083 | -3,952741045 | 0,004434669 | -4,01231601 | 0,056765417 |
| 0 | 0,294406899 | 0,502122093 | 0,465598083 | -3,952741045 | 0,004434669 | -4,01231601 | 0,056765417 |
| 0,117470518 | 0,363500503 | 0 | 0,148105898 | -2,24305933 | 0,004679979 | 0,205369528 | 0,874527887 |
| 0,070276484 | 0,189943673 | 0 | 0,083299735 | -2,910910033 | 0,004803756 | 0,976124223 | 0,401974213 |
| 0,081994509 | 0,578638657 | 0,132168028 | 0,143639123 | -2,71549682 | 0,004807962 | -0,534803739 | 0,567946771 |
| 0,070367083 | 0,215138154 | 0,192187222 | 0,12313892 | -1,926826979 | 0,005098311 | -0,416953888 | 0,454888334 |
| 0,356325771 | 1,717373575 | 2,395134777 | 1,926191365 | -1,950943012 | 0,005419431 | -1,133982079 | 0,221999796 |
| 0,03156355 | 0,270753754 | 0,195360456 | 0,270857931 | -1,802954742 | 0,005447205 | -1,544407293 | 0,104830278 |
| 0,055990908 | 0,240019732 | 0,228097345 | 0,230010819 | -1,446985068 | 0,0059439 | -0,863963536 | 0,238149438 |
| 0 | 0,075619369 | 0,042541056 | 0,092886445 | -2,189135576 | 0,006109273 | -1,410149329 | 0,253039326 |
| 0,040575926 | 0,023319381 | 0,050000973 | 0,064226044 | -2,446754313 | 0,006184682 | -0,439010414 | 0,533696946 |
| 0 | 0,048451282 | 0 | 0,106080287 | -2,297445359 | 0,00622877 | -1,531607399 | 0,18851432 |
| 0 | 0 | 0 | 0 | -1,113189235 | 0,006402421 | -0,709904585 | 0,381215537 |
| 0,021269596 | 0,167970933 | 0 | 0,170032425 | -2,054608938 | 0,006560264 | -2,10709001 | 0,099642295 |
| 0,018103918 | 0,074908964 | 0,051577263 | 0,046211456 | -2,040035694 | 0,006779659 | -0,964152175 | 0,330586293 |
| 0,014830928 | 0 | 0,143490279 | 0,141799036 | -1,703978147 | 0,007348442 | -1,281097629 | 0,3159593 |
| 0,051344864 | 0,148744786 | 0,098830291 | 0,141265271 | -1,835716532 | 0,00747354 | -0,564210133 | 0,248524946 |
| 0,026388866 | 0 | 0,086012489 | 0,097350411 | -2,376844334 | 0,007623392 | -1,517388857 | 0,073530267 |
| 0,048022616 | 0 | 0,136909103 | 0,152109139 | -3,385380142 | 0,007726124 | -0,914443936 | 0,347911028 |
| 0,012882615 | 0 | 0,026064647 | 0 | -1,145943757 | 0,007825822 | -0,726858607 | 0,104652449 |
| 0,012882615 | 0 | 0,026064647 | 0 | -1,145943757 | 0,007825822 | -0,726858607 | 0,104652449 |
| 0 | 0 | 0,048365537 | 0,047426475 | -1,797998704 | 0,007895173 | -0,738337294 | 0,46023936 |
| 0,061868392 | 0,19957413 | 0,29092363 | 0,183292286 | -1,831268084 | 0,008145302 | -0,888310583 | 0,175642922 |
| 0,067761709 | 0 | 0,130572023 | 0,191607791 | -1,161789796 | 0,008681868 | -0,38030737 | 0,754628621 |
| 0,067761709 | 0,303353605 | 0,130572023 | 0,191607791 | -1,161789796 | 0,008681868 | -1,506158179 | 0,152346023 |
| 0,010444365 | 0 | 0,048125197 | 0 | -1,778357006 | 0,008718827 | -0,980743437 | 0,104784733 |
| 0,392837947 | 0,867668261 | 0,4971557 | 0,410044325 | -1,689956005 | 0,008784546 | -0,097061572 | 0,822545439 |
| 0 | 0,208752616 | 0,139406382 | 0,154960571 | -1,386971618 | 0,00896569 | -1,624052691 | 0,310691466 |
| 0,047412174 | 0,155709774 | 0,094690065 | 0,106542416 | -2,960376702 | 0,008977899 | -0,71358743 | 0,262487152 |
| 0 | 0,081241192 | 0,096943249 | 0,098780341 | -1,777807486 | 0,009237439 | -1,69201124 | 0,155514498 |
| 0 | 0 | 0,041573126 | 0,035145653 | -1,64724554 | 0,009524588 | -1,099564149 | 0,123800888 |
| 0 | 0,082608694 | 0,062652602 | 0 | -1,408518184 | 0,009605078 | -1,357643254 | 0,243193031 |
| 0 | 0 | 0 | 0 | -1,505547586 | 0,010065591 | -0,751414391 | 0,333631301 |
| 0,271857828 | 0,396065587 | 0,249080122 | 0,391755832 | -1,194897716 | 0,010390054 | -0,461885052 | 0,182844843 |
| 0 | 0,258191281 | 0,32077832 | 0,257724489 | -1,545333956 | 0,010634002 | -1,766509885 | 0,361191123 |
| 0,033852268 | 0,211476031 | 0,295918188 | 0,302265821 | -2,664249571 | 0,011103957 | -1,567848234 | 0,116648081 |
| 0,274813282 | 1,785864552 | 0,158398847 | 0,709725597 | -1,371457118 | 0,011205524 | 0,210539528 | 0,87573984 |
| 0 | 0 | 0,043237665 | 0,029809402 | -1,482438561 | 0,01206559 | -1,202849578 | 0,09627331 |
| 0 | 0 | 0,105599232 | 0 | -2,066701856 | 0,012341876 | -0,83548085 | 0,589259459 |
| 0,059485557 | 0,146531287 | 0,192346822 | 0,190455981 | -2,284460577 | 0,012535909 | -0,74388006 | 0,164793448 |
| 0,016349996 | 0,037847349 | 0,046232523 | 0 | -1,493629336 | 0,012577441 | -0,453187181 | 0,391123226 |
| 0 | 0,04364996 | 0 | 0,063734418 | -1,359927056 | 0,013074938 | -1,402927635 | 0,122315042 |
| 0,864954201 | 1,048549337 | 0,627887323 | 0 | -1,242377711 | 0,013188263 | 1,648653303 | 0,360408399 |
| 0 | 0,025850418 | 0 | 0,04890978 | -2,112221168 | 0,013389046 | -1,089541859 | 0,153819743 |
| 0 | 0,075357458 | 0,069495711 | 0 | -1,360014196 | 0,013474819 | -0,804302925 | 0,505538689 |
| 0,01658309 | 0 | 0,029351479 | 0 | -1,229846816 | 0,013583324 | -0,708817896 | 0,089268894 |
| 0,040804622 | 0 | 0,093272437 | 0 | -1,360335996 | 0,013672853 | -0,018734916 | 0,987048742 |
| 0 | 0,040938135 | 0 | 0 | -1,877495495 | 0,013676075 | -1,08043487 | 0,13231446 |
| 0 | 0 | 0,032288129 | 0 | -1,38108517 | 0,01380997 | -1,392169218 | 0,08860223 |
| [illegible] | [illegible] | [illegible] | [illegible] | [illegible] | [illegible] | [illegible] | [illegible] |

| | | | | | | | | |
|---|---|---|---|---|---|---|---|---|
| 0,03174874 | 0,023173929 | 0,074494077 | 0,109748846 | 0,090228855 | -2,305231224 | 0,025724311 | -1,147426492 | 0,139901317 |
| 0,014704006 | 0,021398897 | 0,042755289 | 0,101027145 | 0 | -1,957820667 | 0,026095354 | -0,498743888 | 0,687543578 |
| 0,036985922 | 0,018299189 | 0,074310971 | 0 | 0,083985202 | -1,573063451 | 0,026167226 | -0,524739544 | 0,557235289 |
| 0 | 0 | 0 | 0,049284648 | 0,039220532 | -2,428367514 | 0,02672055 | -1,397746704 | 0,069577721 |
| 0,054507523 | 0,035837524 | 0,107792745 | 0,097206121 | 0,154637503 | -1,295267209 | 0,026726165 | -0,966458352 | 0,13558866 |
| 0,082849542 | 0 | 0,144827705 | 0,064157542 | 0,028512914 | -1,535232365 | 0,02681163 | -0,525473927 | 0,734169349 |
| 0,125451172 | 0,067065066 | 0,164053852 | 0,115597736 | 0,151856303 | -1,650049977 | 0,026861182 | -0,55663221 | 0,149013248 |
| 0,125451172 | 0,067065066 | 0,164053852 | 0,115597736 | 0,151856303 | -1,650049977 | 0,026861182 | -0,55663221 | 0,149013248 |
| 0,099538946 | 0,040400007 | 0,121846722 | 0,087896715 | 0,090136148 | -1,470304465 | 0,02709359 | -0,593823653 | 0,218823205 |
| 0,107879473 | 0,048636577 | 0 | 0 | 0,01615905 | -3,003600148 | 0,027336958 | 1,960168464 | 0,016114188 |
| 0 | 0 | 0 | 0 | 0 | -1,649778453 | 0,027457406 | -0,300479861 | 0,80022284 |
| 0 | 0,021045298 | 0,13666905 | 0,151169882 | 0,113741228 | -2,927019816 | 0,028021141 | -1,802664512 | 0,191963553 |
| 0,073471784 | 0,03272638 | 0,052685902 | 0,134552648 | 0,18712697 | -1,947496213 | 0,028148938 | -0,612954869 | 0,509226867 |
| 0 | 0,472600033 | 0,719827417 | 1,001164834 | 0,689273948 | -1,852088984 | 0,028263281 | -1,745115609 | 0,454863287 |
| 0,180903151 | 0,088936309 | 0,447509148 | 0,211348675 | 0,219490019 | -1,004993388 | 0,028758533 | -0,580355612 | 0,45696043 |
| 0,055329515 | 0 | 0,107042937 | 0 | 0,109400871 | -1,776606535 | 0,028780895 | -0,850946049 | 0,564008472 |
| 0,047479771 | 0,033788057 | 0,20280746 | 0,017123261 | 0,097718428 | -2,083745424 | 0,029017821 | 0,015624303 | 0,991277365 |
| 0,111933767 | 0,083146785 | 0,202332312 | 0,346229913 | 0 | -2,962561948 | 0,029130975 | 0,555683686 | 0,720358365 |
| 0,598291317 | 0,155645137 | 0,964563871 | 0,789008768 | 0,906544626 | -1,249423878 | 0,029440442 | -0,92729859 | 0,326733109 |
| 0,054552055 | 0,103212209 | 0,130480526 | 0,202664528 | 0,075675315 | -1,428353489 | 0,029510929 | -0,729246088 | 0,210687532 |
| 0,160705902 | 0,017014973 | 0,053333726 | 0,166116003 | 0,101262351 | -2,189081725 | 0,029553627 | -0,589002247 | 0,617352014 |
| 0,22186358 | 0 | 0,202042587 | 0,231786934 | 0,276546748 | -2,437761392 | 0,029828448 | -1,700230121 | 0,350537427 |
| 0 | 0 | 0 | 0 | 0 | -1,682936716 | 0,030314026 | -0,917745118 | 0,162201914 |
| 0,038538986 | 0,022311922 | 0,095889685 | 0 | 0,067597195 | -3,075871935 | 0,031190279 | -0,896210317 | 0,240939424 |
| 0,022011012 | 0 | 0,053178434 | 0,043509925 | 0,046858998 | -1,586127302 | 0,031196425 | -1,176459495 | 0,222248781 |
| 0,038266226 | 0 | 0,057008598 | 0 | 0,063622046 | -1,445901972 | 0,031320971 | -0,614584023 | 0,602147498 |
| 0,077463919 | 0 | 0 | 0,097647369 | 0 | -2,381589442 | 0,0315798 | -0,792111335 | 0,516598982 |
| 12,21205279 | 9,691427656 | 30,68418931 | 22,40134482 | 25,01113064 | -1,125273307 | 0,031643205 | -0,889106036 | 0,086538929 |
| 0 | 0 | 0,30144142 | 0 | 0,18021611 | -2,860127556 | 0,031865264 | -2,897925601 | 0,079628381 |
| 0,072916986 | 0 | 0 | 0 | 0 | -2,123941858 | 0,031876233 | 0,510789432 | 0,68714615 |
| 0 | 0 | 0 | 0 | 0 | -1,700256836 | 0,03188067 | -0,917745118 | 0,162201914 |
| 0 | 0 | 0,103095724 | 0 | 0 | -2,866186886 | 0,032108743 | -1,524590607 | 0,144241426 |
| 0 | 0,080102491 | 0 | 0 | 0 | -2,870891382 | 0,032416273 | 1,101218638 | 0,36493752 |
| 0,476940826 | 0,19146243 | 1,036798094 | 0,627690169 | 0,476146977 | -1,266026694 | 0,032812886 | -0,837851706 | 0,234134761 |
| 0,157319592 | 0,051080104 | 0,268320065 | 0,237363564 | 0,240461387 | -2,573410161 | 0,032864401 | -0,854302556 | 0,33257504 |
| [illegible] | [illegible] | [illegible] | [illegible] | [illegible] | [illegible] | [illegible] | [illegible] | [illegible] |

| | | | | | | | | |
|---|---|---|---|---|---|---|---|---|
| 0,041334315 | 0,017847954 | 0,209668147 | 0,074266828 | 0,066018372 | -1,015506829 | 0,222619288 | -1,587970879 | 0,087258778 |
| 0,425162762 | 0,421090684 | 0,879210902 | 0,525029466 | 0,58011889 | -2,003833121 | 0,222735043 | -0,445913607 | 0,185411409 |
| 0,036971078 | 0,033368488 | 0,096827235 | 0,053192984 | 0 | -1,117728829 | 0,223880502 | -0,303836106 | 0,626656202 |
| 0,257257473 | 0,204524483 | 0,765163576 | 0,558677016 | 0,786180491 | -1,694023716 | 0,22391173 | -1,201642139 | 0,052037476 |
| 0,055467751 | 0,040234642 | 0,083066459 | 0,092602114 | 0,099602059 | -1,068680394 | 0,22418285 | -0,461141908 | 0,422090467 |
| 0 | 0 | 0,042664895 | 0,041357196 | 0,06928558 | -1,226185687 | 0,224330882 | -1,769619853 | 0,037360434 |
| 0 | 0,064120167 | 0 | 0 | 0 | -1,697539899 | 0,224424293 | 1,082917105 | 0,393748595 |
| 0 | 0,033104608 | 0,082751238 | 0 | 0 | -1,677783298 | 0,229874505 | 0,13607236 | 0,913145275 |
| 0 | 0,049128273 | 0 | 0 | 0 | -1,182347215 | 0,230127076 | -0,033517282 | 0,952161215 |
| 0,113798556 | 0,080119203 | 0,260219355 | 0,102116184 | 0,042876826 | -1,182347215 | 0,230127076 | -0,041649405 | 0,95947028 |
| 0,075587068 | 0,049745752 | 0,162999253 | 0 | 0 | -1,512513836 | 0,236297389 | -0,056016767 | 0,956870646 |
| 0 | 0 | 0 | 0 | 0 | -1,373433823 | 0,240296999 | -0,368349192 | 0,74240764 |
| 0 | 0 | 0 | 0 | 0 | -1,373433823 | 0,240296999 | -0,368349192 | 0,74240764 |
| 0,263445606 | 0,15787932 | 1,053625318 | 0,310347955 | 0,420817962 | -1,149453085 | 0,241041035 | -0,46208436 | 0,681881517 |
| 0,263445606 | 0,15787932 | 1,053625318 | 0,310347955 | 0,420817962 | -1,149453085 | 0,241041035 | -0,46208436 | 0,681881517 |
| 0,617356703 | 0,456301054 | 1,626168163 | 1,105092954 | 1,291516016 | -2,511288561 | 0,242852689 | -1,069751391 | 0,029488281 |
| 0 | 0 | 0 | 0 | 0 | -1,367136976 | 0,246040411 | -0,917745118 | 0,162201914 |
| 0,102210883 | 0 | 0,199307583 | 0,172086941 | 0 | -1,206491897 | 0,251302662 | -1,012508133 | 0,563832287 |
| 0 | 0 | 0 | 0 | 0,131354087 | -1,71771676 | 0,254033079 | -1,01716881 | 0,491319704 |
| 0,196220403 | 0,089947848 | 0,292958274 | 0,104782452 | 0,287797965 | -1,722835085 | 0,254567695 | -0,38681119 | 0,577870364 |
| 0,067021097 | 0,020769984 | 0 | 0,109664351 | 0,170116704 | -1,555869911 | 0,254878966 | -1,276987761 | 0,236839512 |
| 0,113919164 | 0,122361091 | 0,430705101 | 0,234471978 | 0,266166413 | -1,54033044 | 0,25883317 | -1,083039939 | 0,044355894 |
| 6,160114571 | 1,964145757 | 13,93692868 | 3,573362334 | 4,001696624 | -2,088819953 | 0,263072741 | -1,023130792 | 0,287945807 |
| 0 | 0 | 0,227851285 | 0,068421693 | 0,198673162 | -1,595358127 | 0,263505432 | -2,586560858 | 0,125774771 |
| 0 | 0 | 0,227851285 | 0,068421693 | 0,198673162 | -1,595358127 | 0,263505432 | -2,586560858 | 0,125774771 |
| 0 | 0,280134859 | 0,278738574 | 0,279929969 | 0,244267978 | -1,47262154 | 0,264428871 | -1,172852694 | 0,486640886 |
| 0 | 0,039507213 | 0,215057031 | 0,174593609 | 0 | -1,673084918 | 0,264886395 | -1,088532346 | 0,491394791 |
| 0 | 0 | 0 | 0 | 0 | -1,375816676 | 0,267493935 | -0,917745118 | 0,162201914 |
| 0 | 0 | 0 | 0 | 0 | -1,553341537 | 0,269493138 | -0,534802205 | 0,578769883 |
| 0,039501997 | 0 | 0,157923272 | 0,078433341 | 0 | -1,46461867 | 0,277255234 | -1,822531515 | 0,139158852 |
| 0 | 0 | 0,102335486 | 0 | 0 | -1,325026046 | 0,277459694 | -1,521031276 | 0,143958739 |
| 0,206833932 | 0,164740197 | 0,451843433 | 0,349778678 | 0,530801761 | -1,592671471 | 0,282520016 | -1,164675853 | 0,005635276 |
| 0,097108225 | 0,034981673 | 0,192875689 | 0,091200446 | 0,141616432 | -1,36222368 | 0,282710793 | -0,85612788 | 0,252121054 |
| 0 | 0 | 0 | 0 | 0 | -1,365954159 | 0,285707474 | 0,04429178 | 0,977056477 |
| 0 | 0 | 0,144051242 | 0,130675294 | 0 | -1,718939102 | 0,291347557 | -1,544899773 | 0,394007584 |
| [illegible] | [illegible] | [illegible] | [illegible] | [illegible] | [illegible] | [illegible] | [illegible] | [illegible] |
